# Ultra-Conserved Elements and morphology reciprocally illuminate conflicting phylogenetic hypotheses in Chalcididae (Hymenoptera, Chalcidoidea)

**DOI:** 10.1101/761874

**Authors:** Astrid Cruaud, Gérard Delvare, Sabine Nidelet, Laure Sauné, Sujeevan Ratnasingham, Marguerite Chartois, Bonnie B. Blaimer, Michael Gates, Seán G. Brady, Sariana Faure, Simon van Noort, Jean-Pierre Rossi, Jean-Yves Rasplus

**Affiliations:** CBGP, Univ Montpellier, CIRAD, INRA, IRD, Montpellier SupAgro, Montpellier, France; CIRAD, UMR CBGP, F-34398 Montpellier, France; Centre for Biodiversity Genomics, University of Guelph, Canada; Department of Entomology, National Museum of Natural History, Smithsonian Institution, Washington, DC, USA; USDA, ARS, SEL, c/o Smithsonian Institution, National Museum of Natural History, Washington, USA; Department of Zoology and Entomology, Rhodes University, Grahamstown, South Africa; Research and Exhibitions Department, South African Museum, Iziko Museums of South Africa, P.O. Box 61, Cape Town, 8000, South Africa; Department of Biological Sciences, University of Cape Town, Private Bag, Rondebosch, 7701, Cape Town, South Africa; North Carolina State University, Raleigh, NC, USA

**Keywords:** morphology, systematic bias, topological conflict, Ultra Conserved Elements.

## Abstract

Recent technical advances combined with novel computational approaches promised the acceleration of our understanding of the tree of life. However, when it comes to hyperdiverse and poorly known groups of invertebrates, studies are still scarce. As published phylogenies will be rarely challenged by future taxonomists, careful attention must be paid to potential analytical bias. We present the first molecular phylogenetic hypothesis for the family Chalcididae, an emblematic group of parasitoid wasps, with a representative sampling (144 ingroups and 7 outgroups) that covers all described subfamilies and tribes and 82% of the known genera. Analyses of 538 Ultra-Conserved Elements (UCEs) with supermatrix (RAxML and IQTREE) and gene-tree reconciliation approaches (ASTRAL, ASTRID) resulted in highly supported topologies in overall agreement with morphology but reveal conflicting topologies for some of the deepest nodes. To resolve these conflicts, we explored the phylogenetic tree space with clustering and gene genealogy interrogation methods, analyzed marker and taxon properties that could bias inferences and performed a thorough morphological analysis (130 characters encoded for 40 taxa representative of the diversity). This joint analysis reveals that UCEs enable attainment of resolution between ancestry and convergent /divergent evolution when morphology is not informative enough, but also shows that a systematic exploration of bias with different analytical methods and a careful analysis of morphological features is required to prevent publication of artefactual results. We highlight a GC-content bias for ML approaches, an artefactual mid-point rooting of the ASTRAL tree and a deleterious effect of high percentage of missing data on gene tree reconciliation methods. Based on the results we propose a new classification of the family into eight subfamilies and 10 tribes that lay the foundation for future studies on the evolutionary history of Chalcididae.

## INTRODUCTION

At a time when biodiversity studies are of critical importance (Dirzo et al. 2014, Hallmann et al. 2017), efforts made to solve the tree of life are unequal between the different groups of organisms. In animals, most attempts using pangenomic data focus on vertebrates, for which past research has provided a solid framework based on external morphology, anatomy, biology and fossils (Titley et al. 2017). Many teams continuously add to our knowledge of vertebrate groups by performing phylogenomic studies to test previous hypotheses and resolve long-standing taxonomic disputes. When it comes to invertebrate groups, specifically to insects, which are the most speciose terrestrial organisms (Foottit and Adler, 2009), the picture is quite different. Background knowledge is poor for most groups, essentially based on a small number of morphological features, and only a few phylogenomic hypotheses using representative but limited sampling have been published at the family level. This is certainly a consequence of the so-called taxonomic impediment (Ebach et al. 2011, Wägele et al. 2011), difficulty in adapting to new technologies and the inherent complexity of working with hyperdiverse groups. Obtaining a representative taxon sampling is problematic as new species and genera are constantly discovered, whereas some of the described taxa have only been found once. In addition, sampling is complicated by the recent restrictive access regulations to reduce the risk of supposed biopiracy (Prathapan et al. 2018). Finally, in many groups, species complexes do exist that are difficult to untangle based on morphology alone. Consequently, pangenomic data must be obtained from single, often tiny specimens, which is technically challenging.

Parasitic wasps and more precisely chalcid wasps (ca 500,000 species, Heraty et al. 2013) belong to these hyperdiverse and poorly known groups. As they naturally regulate populations of others insects, chalcid wasps have a key functional role in the ecosystems (Godfray 1994) and are frequently used as biological control agents (Consoli et al. 2010). However, families of chalcid wasps are understudied and only a few family-wide, Sanger based and poorly resolved phylogenetic hypotheses with reduced sampling have been published (Chen et al. 2004, Desjardins et al. 2007, Owen et al. 2007, Burks et al. 2011, Cruaud et al. 2012, Murray et al. 2013, Janšta et al. 2017). Consequently, classifications are still based on morphological characters, although morphological convergence due to similar biology is widespread (van Noort and Compton 1996, Heraty et al. 2013).

By reducing stochastic errors, genome-scale data offer greater opportunities to better resolve phylogenetic relationships (Philippe et al. 2005). Phylogenomic trees are usually highly supported but high statistical support is still often confused with accuracy (Lartillot et al. 2007). However, as for reduced-size data sets, when the strongest signal that emerges from the data is not the historical signal, models or methods can be misled and infer incorrect topologies with high support (systematic error; Swofford et al. 1996, Phillips et al. 2004)). With the increase in the number of gene regions, the probability to observe conflicting signal between markers due to violation of model assumption also increases (Kumar et al. 2012, Philippe et al. 2017) and total evidence approaches can lead to incorrect yet highly supported trees. This is why it is crucial to interpret molecular results in the light of morphological and biological data to point out possible contradictions (Wiens 2004). However, in the case of poorly known groups, a feedback on morphological features is not straightforward.

Here, we were interested in testing to which extent a total evidence approach using pangenomic data with or without knowledge on the morphology and biology of the target group could lead to artefactual results and fill knowledge gaps. We focused on the Chalcididae (Delvare 2017), the type family of Chalcidoidea that comprises 1,548 described species and 83 genera classified into six subfamilies and seven tribes (Noyes 2018) (Table 1). The family is found on all continents except the polar regions but has its greatest diversity in the tropics. Species diversity within genera varies greatly, and four genera (*Antrocephalus*, *Brachymeria*, *Conura, Hockeria*) represent more than half (54%) of the species diversity while 65 genera (78%) comprise less than five described species. The evolutionary history of the Chalcididae has been the focus of a single study based on a limited sampling (22 taxa) and a few (34) morphological characters (Wijesekara 1997b). Another morphological study has addressed relationships within two tribes (Wijesekara 1997a). Currently, our knowledge of the infra-familial relationships comes mainly from the three analyses that have focused on the higher level classification of Chalcidoidea using 18S and 28S ribosomal gene regions (Campbell et al.(2000); 11 taxa and Munro et al. (2011); 41 taxa) or morphology plus rRNA (Heraty et al.(2013); 25 taxa). In two of the three analyses, the family was retrieved polyphyletic and infrafamilial relationships were never resolved. Although Chalcididae are understudied, they are among the best documented chalcid wasps. Their medium size (1.5 to 15mm) enables the observation of multiple morphological features. Furthermore, characters used in previous studies can serve as a starting point for a thorough morphological analysis. We chose to infer the molecular phylogeny of the family using Ultra-Conserved Elements (UCEs) and their flanking regions (Faircloth et al. 2012, McCormack et al. 2012) that are increasingly used to solve ancient and recent divergences in insects (Blaimer et al. 2015, Faircloth et al. 2015, Blaimer et al. 2016a, Bossert et al. 2017, Branstetter et al. 2017a, Branstetter et al. 2017b, Jesovnik et al. 2017, Prebus 2017, Van Dam et al. 2017, Ward and Branstetter 2017, Bossert et al. 2019, Cruaud et al. 2019, Kieran et al. 2019).

**Table 1.**
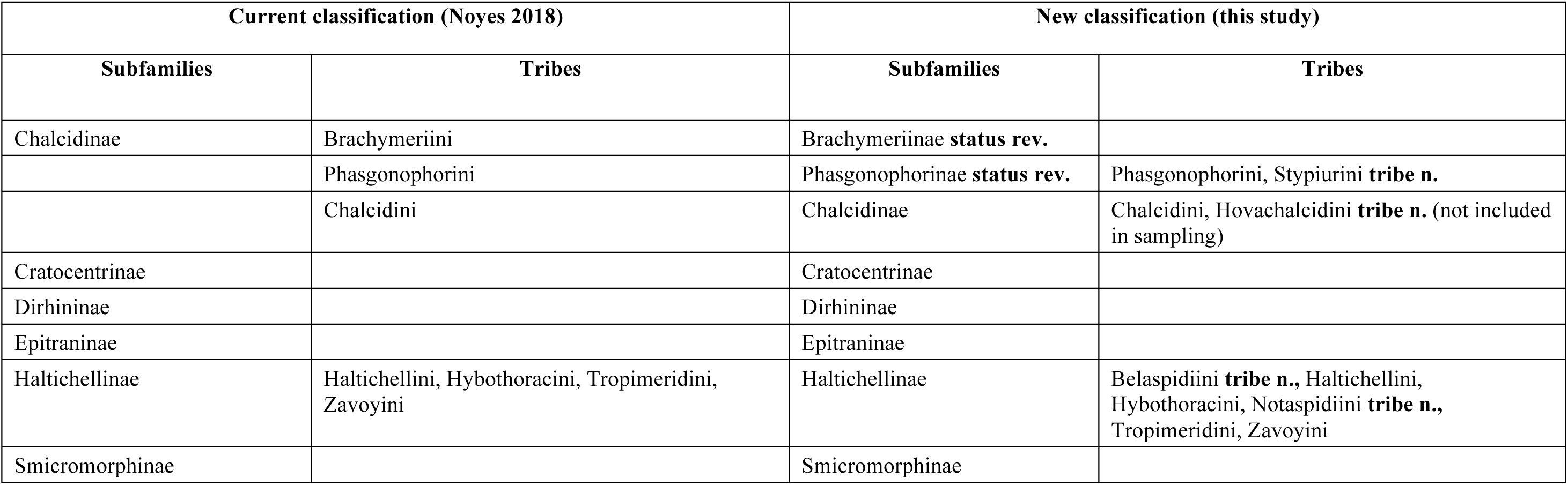
Current and new higher classification of the Chalcididae.

| Current classification (Noyes 2018) |  | New classification (this study) |  |
| --- | --- | --- | --- |
| Subfamilies | Tribes | Subfamilies | Tribes |
| Chalcidinae | Brachymeriini | Brachymeriinae **status rev.** |  |
|  | Phasgonophorini | Phasgonophorinae **status rev.** | Phasgonophorini, Stypiurini **tribe n.** |
|  | Chalcidini | Chalcidinae | Chalcidini, Hovachalcidini **tribe n.** (not included in sampling) |
| Cratocentrinae |  | Cratocentrinae |  |
| Dirhininae |  | Dirhininae |  |
| Epitraninae |  | Epitraninae |  |
| Haltichellinae | Haltichellini, Hybothoracini, Tropimeridini, Zavoyini | Haltichellinae | Belaspidiini **tribe n.**, Haltichellini, Hybothoracini, Notaspidiini **tribe n.**, Tropimeridini, Zavoyini |
| Smicromorphinae |  | Smicromorphinae |  |

We sequenced UCEs from 144 ingroup and 7 outgroup taxa, analyzed UCE /taxa properties, performed exploration of the phylogenetic tree space and used different analytical approaches to detect possible systematic bias and conflicts among gene regions. We interpreted results in the light of a thorough morphological analysis (130 characters encoded on 40 taxa representing all major lineages) to propose a new classification for the higher relationships within the family.

## MATERIALS AND METHODS

### Sampling for the molecular study

The data set contained 144 ingroup taxa (Table S1). All subfamilies and tribes as well as 82% of the world genera were included in the analysis. At the beginning of our study, 83 genera of Chalcididae were considered as valid and two as *incertae sedis* (*Antrochalcis* Kieffer, 1910 and *Chalcitiscus* Ghesquière, 1946, a fossil genus). The within tribe classification was not the purpose of this study and will be reviewed elsewhere. However, the examination of numerous specimens in several museums and personal collections, together with a review of the literature, suggested that we need to remove six genera from synonymy (*Eniacomorpha* Girault*, Hontalia* Cameron*, Invreia* Masi*, Pareniaca* Crawford*, Parinvreia* Steffan*, Peltochalcidia* Steffan). We also discovered seven new genera awaiting description. Thus, we now consider 94 genera of Chalcididae as valid, 77 of which were included in our study. Most missing genera are extremely rare, known mostly from the type series only. Relationships within the Chalcidoidea are unclear (Heraty et al. 2013) but morphological features (Lotfalizadeh et al. 2007) as well as preliminary results obtained with anchored hybrid enrichment and UCEs (work in prep) suggest that the Eurytomidae may be the sister family of the Chalcididae. As a consequence, five species of Eurytomidae representing three subfamilies, as well as two species of Cerocephalinae (that form a clade with Eurytomidae + Chalcididae in the chalcidoid tree from the anchored/UCE approach) were used as outgroups. We used dried specimens (35% of the samples), specimens kept in 75-96% EtOH (50%), as well as DNA extracts (15%) kept at −20°C for about the last 10 years. The oldest specimen was collected in 1951 though most specimens were collected in the last 20 years. The paratypes of two species: *Kopinata partirubra* Bouček, 1988 and *Chalcis vera* Bouček, 1974 housed in NHMUK, London were also included.

### DNA extraction, library preparation and sequencing

DNA was extracted non-destructively and vouchers were subsequently remounted on cards. DNA was extracted using the Qiagen DNeasy Blood and Tissue kit following manufacturer’s protocol with a few modifications detailed in Cruaud et al. (2019). Library preparation followed Cruaud et al. (2019). Briefly, input DNA was sheared to a size of ca 400 bp using the Bioruptor® Pico (Diagenode). End repair, 3’-end adenylation, adapters ligation and PCR enrichment were then performed with the NEBNext Ultra II DNA Library prep kit for Illumina (NEB). Adapters that contained amplification and Illumina sequencing primer sites, as well as a nucleotide barcode of 5 or 6 bp long for sample identification were used to tag samples. Pools of 16 samples were made at equimolar ratio. Each pool was enriched using the 2749 probes designed by Faircloth *et al*. (2015) using a MYbaits kit (Arbor Biosciences) and following manufacturer’s protocol. The hybridization reaction was run for 24h at 65°C. Post enrichment amplification was performed on beads with the KAPA Hifi HotStart ReadyMix. The enriched libraries were quantified with Qubit, an Agilent Bioanalizer and qPCR with the Library Quantification Kit - Illumina/Universal from KAPA (KK4824). They were then pooled at equimolar ratio. Paired-end sequencing (2*300bp) was performed on an Illumina Miseq platform at UMR AGAP (Montpellier, France).

### UCE data analysis (from raw reads to UCEs)

Quality control checks were performed on raw sequence data with FastQC v.0.11.2 (Andrews 2010). Quality filtering and adapter trimming were performed with Trimmomatic-0.36 (Bolger et al. 2014). Overlapping reads were merged using FLASH-1.2.11 (Magoc and Salzberg 2011). Demultiplexing was performed using a bash custom script (Cruaud et al. 2019). Assembly was performed with Trinity (Haas et al. 2013). UCE loci were identified with PhylUCE (Faircloth 2016) (*phyluce_assembly_match_contigs_to_probes*, *phyluce_assembly_get_match_counts --incomplete matrix*, *phyluce_assembly_get_fastas_from_match_counts*, all scripts were used with default parameters).

### UCE data analysis (from UCEs to phylogenetic trees)

UCEs present in more than 70% of the taxa were retained for analysis. UCEs were aligned with MAFFT v7.245 (-linsi option) (Katoh and Standley 2013). Sites with more than 50% gaps were removed from the alignments using the program seqtools implemented in the package PASTA (Mirarab et al. 2014b). Sequences exhibiting up to 500 gaps and 10 substitutions in pairwise alignments between all members of a UCE set were flagged by a custom script (available from https://github.com/DNAdiversity/UCE-Cross-Contamination-Check) as potential contaminations and reviewed before exclusion in subsequent analysis. Individual gene trees were inferred with raxmlHPC-PTHREADS-AVX version 8.2.4 (Stamatakis 2014). As α and the proportion of invariable sites cannot be optimized independently from each other (Gu et al. 1995) and following Stamatakis’ personal recommendation (RAxML manual), the proportion of invariant sites was not included in the model. A rapid bootstrap search (100 replicates) followed by a thorough ML search (-m GTRGAMMA) was performed. TreeShrink (Mai and Mirarab 2018) was used to detect and remove abnormally long branches in individual gene trees (e.g. due to misalignment). The per-species mode was used and preliminary analyses together with gene tree visualization showed that the optimal value of b (the percentage of tree diameter increasing from which a species should be removed) was 20. Indeed, as reported on TreeShrink tutorial, lower value of b led to the removal of species even when they were not on particularly long branches. All other parameters were set to default values. Once outliers were removed, UCEs were re-aligned with MAFFT-linsi and alignments were cleaned using seqtools. TreeShrink was used a second time to clean gene trees from long branches that might have been missed due to the presence of extra-long branches in the original gene trees. The per-species mode was used with b set to 20 but only the longest outliers were removed (k was set to 1). All other parameters were set to default values. Once outliers were removed, UCEs were re-aligned with MAFFT-linsi and alignments were cleaned using seqtools.

The final data set was analyzed using supermatrix and coalescent-based summary methods. Phylogenetic trees were estimated from the concatenated data set using Maximum Likelihood (ML) as implemented in raxmlHPC-PTHREADS-AVX version 8.2.4 (Stamatakis 2014) and IQTREE v1.5.3 (Nguyen et al. 2015). For the RAxML analysis, a rapid bootstrap search (100 replicates) followed by a thorough ML search (-m GTRGAMMA) was performed. IQTREE analysis employed an ML search with the GTR+G model with branch supports assessed with ultrafast bootstrap (Minh et al. 2013) and SH-aLRT test (Guindon et al. 2010) (1000 replicates). For both RAxML and IQTREE, two analyses were conducted: i) on the unpartitioned data set, ii) on the dataset partitioned according to the best partitioning scheme inferred by PartitionFinder 2.1.1 (Lanfear et al. 2017) using the Sliding-Window Site Characteristics (SWSC) method. This method has been recently developed to account for within-UCE heterogeneity (conserved core versus flanking, variable regions) (Tagliacollo et al. 2018). To fit with RAxML models and as α and the proportion of invariable sites cannot be optimized independently from each other, only the GTR+G model was evaluated. Branch lengths were considered as linked, model selection and partitioning scheme comparison were performed with the corrected Akaike Information Criterion (AICc) and the rclusterf algorithm. Finally, we used the GHOST model implemented in IQTREE to account for heterotachous evolution as it does not require *a priori* data partitioning, a possible source of model misspecification (Crotty et al. 2019). ASTRAL-III v5.6.1 (Zhang et al. 2018) and ASTRID (Vachaspati and Warnow 2015) were used to infer a species tree from the individual UCE trees inferred by RAxML. To improve accuracy (Zhang et al. 2018) nodes with BP support < 10 were collapsed in individual gene trees with the perl script AfterPhylo.pl (Zhu 2014). For the analysis with ASTRID and following recommendations for incomplete distance matrices, BioNJ was used to compute the phylogeny. Node supports were evaluated with local posterior probabilities (local PP) for the ASTRAL tree and 100 multi-locus bootstrapping (Seo 2008) for the ASTRID tree. Summary statistics were calculated using AMAS (Borowiec 2016). Tree annotation was performed with TreeGraph 2.13 (Stöver and Müller 2010). Correlation analysis between properties of the UCEs was performed with the R package Performance Analytics (Peterson and Carl 2018).

### Exploration of topological conflicts

New approaches have been recently developed to identify markers /sites supporting conflicting topologies that either make explicit assumptions about the biological basis of conflict (e.g. horizontal transfer, incomplete lineage sorting ILS, recombination, gene duplication) or not. Early approaches (e.g. Abby et al. 2010, Heled and Drummond 2010, Szöllősi and Daubin 2012) were computationally too intense to be implemented on large data sets. Furthermore, they are limited in their focus, constrained to one or two sources of incongruence and may not be robust to additional sources (Gori et al. 2016). Here we used more recent approaches, that do not rely on any assumptions about the biological basis of conflicts, and are computationally tractable on large data sets. This involves: partitioning of the data into coherent groups by clustering of tree to tree distances (Gori et al. 2016, Duchêne et al. 2018) or statistical tests of incongruence (Gene Genealogy Interrogation GGI; Arcila et al. 2017, Zhong and Betancur-R 2017, Betancur-R et al. 2019). For the clustering approach, geodesic distances between all pairs of trees were calculated with TreeCl (Gori et al. 2016) that requires overlapping set of samples but allows missing samples. Then, the optimal number of clusters obtained with the Partitioning Around Medoids (PAM) algorithm (Kaufman and Rousseeuw 1990) was estimated with the gap statistics as implemented in the R package cluster (Maechler et al. 2018, R Core Team 2018) (Kmax was set to 10 and number of bootstrap samples was set to 500 to keep computation time reasonable). The R package factoextra (Kassambara and Mundt 2017) was used to visualize clusters (ggplot2-based scatter plot (Wickham 2016)). Possible effects of missing data or gap content were evaluated by testing whether these variables were phylogenetically clustered on any of the conflicting topology. Tests were conducted using the K statistic (Blomberg et al. 2003) with the R package Phytools 0.6-00 (Revell 2012). The null expectation of K under no phylogenetic signal was generated by randomly shuffling the tips of the phylogeny 1000 times.

We also used Gene Genealogy Interrogation (GGI) to compute the relative support of the UCEs for each competing topology following Arcila et al. (2017). RAxML was used to infer trees from UCEs using each of the three competing topologies (Figure 1) as multi-furcating constraints (the structure of the backbone was fixed, but taxa within clades were free to move around). Thus, three constrained trees were inferred from each UCE. Per-site log likelihood scores for all constrained trees were calculated with RAxML and used to perform AU tests (Shimodaira 2002) in the package CONSEL (Shimodaira and Hasegawa 2001). The program makermt was used to generate K = 10 sets of bootstrap replicates with each set consisted of 100 000 replicates of the row sums. For each UCE, constrained trees were ranked by p-values of the AU test. The constrained tree with the highest p-value was considered as the best explanation of the data.

**Figure 1.**
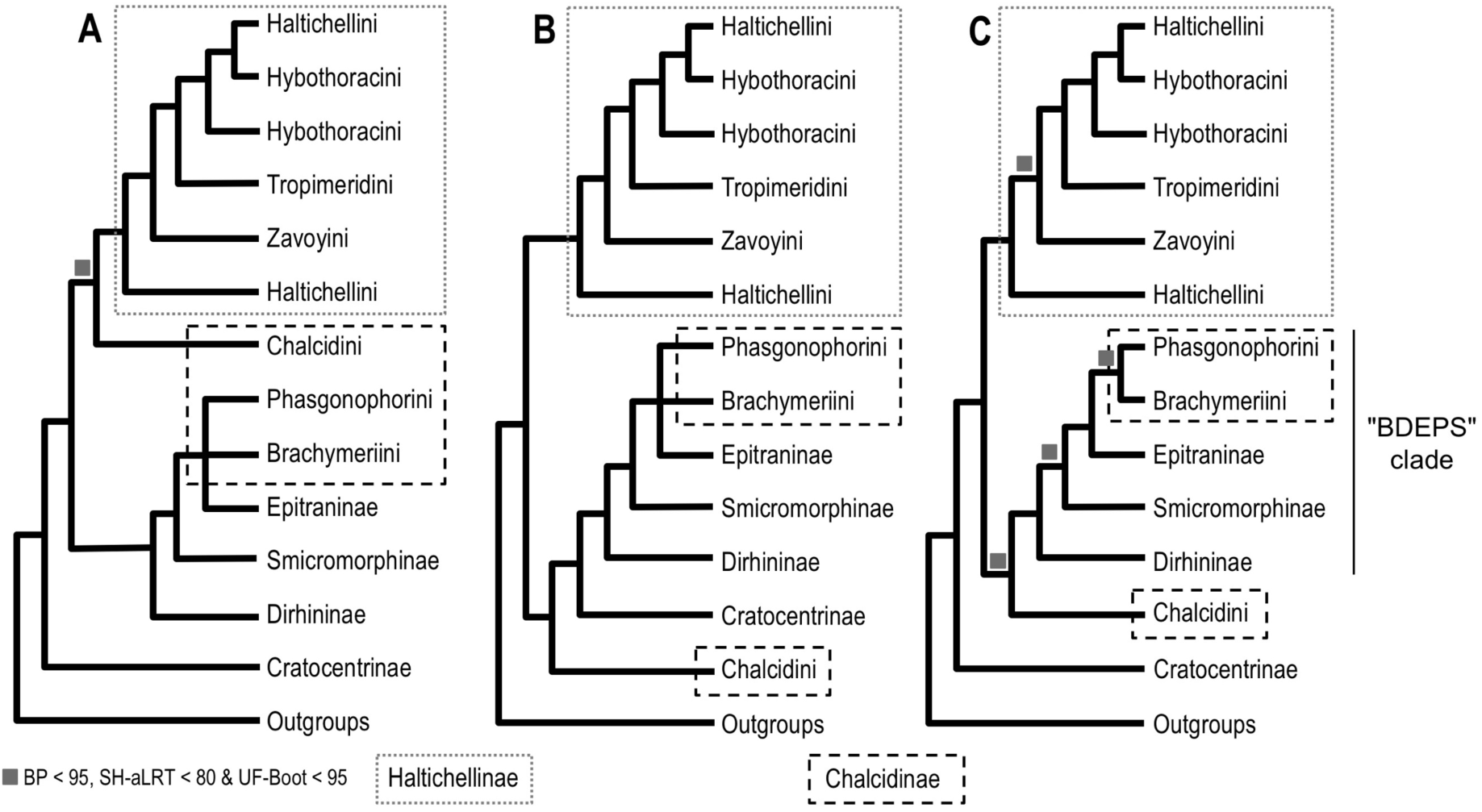
Summary of the topologies recovered from the UCE data set. The current classification is used to annotate the trees. Nodes are collapsed when BP support or SH-aLRT /UFBoot supports < 50. All nodes with BP > 95 and SH-aLRT > 80 /UFBoot > 95 unless specified with a colored box. **Topology A** is observed when the complete data set is analyzed with concatenation approaches (RAxML, IQTREE, GHOST, with or without partition). The node grouping Chalcidini with Haltichellinae is supported by moderate BP support (<80) and low SH-aLRT /UFBoot supports (<80/<95). Other nodes are highly supported. **Topology B** is observed when the complete data set is analyzed with ASTRAL. All nodes are highly supported (localPP=1.0). Topology B is also inferred when mid-point rooting instead of outgroup rooting is used to root the trees obtained with the concatenation approaches. **Topology C** is observed when the complete data set is analyzed with ASTRID. However, the node grouping Chalcidini with Dirhininae-Smicromorphinae-Epitraninae-Brachymeriini-Phasgonophorini is poorly supported (BP=32). Topology C is also observed in about 20% of the bootstrap trees obtained with the concatenation approaches. **Topology C** is inferred with concatenation approaches when UCEs (resp. nucleotides) with GC content strictly superior to 0.48 (resp. strictly superior to 0.57) are removed from the data set, though the node grouping Chalcidini with the Dirhininae-Smicromorphinae-Epitraninae-Brachymeriini-Phasgonophorini group is only moderately supported (BP=67 (resp. 80)). Finally, Topology C is observed when the set of input gene trees for the ASTRAL analysis is reduced to trees for which ingroups are monophyletic *Nota*: Current higher classification is used to annotate the trees. For brevity, we will refer to the clade grouping Brachymeriini + Dirhininae + Epitraninae + Phasgonophorini + Smicromorphinae as the “BDEPS” clade throughout text.

Finally, we examined whether compositional heterogeneity among loci and nucleotide positions as well as evolutionary rate heterogeneity among taxa could explain topological conflict. GC content and long branch (LB) score heterogeneity were calculated for each UCE and each taxon in all UCEs. GC content was calculated with AMAS (Borowiec 2016) and LB heterogeneity scores were calculated with TreSpEx (Struck 2014). In a given tree, the per sample LB score measures the percentage deviation of the patristics distance (PD) of a sample to all others, from the average PD across all samples (Struck 2014). For a given tree, the LB score heterogeneity is the standard deviation of the LB scores of the samples present in the tree. Thus, the LB score heterogeneity reflects the branch length heterogeneity of a given tree and is independent of the root of the tree. Consequently, calculation of LB score heterogeneities can help to prevent possible Long Branch Attraction (LBA) artefacts (Phillips et al. 2004, Bergsten 2005). Hierarchical clustering of taxa based on GC content and LB scores was performed with the R package cluster. The LBA artefact was also tested by removing outgroups from the analysis (Bergsten 2005).

### Morphological analyses

A matrix of 130 characters (Appendix S1) was assembled for 40 species that covered all major lineages of the Chalcididae (Appendix S2). The matrix was analyzed with parsimony (PAUP* version 4.0a; Swofford 2003), maximum likelihood (RAxML) and Bayesian (MrBayes-3.2.6; Ronquist et al. 2012) approaches. PAUP analyses were performed with unordered, equally weighted and non-additive characters. A traditional heuristic search was conducted using 1000 random addition sequences (RAS) to obtain an initial tree and “tree bisection and reconnection (TBR)” as branch swapping option. Ten trees were retained at each step. Robustness of the topology was assessed by bootstrap procedures (100 replicates; TBR RAS 10; one tree retained at each step). RAxML and MrBayes analyses were conducted with the Mk model (Lewis 2001) with only variable characters scored, equal state frequencies and assuming a gamma-distributed rate variation across characters. Robustness of the ML topology was assessed by bootstrap (100 replicates). Two independent runs of 1 million generations were run for the MrBayes analysis, with each run including a cold chain and three incrementally heated chains. The heating parameter was set to 0.02 in order to allow swap frequencies from 20% to 70%. Parameter values were sampled every 1000 generations. Convergence was examined in Tracer 1.6 (Rambaut et al. 2014) and the results were based on the pooled samples from the stationary phases of the 2 independent runs.

### Computational resources

Analyses were performed on a Dell PowerEdge T630 server with two 10-core Intel(R) Xeon(R) CPUs E5-2687W v3 @ 3.10GHz and on the Genotoul Cluster (INRA, Toulouse).

## RESULTS

### UCE data set

The final data set included 151 taxa (Table S1). No cross-contaminations were detected. In the first round, TreeShrink detected outlier long branches in 155 gene trees. Between 1 to 12 samples were flagged and removed per gene tree (average = 2 samples; Table S2). In the second round, TreeShrink detected outlier long branches in 28 gene trees. The final matrix (70% complete) included 538 UCEs and taxa were represented by 19 to 528 UCEs (median 478, Table S1). Eight taxa had more than 80% missing UCEs. The alignment contained 283,634 bp, 70.6% of which were parsimony informative. The percentage of missing data was 20.1%, the percentage of gaps was 16.1% and the GC content was 40.9%. Gap content for taxa (that can either result from the alignment of full length UCEs or capture of partial UCEs) ranged from to 2.2 to 37.0% (median 13.3%).

### Phylogenetic inference from the UCE data set

PartitionFinder2 used in combination with the SWSC method split the data set into 890 partitions. ML (Figures S1-S2, Appendix S3), ASTRAL (Figure S3) and ASTRID (Figure S4) trees were globally well resolved. The normalized quartet score of the ASTRAL tree was 0.91, which indicates a high degree of congruence between the species-tree and the input gene trees. Regardless of the method used, Chalcididae was always recovered as monophyletic with strong support. Of the six recognized subfamilies, only Chalcidinae was not monophyletic. With the exception of Hybothoracini and Haltichellini that were recovered as polyphyletic, all tribes were monophyletic with high support values. Of the 29 non-monotypic genera in the data set only 16 were recovered monophyletic.

Shallow and intermediate relationships were similar among analytical methods. About 10 unsupported topological conflicts were highlighted. Most of the time, conflicts involved taxa with a high level of missing UCEs (> 70%). Highly supported conflicts were observed in the deepest nodes of the phylogeny. While four moderately to highly supported clades were inferred by ML (Figure 1, topology A), Chalcididae clustered into two highly supported clades in the ASTRAL tree (Haltichellinae *versus* all other Chalcididae; Figure 1, topology B) and three poorly to highly supported clades were inferred by ASTRID (Figure 1, topology C). Notably, these conflicts were still observed when nodes with BP < 50 were collapsed in individual gene trees prior to species tree inference by ASTRAL (Figure S3; normalized quartet score = 0.98). For brevity, we will hereafter refer to the Brachymeriini + Dirhininae + Epitraninae + Phasgonophorini + Smicromorphinae clade (Figure 1) as the “BDEPS” clade.

### Exploration of topological conflicts

Analysis of bootstrap ML trees showed that topology A was recovered in ca 80% of the replicates and topology C was recovered in the remaining 20%. Topology B was recovered when ML trees were mid-point rooted (Figure S1C). Visualization of individual UCE trees revealed that ingroups were monophyletic in only 51 of the 538 trees (i.e 9.48%, Appendix S3). Interestingly, when the set of input trees for the ASTRAL analysis was reduced to these 51 trees, ASTRAL inferred topology C (Figure 3C). Neither missing data nor gap content appeared phylogenetically clustered on the ML, ASTRAL or ASTRID topologies (p-values > 0.05). The optimal number of clusters of loci as estimated by the gap statistics on the matrix of geodesic distances among individual gene trees was 1 (Figure S5) and the GGI approach showed that none of the topology was significantly preferred over the other (Figure 2). Topology A had a significantly best fit for 32 UCEs according to the GGI approach (p-value of the AU-test < 0.05).

**Figure 2.**
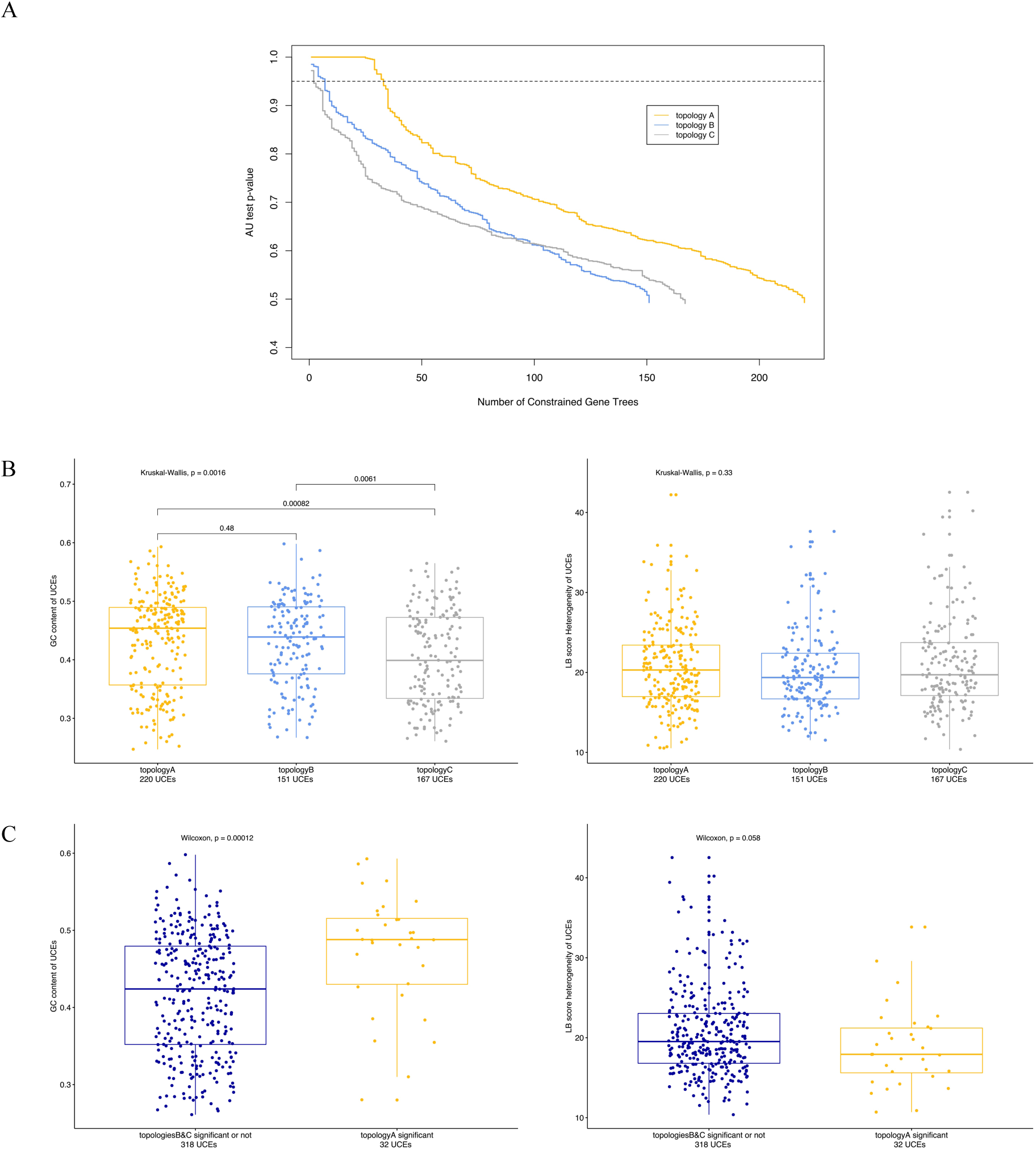
Results of the Gene Genealogy Interrogation approach (GGI) and correlation with UCE properties. A. Cumulative number of UCEs supporting each topology. Each UCE tree is constrained to fit with topologies A, B, and C and the approximately unbiased (AU) test is used to estimate which constrained tree shows the best fit (highest p-value) with the data. Values above the dashed line indicate that the preferred topology had a significantly better fit than the two alternatives (P < 0.05). B. Comparison of the GC content and LB score heterogeneity for the UCEs that support each topology. C. Comparison of the GC content and LB score heterogeneity for the UCEs that significantly support the topology A over topologies B and C.

**Figure 3.**
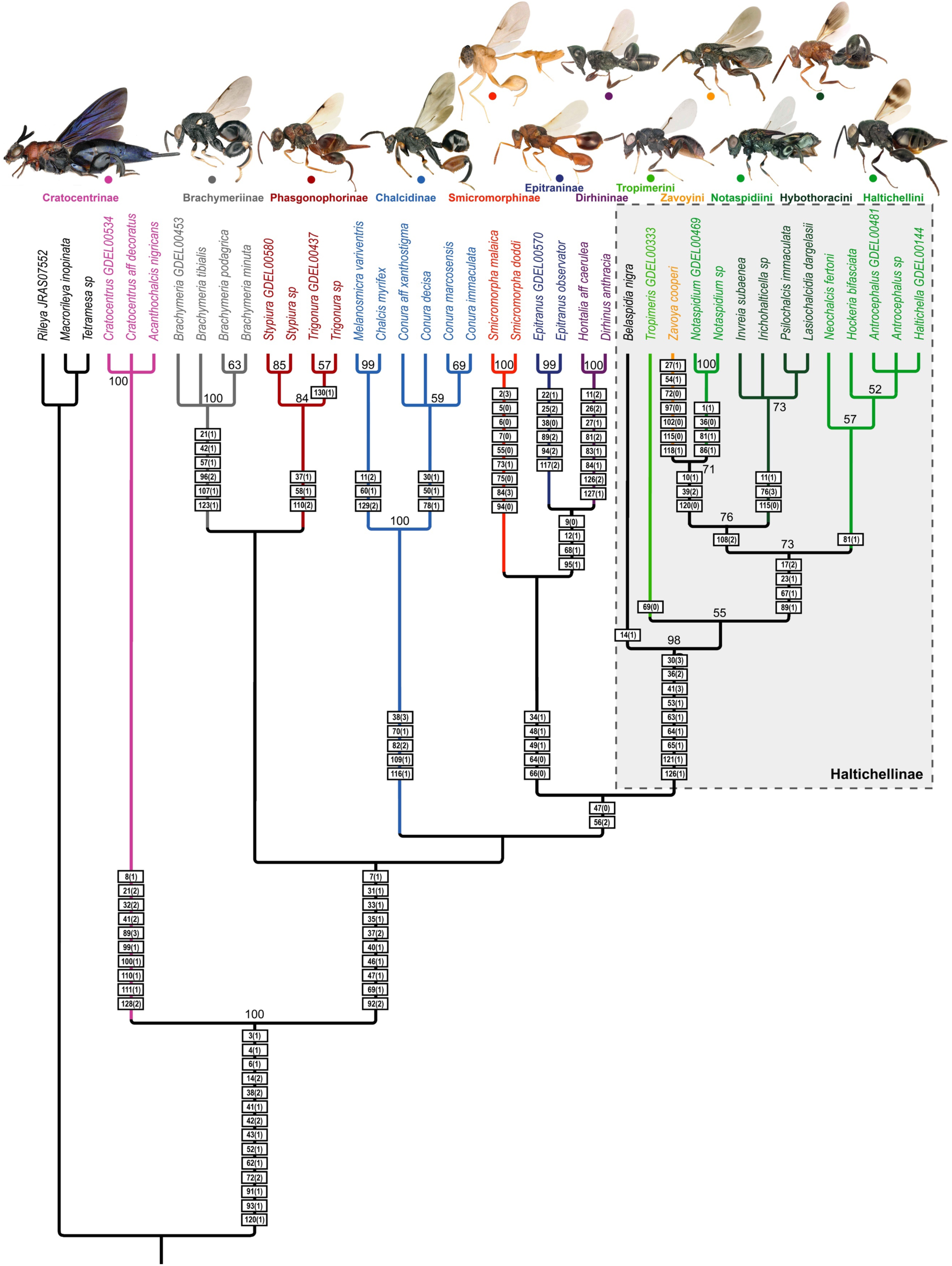
Results of the morphological analysis. Strict consensus of the 1439 equally parsimonious trees obtained with PAUP. Tree length = 330, CI=0.545, RI=0.840, RC=0.458. Bootstrap supports are depicted at nodes (100 replicates). Boxes on branches indicate the characters (states) that support the different nodes. The full list of morphological character and character states is provided in Appendix S1. A detailed discussion of the character states that support the different clades can be found in Appendix S5. The new classification proposed in this paper and associated color coding is used to annotate the tree.

Spearman’s rank correlation tests showed a significant negative correlation between GC content of the UCEs (Table S3) and the average support of individual gene trees (Figure S6). There was a higher difference between observed base composition and that predicted under the substitution model for GC-rich UCEs. The alpha parameter of the Gamma distribution was positively correlated with the number of sites informative for the parsimony (i.e. UCEs with more homogeneous rates among sites are more informative) and the average support of individual gene trees. LB score heterogeneity (Table S3) was negatively correlated with the number of sites informative for the parsimony and the average support of individual gene trees.

Hierarchical clustering of taxa based on the heterogeneity of LB scores (Table S4, Figure S7A) suggested that, with a few exceptions (Zavoyini, Tropimeridini, *Belaspidia*), the diversification dynamics of the Haltichellinae was somehow different compared to other subfamilies. Visual observation of the branch lengths of the ML trees confirmed this pattern (Figures S1-S2). Brachymeriini, Chalcidini, Cratocentrinae, Dirhininae, Epitraninae, Phasgonophorini and Smicromorphinae are supported by more elongated external branches. However, the dendrogram did not show obvious evidence to suspect that the position of Cratocentrinae as sister to the other Chalcididae in the topology A could result from an LBA artefact. In the dendrogram built from GC content (Table S4, Figure S7B), outgroups were not monophyletic and the Haltichellinae split into two groups. One of the subgroups shared similar properties as a few species of Chalcidini and Phasgonophorini.

Outgroup removal did not result in a shift of position for Cratocentrinae and the GHOST tree that accounts for heterotachous evolution was identical to other ML trees (Appendix S3). Joint analyses (GGI+UCE properties) revealed that heterogeneities of LB scores were not significantly different among the UCEs that supported either topology (Figure 2B-C). On the contrary, UCEs supporting topology C (with p-values significant or not) had a significantly lower GC content (Figure 2B). Furthermore, the 32 UCEs that significantly supported topology A had a significantly higher GC content (Figure 2C).

These results suggest that GC content may bias the results towards topology A. To test this hypothesis, data subsets were constructed by incrementally removing the most biased UCEs and nucleotide sites (Tables 2&3). To keep computation time reasonable, ML analyses were performed only with RAxML without partitioning. Indeed, all other approaches /models /softwares infer the same topology from the complete data set and are certainly subject to the same bias, if any. While ASTRAL and ASTRID inferred the same topology whatever the data subset analyzed, RAxML inferred topology C instead of topology A when UCEs with GC content > 0.48 (i.e. 28% of the UCEs) were removed from the data set (Table 2, Appendix S3). However, as already observed with the complete data set, bootstrap support for position of Chalcidini never reached 100%. The removal of nucleotide sites with GC > 0.57 (i.e 18.9% of the sites) also induced a shift from topology A to topology C for the ML analyses (Table 3, Appendix S3), though here again, the position of Chalcidini was not supported by 100% bootstrap (BP=80).

**Table 2.** Properties of the data subsets analyzed to detect possible systematic bias due to compositional heterogeneity among loci (GC content) and impact on the branching patterns of the deepest nodes.

|  | GC content threshold to remove UCEs |  |  |  |  |  |  |  |  |  |  |  |  |  |  | Complete data set |
| --- | --- | --- | --- | --- | --- | --- | --- | --- | --- | --- | --- | --- | --- | --- | --- | --- |
|  | > 0.41 | > 0.42 | > 0.43 | > 0.44 | > 0.45 | > 0.46 | > 0.47 | > 0.48 | > 0.49 | > 0.50 | > 0.51 | > 0.52 | > 0.53 | > 0.54 | > 0.55 | NA |
| Removed UCEs | 318<br>(59.1%) | 310<br>(57.6%) | 286<br>(53.2%) | 265<br>(49.3%) | 240<br>(44.6%) | 214<br>(39.8%) | 182<br>(33.8%) | 152<br>(28.3%) | 120<br>(22.3%) | 88<br>(16.4%) | 69<br>(12.8%) | 45<br>(8.36%) | 33<br>(6.13%) | 22<br>(4.09%) | 15<br>(2.79%) | 538<br>(0.00%) |
| Resulting alignment (bp) | 135,091 | 139,229 | 151,323 | 162,104 | 175,146 | 187,323 | 202,875 | 216,434 | 230,824 | 245,194 | 254,079 | 264,653 | 270,267 | 274,728 | 277,778 | 283,634 |
| GC content | 0.340 | 0.342 | 0.349 | 0.354 | 0.361 | 0.367 | 0.374 | 0.380 | 0.386 | 0.392 | 0.396 | 0.400 | 0.402 | 0.405 | 0.406 | 0.409 |
| Parsimony informative sites | 0.700 | 0.701 | 0.703 | 0.705 | 0.706 | 0.707 | 0.708 | 0.709 | 0.709 | 0.710 | 0.710 | 0.710 | 0.710 | 0.709 | 0.709 | 0.706 |
| Gap content | 0.162 | 0.162 | 0.161 | 0.162 | 0.161 | 0.162 | 0.162 | 0.162 | 0.162 | 0.162 | 0.162 | 0.161 | 0.162 | 0.162 | 0.162 | 0.161 |
| Missing data | 0.180 | 0.181 | 0.184 | 0.185 | 0.187 | 0.189 | 0.192 | 0.194 | 0.196 | 0.198 | 0.198 | 0.199 | 0.200 | 0.200 | 0.201 | 0.201 |
| Inferred Topology RAxML | C | C | C | C | C | C | C | C | A | A | A | A | A | A | A | A |
| Inferred Topology ASTRAL | B | B | B | B | B | B | B | B | B | B | B | B | B | B | B | B |
| Inferred Topology ASTRID | C | C | C | C | C | C | C | C | C | C | C | C | C | C | C | C |

**Table 3.** Properties of the data subsets analyzed to detect possible systematic bias due to compositional heterogeneity among nucleotide sites (GC content) and impact on the branching patterns of the deepest nodes.

|  | GC content threshold to remove nucleotide sites |  |  |  |  |  |  |  |  |  |  |  |  |  | Complete data set |
| --- | --- | --- | --- | --- | --- | --- | --- | --- | --- | --- | --- | --- | --- | --- | --- |
|  | > 0.50 | > 0.51 | > 0.52 | > 0.53 | > 0.54 | > 0.55 | > 0.56 | > 0.57 | > 0.58 | > 0.59 | > 0.60 | > 0.70 | > 0.80 | > 0.90 | NA |
| Removed sites | 63,436<br>(22.4%) | 61440<br>(21.7%) | 60,471<br>(21.3%) | 58711<br>(20.7%) | 57820<br>(20.4%) | 56,062<br>(19.8%) | 55,227<br>(19.5%) | 53551<br>(18.9%) | 52713<br>(18.6%) | 51063<br>(18.0%) | 50,242<br>(17.7%) | 37,791<br>(13.3%) | 14,040<br>(5.00%) | 221<br>(0.08%) | 0 |
| Resulting alignment (bp) | 220,198 | 222194 | 223,163 | 224923 | 225814 | 227,572 | 228,407 | 230083 | 230921 | 232571 | 233,392 | 245,843 | 269,594 | 283,413 | 283,634 |
| Inferred Topology RAxML | C | C | C | C | C | C | C | C | A | A | A | A | A | A | A |

### Analysis of morphological data

This study is the first in the whole superfamily Chalcidoidea to investigate, among other characters, the inner skeleton of the cephalic capsule and its external landmarks on the back of the head. Detailed results about character coding in the different taxa are provided in Appendix S4. Parsimony analysis of the 130 morphological characters (Appendix S1, S2) produced 1439 equally parsimonious trees (length = 330, consistency index (CI) = 0.545, retention index (RI) = 0.840, rescaled consistency index (RC) = 0.458; Figure 3). ML and Bayesian analyses recovered Cratocentrinae as sister to Phasgonophorini with moderate support (Figure S8). Otherwise, the ML, Bayesian and parsimony topologies were similar. Unsurprisingly, trees inferred from the morphological matrix were poorly supported as compared to the UCE trees. Just as UCE trees, morphological data support the monophyly of the family, the monophyly of all subfamilies except Chalcidinae, and the polyphyly of the tribes Haltichellini and Hybothoracini. In contrast with UCE trees, *Zavoya* was recovered sister to *Notaspidium* with high support and Dirhininae + Epitraninae formed a strongly supported clade. Tree backbones were too poorly supported to draw any firm conclusions. An in-depth analysis of the character states that support each clade is provided in Appendix S5. Mapping of character state transformation revealed that no one morphological character supported a sister taxa relationship between Chalcidini and Haltichellinae as observed in the UCE ML trees. Five characters unambiguously supported Cratocentrinae as sister to all other Chalcididae as observed in the UCE ML+ASTRID trees: mandibular base exposed, condyles elongate and visible externally, mouth margin not incised for reception of mandible (lateral to clypeus); subforaminal bridge depressed compared to postgena; lateral lamella on anterior tentorial arm narrow but with broad apical lobe; single metafurcal pit medially; two parallel and short submedian carinae present between metacoxae. Only two characters unambiguously supported a sister taxa relationship between Cratocentrinae, Chalcidini and the BDEPS clade as observed in the UCE ASTRAL tree: no raised carina on the inner margin of axillula and apex of metatibia diagonally truncate (Appendix S5). Consequently, morphological data provided no support for topology B and provides more support for topology C over topology A.

## DISCUSSION

### The power of UCEs to resolve the tree of life of poorly known groups

So far, UCEs have been successfully used to infer the phylogenies of a few groups of large to medium-sized insects (bees, wasps, ants and weevils: (Blaimer et al. 2015, Blaimer et al. 2016a, Branstetter et al. 2017a, Jesovnik et al. 2017, Prebus 2017, Van Dam et al. 2017, Bossert et al. 2019). Here, we highlight the power of UCEs for the exploration of hyperdiverse and poorly known groups of small to medium-sized chalcid wasps.

Interestingly, we were able to successfully use the universal Hymenoptera probe set designed by Faircloth et al (2015) to capture UCEs from chalcid wasps without any optimisation. This confirms the genericness of UCEs (Bossert and Danforth 2018). Besides Hymenoptera, universal probe sets have been designed for Arachnida, Coleoptera, Diptera, Hemiptera and Lepidoptera (Faircloth and Gilbert 2017) that could likely be used to infer relationships within many other poorly known insect groups. Easy to set-up and affordable protocols are freely available (http://ultraconserved.org; Faircloth et al. 2015, Cruaud et al. 2019), that do not require advanced skills or the use of an expensive robotic molecular biology platform. A software with a detailed tutorial (Faircloth 2016) exists for the easy analysis of raw data. Three situations can be distinguished in our study that may be encountered in other groups. First, when morphology is informative enough to circumscribe taxa and establish relationships, UCEs and morphology converge to the same results. Second, when morphology is not informative enough or misleading, UCEs are helpful to circumscribe taxa (e.g. genera or tribes) and clarify relationships within taxa. Additional studies of the morphology can then be performed and help discern characters that support the taxa. In summary, UCEs help to enable attainment of resolution between ancestry, convergent evolution or divergent evolution. We confirm that UCEs can be captured from museum specimens (Blaimer et al. 2016b, McCormack et al. 2016), which is key requirement when working on poorly known/rare groups, as sequencing of type specimens is often required to fix species names. Lastly, when neither morphology nor molecules are informative enough to resolve relationships, a systematic exploration of bias as well as the use of different analytical methods and a careful feedback assessment with morphological features is required.

### Resolving conflicts among methodological approaches

While there is a global agreement between morphology and molecules on the one hand, and between the different UCE analytical approaches on the other hand, there are a few conflicts that we discuss below.

First, there are unsupported conflicts between concatenation and gene tree reconciliation approaches for the position of taxa with a high level of missing UCEs (>90% e.g *Hastius* Schmitz, *Solenochalcidia* Steffan*, Pseudeniaca* Masi). The impact of missing data on phylogenetic inference has been widely discussed but no consensus has been reached.

Missing data are either considered as deleterious (Lemmon et al. 2009) or not problematic if enough informative characters are available to infer relationships (Wiens 2003, Wiens and Morrill 2011, Hosner et al. 2016, Streicher et al. 2016). Gene tree reconciliation approaches have been proven robust enough to a high, global level, of missing data (Nute et al. 2018). However, according to our knowledge of their morphology, the position of taxa with a high level of missing data is more accurate in the concatenation approach than in the gene tree reconciliation approaches.

Second, while shallow and intermediate relationships are similar among analytical approaches, two strongly supported conflicts are highlighted for the deepest nodes of the UCE phylogeny (Figure 1):

1. Cratocentrinae is recovered either as sister to all other Chalcididae (ML + ASTRID) or nested within Chalcididae (ASTRAL).
2. Chalcidini is either sister to Haltichellinae (ML), sister to the BDEPS clade (ASTRID) or sister to the BDEPS + Cratocentrinae clade (ASTRAL).

Spearman correlation tests suggest that evolutionary rate heterogeneity among taxa and nucleotide sites as well as compositional heterogeneity among UCEs could bias the analyses (Figure S6). Thus, the first hypothesis we explored was that the position of Cratocentrinae as sister to all other chalcidids in the ML tree resulted from an LBA artefact. Indeed, supermatrix approaches are more sensitive to LBA which, in addition, tend to be reinforced as more and more markers are considered (Boussau et al. 2014). However, hierarchical clustering of taxa properties (Figure S7), analysis with complex models that considers heterotachous evolution (GHOST, Appendix S3), outgroup removal analysis (Appendix S3) and the non-significant difference of scores of LB heterogeneity among UCEs that support topology A and other UCEs (Figure 2), show that LBA should be excluded. Sampling used in this study could certainly be improved but is representative of the group. More importantly, the outgroups used for this study are the closest relatives to the ingroups (work in prep., see the method section). Finally, morphological analysis reveals that Chalcididae minus Cratocentrinae possess apomorphic characters (Figure 3, Appendix S5) for which a scenario of loss and reacquisition implied by the ASTRAL tree seems unlikely.

On the other hand, our results suggest that the position of Chalcidini as sister to Haltichellinae in the ML trees, could result from a GC content bias. Indeed, in the morphological analysis, Chalcidini was never recovered sister to Haltichellinae. Besides, Chalcidini does not share any uniquely derived characters with Haltichellinae (Figure 3, Appendix S5). The largest difference between the observed GC content and that predicted under the substitution model is obtained for GC-rich UCEs and GGI analyses show that topology A is preferred by UCEs with a significantly higher GC content (Figure 2). Furthermore, hierarchical clustering of taxa based on their GC content show that some Haltichellinae share more similar GC content with Chalcidini or other subfamilies than with members of their own subfamily. GC-rich markers are more subject to recombination (Lartillot 2013, Romiguier and Roux 2017) and, as a consequence, to ILS (Pease and Hahn 2013). Thus, it is not surprising that only methods that are statistically consistent under the multi-species coalescent model (i.e. ILS-aware methods ASTRAL, ASTRID) do not recover Chalcidini sister to Haltichellinae from the analysis of the complete data set. ILS-aware methods have indeed been developed primarily to solve the deepest relationships for rapid radiations (e.g. birds (Mirarab et al. 2014a, Mirarab et al. 2014c). For the concatenation approach, when the most GC-biased UCEs (GC content >0.48; ca 28% of the total UCEs which corresponds to 23.7% of the total sites) or nucleotide sites (GC content > 0.57; 18.9% of the total sites) were removed, a sister taxa relationship between Chalcidini and the BDEPS clade was inferred, though with moderate bootstrap support (Tables 2-3, Figure 4, Appendix S3). This result agrees with other studies on reduced data sets, which revealed that UCEs can be GC-biased and support conflicting topologies (Sun et al. 2014, Bossert et al. 2017). However, we encourage the removal of the most GC biased nucleotide sites instead of the full UCEs to preserve phylogenetic signal in the analyzed subset.

**Figure 4.**
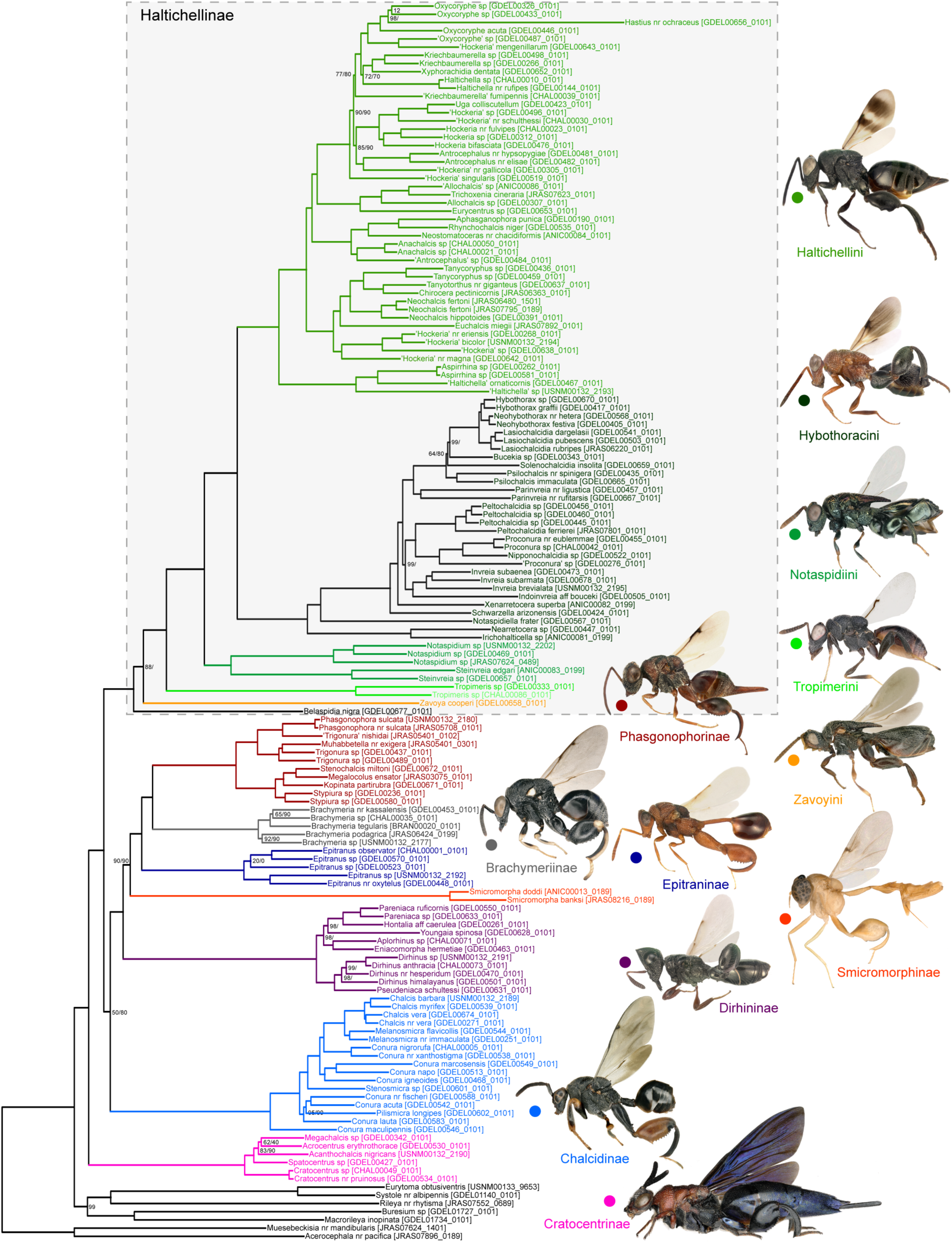
Preferred topology (topology C) showing the new higher classification of the Chalcididae. The RAxML tree inferred from the less biased UCEs (GC content ≦ 0.48) is used as a template and bootstrap support values less than 100 are reported at nodes as follows: RAxML on less biased UCEs /RAxML on less biased nucleotides (i.e. with GC content ≦ 0.57). This topology received the highest support from our morphological analysis (Appendix S5). The new classification proposed in this paper is used to annotate the tree. Color coding is similar to Figure 3. Within tribe classification including synonymization of invalid genera was not the purpose of this study and will be reviewed elsewhere. Single quotes indicate new genera awaiting description, the genus name used in the annotation is the one obtained when using current identification keys.

That said, one question still remains unanswered: how can we explain that Cratocentrinae are nested within Chalcididae in the ASTRAL tree? It is difficult to draw firm conclusions regarding this point. Our analyses suggest that individual UCE trees are not sufficiently resolved to allow a proper inference on the position of the outgroups from all bipartitions present in the input gene trees. Indeed, a majority of trees with intermixed outgroups and ingroups can complicate species tree inferences (Mai et al. 2017). The consequence of this lack of information is that outgroups are placed in an intermediate position. Indeed, the position of the outgroups in the ASTRAL tree corresponds to a mid-point rooting of the ML trees (Figure S1C) and topology C is recovered by ASTRAL when the set of input trees is reduced to those for which the ingroup is monophyletic. Outgroup choice is a difficult decision. Ideally, outgroups should be as close as possible to the ingroups to reduce LBA artefact while remaining sufficiently distantly-related to reduce impacts of ILS. However, an informed choice is often difficult if not impossible for hyperdiverse and poorly known groups.

Bringing together all species for phylogenetic inference is also impossible and such large data sets would be impossible to analyse with current methods (Philippe et al. 2017). This is why exploration of phylogenetic incongruence and systematic bias is particularly needed. The results of the ASTRAL analysis on the complete set of gene trees could also be explained by the nature of the marker used. UCEs are short which leads to poorly resolved gene trees with unsupported, incorrect bipartitions. Alignment cleaning (here indel removal) contributes to further reduce UCE length and, to some extent, to a loss of signal that should be quantified. Further research is needed to better understand possible drawback of tree reconciliation methods based on gene tree topology when they are used on UCEs and what could be the best strategy for alignment cleaning to preserve signal contained in gaps (Donath and Stadler 2018).

When it comes to the analysis of ancient groups that have undergone an explosive radiation like the Chalcidoidea (Heraty et al. 2013), another issue is the presence of monotypic or species-poor groups that are the only extant representatives of a long line of ancestors. These lineages are characterized by long external branches and insidious LBA artefacts may occur (not necessarily with the outgroups). On the morphological side, these taxa can be highly transformed and homologies between their features and those observed in the remaining species difficult to assess. Here, the position of two genera, *Zavoya* and *Smicromorpha,* remains ambiguous. Contrary to the results from the UCEs, *Zavoya* (3 species known) is strongly supported as sister to *Notaspidium* in the morphological tree, with which it shares many characters (Figures 4-S8, Appendix S4 & S5), though only one is an unambiguous synapomorphy. Additionally, this synapomorphy is a character loss (the ventral carinae of the metatibia are absent), that may confound interpretation of homology and relationships (Bleidorn 2007). For all other characters shared by *Zavoya* and *Notaspidium*, the same character state is observed only in species that do not belong to the Haltichellinae. These characters could thus be considered as local synapomorphies and, together with the absence of the ventral carinae on the metatibia, could reveal an undetected LBA artefact in our UCE analysis that may be reduced with an increasing sampling of *Zavoya, Notaspidium,* and Haltichellinae species. Further studies are nevertheless required to assess whether morphological convergence or systematic bias drive the position of *Zavoya*.

The position of *Smicromorpha* is also doubtful both in the morphological and the molecular trees. Even after UCE removal based on TreeShrink results (*Smicromorpha* is the most flagged taxon, Table S2), the long branch is still obvious (Appendix S3, Figure 4). Smicromorphinae are highly transformed parasitoids of weaver-ant larvae (*Oecophylla*, Formicinae) and character homologies are difficult to assess (Appendix S4-S5). During the day or at dusk, females *Smicromorpha* deposit their eggs on silk-spinning larva of weaver-ants held by workers to seal the leaves that are being pulled together by other workers when building their nest. Of the seven known species of *Smicromorpha*, two were reared from and three were collected flying around nests of *Oecophylla smaragdina* (Fabricius) in the Oriental and Australasian regions (Naumann 1986, Darling 2009). The biology of other species is unknown. While there is no fossil record for *Smicromorpha*, multiple fossils of *Oecophylla* are known from Europe (Barden 2017) suggesting that numerous species have become extinct since the Eocene Epoch (the estimate age of the first fossil is about 56 Ma). Interestingly, *Oecophylla* is also on a long branch and its putative relationships might therefore be artefactual (Ward et al. 2016). The highly specialized interaction *Smicromorpha*-*Oecophylla* and the numerous extinct lineages may explain the long branches and the difficulties encountered in correctly placing these taxa in the phylogenies. Morphological data support a sister taxa relationship between *Smicromorpha*, Dirhininae and Epitraninae but this clade is not recovered in the UCE tree. A close relationship between Epitraninae and *Smicromorpha* could be hypothesized based on UCEs and morphology /behavioral data (Appendix S5, Figure 4). Further studies are required to clarify the position of *Smicromorpha* in the chalcidid phylogenetic tree. The addition of more species of *Smicromorpha* and the inclusion of an undescribed genus, the probable sister taxon of *Smicromorpha* in the Afrotropical region, that we unfortunately failed to sequence due to poor specimen preservation, may improve the results. However, all these species are extremely rare and difficult to collect.

Finally, a conflict is also observed between morphological and molecular data for Dirhininae and Epitraninae. These subfamilies are recovered as sister taxa in the morphological tree while they are shown to be distantly related in the molecular trees. The sister taxa relationship is mostly supported by a closely related structure of the tentorium and other characters that are homoplastic especially those of the mesoscutellum, the fore wings and the hind legs. Therefore, their close relationships in our morphological analysis may reflect convergence.

### A new higher classification for the Chalcididae

The monophyly of the family is confirmed by our study. Chalcididae is supported by multiple synapomorphies: the presence of a genal carina and a postoral bridge, a sclerotized labrum with an exposed ventral plate, the parascutal and axillar carina ∩ -shaped, a metanotal scutellar arm reduced to a thin carina, the cubital vein present as a non-pigmented fold, and a petiole fused ventrally without suture (Appendix S5). Based on our results, we propose a revised higher classification (subfamilies and tribes) for the family (Table 1, Figures 3-4, Appendix S5). The within tribe classification was not the purpose of this study and will be reviewed elsewhere. To be conservative, we propose to keep subfamilies in their historical taxonomic rank, but to raise the tribes Phasgonophorini and Brachymeriini to subfamily rank. Within Haltichellinae, we recognise six major monophyletic groups that should be considered as tribes: Belaspidiini (**tribe n.**), Haltichellini, Hybothoracini, Notaspidiini (**tribe n.**), Tropimeridini and Zavoyini. Additionally, the subfamily Phasgonophorinae should include two tribes, Phasgonophorini (**status rev.**) and Stypiurini (**tribe n.**), to denote the corresponding sister monophyletic groups. Diagnoses of subfamilies and new tribes are given in Appendix S6.

### CONCLUSION

The increasing democratization of high-throughput sequencing technologies combined with the decreasing number of taxonomists result in a global overconfidence in phylogenetic hypotheses based on a large amount of molecular data. Furthermore, phylogenetic trees inferred from genome-scale data are usually highly supported which is often wrongly confused with accuracy, and falsely reinforces confidence in molecular results. Several authors have strongly advocated a systematic exploration of biases with different analytical methods. Indeed, this exploration may highlight better, alternative topologies that would not be revealed by a point and click approach. However, such studies are scarce, especially when data sets are composed of hundreds of taxa and genes, which makes computation time prohibitive and not compatible with the current publish or perish system. A thorough exploration of phylogenetic tree space may nevertheless reveal alternative hypotheses that are difficult to rank without independent sources of evidence. Our study highlights the power of a systematic exploration of biases to sort among conflicting phylogenomic hypotheses and the usefulness of a careful analysis of morphological features by expert taxonomist to corroborate (or not) the most likely topology. It may provide guidelines to build the tree of life of other hyperdiverse groups of animals on which little phylogenetic knowledge has been acquired, which is the rule rather than the exception in non-vertebrate taxa.

## Supporting information

AppendixS1

AppendixS2

AppendixS3

AppendixS4

AppendixS5

AppendixS6

## ACKOWLEDGEMENTS

We are grateful to Cedric Mariac and Leila Zekraoui (IRD, DIADE, France) for providing access to the Bioruptor; Audrey Weber (INRA, AGAP, France) for sequencing of the libraries and the Genotoul bioinformatics platform Toulouse Midi-Pyrenees, France for providing computing resources. We thank Natalie Dale-Skey Papilloud (NHM, London) and Nicole Fisher (ANIC, Canberra) for the loan of specimens as well as the Queensland government for collecting permits (WITK18248017-WITK18278817). This work is part of a large NSF project (Award#1555808) led by John Heraty (UC Riverside USA) that attempts to solve the phylogeny of the Chalcidoidea with NGS approaches and was funded by the INRA SPE department (recurrent funding to JYR and AC). USDA is an equal opportunity employer and provider. Trade names mentioned herein are for informational purposes only and do not imply endorsement by USDA.

## DATA ACCESSIBILITY

Demultiplexed cleaned reads are available as a NCBI Sequence Read Archive (ID#XXXX). Custom script to detect cross contamination is available from https://github.com/DNAdiversity/UCE-Cross-Contamination-Check.

## AUTHOR CONTRIBUTIONS

Designed the study: JYR, AC, GD; obtained funding: JYR, AC; contributed samples: GD, JYR, SF, SvN; identified samples: GD, JYR; performed lab work: SN, LS, SF; contribute sequences (UNSM samples): BBB, MG, SB; wrote script to detect cross-contaminations: SR; assembled the morphological matrix: GD; analyzed morphological data: JYR, GD, AC; analyzed molecular data: AC, JYR; provided help with R: MC, JPR; drafted the manuscript: AC, JYR, GD. All authors commented on the manuscript.

