## AppendixS1 for "Ultra-Conserved Elements and morphology reciprocally illuminate conflicting phylogenetic hypotheses in Chalcididae (Hymenoptera, Chalcidoidea)"

### Appendix S1. List of morphological characters

Within the following character list, numbers in brackets after character state descriptions are the consistency, retention and rescaled consistency indexes as inferred by PAUP* on one of the most parsimonious trees.

[1] Metallic body color. (0) present; (1) absent [CI= 0.500; RI= 0.500; RC=0.250].

[2] Sclerotization of body. (0) head and mesosoma moderately sclerotized, mostly reticulate or coriaceous, gaster slightly sclerotized, collapsing when dried; (1) head and mesosoma moderately sclerotized, mostly reticulate or coriaceous, gaster moderately sclerotized, not collapsing when dried; (2) head and mesosoma strongly sclerotized punctured and/or areolate, gaster distinctly sclerotized, not collapsing when dried; (3) head and mesosoma strongly sclerotized but metasoma collapsing when dried [CI=1.000; RI=1.000; RC=1.000].

[3] Relation between labrum and clypeus. (0) labrum overlapped by clypeus; (1) labrum exposed and abutting anterior to clypeal margin, not overlapped by clypeus [CI=1.000; RI=1.000; RC=1.000].

[4] Structure of labrum. (0) lightly sclerotized, without sculpture on surface; (1) plate-like, often with sculpture on surface [CI=1.000; RI=1.000; RC=1.000].

[5] Labral setae. (0) scattered across surface; (1) restricted to apical margin [CI=0.167; RI=0.615; RC=0.103].

[6] Mandibular base. (0) at least dorsally concealed by genal margin; (1) exposed, condyles elongate and visible externally, mouth margin not incised for reception of mandible (lateral to clypeus); (2) exposed, mouth margin thickened and incised for reception of dorsal corner of mandible [CI=0.500; RI=0.750; RC=0.375].

[7] Mouth margin above mandible. (0) mouth margin not incised for reception of mandible; (1) mouth margin thickened and incised for reception of dorsal corner of mandible [CI=0.500; RI=0.857; RC=0.429].

[8] Exposed muscle of mandible. (0) below the exposed plane of the mandible and not extending into an incision; (1) on the same plane with mandible and extending into incision on outer surface of mandible [CI=1.000; RI=1.000; RC=1.000].

[9] Number of teeth on left mandible. (0) three; (1) two; (2) four [CI=0.667; RI=0.500; RC=0.333].

[10] Number of teeth on the right mandible. (0) three; (1) two; (2) four [CI=0.250; RI=0.500; RC=0.500].

[11] Length of mandibular teeth. (0) ventral tooth about the same length as dorsal one; (1) ventral tooth much longer than dorsal one; (2) ventral tooth much shorter than dorsal one [CI=0.400; RI=0.727; RC=0.291].

[12] Orientation of mandibular teeth. (0) endodont; (1) exodont [CI=1.000; RI=1.000; RC=1.000].

[13] Channel on posterior surface of mandible. (0) absent; (1) present [CI=1.000; RI=1.000; RC=1.000].

[14] Delimitation of upper margin of clypeus. (0) impressed line (sulcus); (1) visible through change in sculpture; (2) step-like; (3) no evident limit. [CI=0.429; RI=0.000; RC=0.000].

[15] Lateral clypeal sulcus. (0) present; (1) absent [CI=1.000; RI=1.000; RC=1.000].

[16] Shape of clypeus. (0) about as broad as long; (1) more than 3 times as broad as long [CI=0.333; RI=0.667; RC=0.222].

[17] Position of toruli relative to oral cavity. (0) ventral margin of torulus in lower third of face, not adjacent to clypeus; (1) near middle of head or higher; (2) adjacent to clypeus. [CI=0.500; RI=0.882; RC=0.441].

[18] Anterior tentorial pits. (0) visible; (1) not visible externally [CI=0.333; RI=0.882; RC=0.294].

[19] Malar sulcus. (0) complete; (1) incomplete; (2) absent [CI=0.250; RI=0.700; RC=0.175].

[20] Genal carina. (0) absent; (1) angular but not carinate; (2) clearly present, raised [CI=1.000; RI=1.000; RC=1.000].

[21] Subapical genal tooth. (0) postgena not depressed above oral fossa, genal tooth or protrusion absent; (1) postgena distinctly depressed above oral fossa, hence genal carina forming a

protrusion at lateral corner of mouth; (2) postgena distinctly depressed above oral fossa, genal carina absent just above mouth corner, forming a tooth at some distance from it [CI=0.333; RI=0.714; RC=0.238].

**[22]** Frontal lobe below antennal toruli. (0) absent; (1) present [CI=1.000; RI=1.000; RC=1.000].

**[23]** Orientation of antennal toruli. (0) lateral and ventral margins of toruli not raised; (1) lateral and ventral margins of toruli raised [CI=1.000; RI=1.000; RC=1.000].

**[24]** Separation of toruli. (0) 1-2 times the torular diameter; (1) less than diameter of torulus. [CI=0.500; RI=0.800; RC=0.400].

**[25]** Interantennal projection from lateral view. (0) absent, not visible from lateral view or prominent but not discoid; (1) projection prominent and sulcate; (2) projection prominent, discoid, not or hardly sulcate on top [CI=0.667; RI=0.917; RC=0.611].

**[26]** Antennal scrobes. (0) present and shallow to moderately deep, never carinately margined; (1) present and deep, mostly carinately margined laterally [CI=0.143; RI=0.667; RC=0.095].

**[27]** Frontal horns. (0) absent; (1) present [CI= 0.500; RI= 0.500; RC=0.250].

**[28]** Postoccipital dorsolateral pit for attachment of cervical muscles. (0) absent; (1) present [CI=0.333; RI=0.714; RC=0.238].

**[29]** Postgenal groove. (0) absent; (1) present, not accompanied by postgenal lamina; (2) present, accompanied by distinct postgenal lamina [CI=0.333; RI=0.818; RC=0.273].

**[30]** Posterior tentorial pits/sulci. (0) present as sulci; (1) absent; (2) reduced to sulci linking posterior end of posterior tentorial arm (pta) to tentorial bridge pit (tbp)[CI=0.500; RI=0.833; RC=0.417].

**[31]** Hypostomal bridge (hb). (0) mostly subforaminal bridge expanded, hypostomal bridge quite reduced or not differentiate; (1) hypostomal bridge present, distinct from subforaminal bridge [CI=0.500; RI=0.667; RC=0.333].

**[32]** Level of subforaminal bridge. (0) at same level with postgena; (1) sunk down compared to postgena; (2) in front of postgenal bridge

**[33]** Width of hypostomal bridge relative to occipital foramen (of). (0) narrower than foramen; (1) as least as broad as foramen [CI=0.500; RI=0.800; RC=0.400].

**[34]** Median strip of ornamentation (subforaminal microtrichia of Burks & Heraty, 2015) on hypostomal bridge (hb). (0) present as a set of cuticular ridges or digitiform expansions; (1) absent or virtually so [CI=0.500; RI=0.750; RC=0.375].

**[35]** Width of median strip of ornamentation (mso). (0) strip narrow, less than one quarter width of hypostomal bridge; (1) strip wider e.g. at least one third width of hypostomal bridge [CI=0.500; RI=0.833; RC=0.417].

**[36]** Shape of hypostomal carina (hc). (0) forming a complete arch above the hypostomal bridge; (1) forming an incomplete arch above the hypostoma and hypostomal bridge; (2) extended above and joining the lateral edge postoccipital lateral arm (pola) [CI=0.333; RI=0.733; RC=0.244].

**[37]** Maxillary condyles (mc) (0) close to each other; (1) somewhat distant; (2) far from each other [CI=1.000; RI=1.000; RC=1.000].

**[38]** Cardo. (0) triangular; (1) elongate but with expanded, triangular apex; (2) stick-like; (3) fusiform [CI=1.000; RI=1.000; RC=1.000].

**[39]** Orientation of hypostoma and hypostomal bridge relative to subforaminal bridge (sfb). (0) bridges in same plan or hypostomal bridge hardly sloping; (1) bridges forming together an obtuse angle; (2) bridges forming together a right to acute angle [CI=0.400; RI=0.500; RC=0.200].

**[40]** Length of hypostomal bridge. (0) bridge short or vestigial, much shorter than subforaminal bridge; (1) bridge long to very long, at least as long as subforaminal bridge [CI=0.500; RI=0.667; RC=0.333].

**[41]** Lateral lamella (ll) on anterior tentorial arm (ata). (0) narrow; (1) moderately to very broad; (2) narrow but with broad apical lobe; (3) very broad and continuing on posterior tentorial arm (pta) [CI=1.000; RI=1.000; RC=1.000].

**[42]** Structure of posterior tentorial arm (pta). (0) a sclerotized triangular plate; (1) a simple, thick and strongly sclerotized arm; (2) including 2 arms, dorsally the pta itself, ventrally a subforaminal process (sfp) along surface of subforaminal bridge [CI=1.000; RI=1.000; RC=1.000].

**[43]** Insertion of dorsal tentorial arm (dta). (0) evidently above lower eye margin, at least at mid height of eye; (1) below, at or slightly above ventral eye margin [CI=1.000; RI=1.000; RC=1.000].

**[44]** ata-pta intersection. (0) far from base of maxillary condyles (mc); (1) near or at base of maxillary condyles [CI=0.500; RI=0.667; RC=0.333].

**[45]** dta-pta intersection. (0) at ata-pta intersection; (1) above ata-pta intersection. [CI=1.000; RI=1.000; RC=1.000].

**[46]** Position of ata apex relative to surface of hypostomal bridge. (0) ata not joining surface of hypostomal bridge; (1) apical part of ata forming process along lateral edge of hypostomal bridge [CI=1.000; RI=1.000; RC=1.000].

**[47]** Pits at dorsal end of ata. (0) absent; (1) present [CI=0.500; RI=0.875; RC=0.438].

**[48]** Position of dorsal end of posterior process (ppd). (0) on ventral margin of occipital foramen (of); (1) below ventral margin of OF. [CI=1.000; RI=1.000; RC=1.000].

**[49]** Pits at dorsal end of posterior process. (0) pits absent; (1) pits present [CI=0.500; RI=0.000; RC=0.00].

**[50]** Position of ventral end of posterior process (ppv). (0) intercepting pta; (1) on surface of subforaminal bridge [CI=0.333; RI=0.500; RC=0.167].

**[51]** Pits at ventral end of posterior process (pppv). (0) pits absent; (1) pits present [CI=0.333; RI=0.600; RC=0.200].

**[52]** Tentorial bridge (tb). (0) thick, well sclerotized and forming a T-like structure with anterior process (AP); (1) thin, slightly sclerotized and forming Y-like structure with AP. [CI=1.000; RI=1.000; RC=1.000].

**[53]** Postoccipital lateral arm (pola). (0) visible only on either side occipital foramen (OF); (1) joining ventrally the hypostomal carinae [CI=0.500; RI=0.889; RC=0.444].

**[54]** Number of separate claval segments in female. (0) three; (1) two; (2) one [CI=0.400; RI=0.842; RC=0.337].

**[55]** Multiporous plate sensilla (mps) position relative to antennal surface in female. (0) all mps raised above surface of flagellum; (1) at least some mps sunken [CI=0.500; RI=0.750; RC=0.375].

**[56]** Number of flagellomeres in male. (0) eleven; (1) nine; (2) seven [CI=0.333; RI=0.818; RC=0.273].

**[57]** Modified (long or spatulate) hairs on male flagellomeres. (0) absent; (1) present [CI=1.000; RI=1.000; RC=1.000].

**[58]** Posterior margin of pronotum. (0) straight or slightly concave; (1) strongly concave [CI=1.000; RI=1.000; RC=1.000].

**[59]** Relative position of tegula and humeral plate. (0) tegula not covering humeral plate; (1) tegula covering humeral plate [CI=0.500; RI=0.933; RC=0.467].

**[60]** Tegula position. (0) anterior corner of tegula abutting against marginal rim of mesoscutum; (1) tegula evidently tapering anteriorly, its anterior corner covered by lateral rim of mesoscutum [CI=1.000; RI=1.000; RC=1.000].

**[61]** Large setiferous cells on mesoscutum. (0) absent; (1) present [CI=1.000; RI=1.000; RC=1.000].

**[62]** Parascutal and axillar carinae. (0) V-shaped connection or not meeting at transscutal articulation; (1) U-shaped connection over tegula at transscutal articulation [CI=0.500; RI=0.667; RC=0.333].

**[63]** Axilla, tooth facing projecting anterior inner limit of axillula. (0) absent; (1) present [CI=1.000; RI=1.000; RC=1.000].

**[64]** Differentiation of axillula. (0) axillula absent or not differentiate; (1) axillula present [CI=1.000; RI=1.000; RC=1.000].

**[65]** Structure of inner margin of axillula. (0) no evident structure visible; (1) raised carina [CI=1.000; RI=1.000; RC=1.000].

**[66]** Frenal area of the mesoscutellum. (0) not marked dorsally; (1) defined completely across the mesoscutellum [CI=0.500; RI=0.875; RC=0.438].

**[67]** Frenum orientation. (0) frenum sloping to vertical; (1) frenum reflexed [CI=0.250; RI=0.769; RC=0.192].

**[68]** Declination of propodeum dorsal surface (lateral view). (0) sloping relative to longitudinal axis of mesonotum; (1) flat and in the same plane as mesonotum [CI=0.333; RI=0.750; RC=0.250].

**[69]** Shape of propodeal spiracle. (0) subcircular to elliptical; (1) slit-like [CI=0.500; RI=0.833; RC=0.417].
**[70]** Orientation of propodeal spiracle. (0) spiracle oblique; (1) spiracle vertical [CI=1.000; RI=1.000; RC= 1.000].
**[71]** Setose anterolateral areola on propodeum (horizontal areola between anterior margin of propodeum and spiracle). (0) absent; (1) present [CI=0.500; RI=0.889; RC=0.444].
**[72]** Posteroventral extension of pronotum. (0) not extending ventrally across prepectus; (1) with an extension that articulates or crosses the prepectus [CI=0.500; RI=0.667; RC=0.333].
**[73]** Emargination of pronotum around mesothoracic spiracle. (0) present, lateral panel emarginate around spiracle; (1) absent, lateral panel not emarginate around spiracle; (2) inconspicuous, lateral panel slightly emarginate and bearing dense patch of setae hiding spiracle [CI=1.000; RI=1.000; RC=1.000].
**[74]** Presence and shape of prosternal discrimen. (0) visible as a channel, a groove or a ridge; (1) absent [CI=0.125; RI=0.611; RC=0.076].
**[75]** Prosternum shape. (0) rounded between ventral and posterior surface or surfaces not distinct; (1) angulate or carinate at the limit between ventral and posterior surface; (2) with vertical lamina at limit between ventral and posterior surface [CI=0.500; RI=0.778; RC=0.389].
**[76]** Separation between anterior and posterior surface of prosternum. (0) absent or if present straight line; (1) median process, dentiform or lamelliform [CI=1.000; RI=1.000; RC=1.000].
**[77]** Profurcal pit. (0) present; (1) absent [CI=0.500; RI=0.800; RC=0.400].
**[78]** Lobes on margin of dorsal plate of profurca. (0) absent; (1) two lobes/projections present [CI=1.000; RI=1.000; RC=1.000].
**[79]** Ventral plate of profurca. (0) one surface visible; (1) two surfaces visible, forming a right angle together [CI=1.000; RI=1.000; RC=1.000].
**[80]** Apical stripe of prosternum. (0) apparent not sunken within body; (1) internal, sunken within body [CI=1.000; RI=1.000; RC=1.000].
**[81]** Ornamentation of ventral belt of prepectus. (0) medioventral areola present only; (1) small and sharp medioventral tooth; (2) large medioventral tooth [CI=0.667; RI=0.857 RC=0.571].
**[82]** Projection of mesothoracic spiracle. (0) not projecting, but visible externally; (1) hidden externally; (2) partly and hardly visible as hidden by a patch of hairs on posterior margin of pronotum [CI=1.000; RI=1.000; RC=1.000].
**[83]** Structure of lateral panel of prepectus. (0) without fovea or raised rim; (1) medially foveate, with posterior and/or dorsal rim and small anterodorsal projection [CI=1.000; RI=1.000; RC=1.000].
**[84]** Size and shape of exposed lateral panel of prepectus. (0) as tall or taller than long, more than half tegula length; (1) longer than tall, more than half tegula length; (2) small, less than half tegula length; (3) lateral panel not apparent [CI=0.500; RI=0.500; RC=0.250].
**[85]** Prepectus relationship to tegula. (0) prepectus reaching tegula; (1) prepectus not reaching tegula [CI=0.333; RI=0.667; RC=0.222].
**[86]** Setation of lateral panel of prepectus. (0) setose; (1) bare [CI=0.250; RI=0.833; RC=0.208].
**[87]** Posteroventral margin of prepectus. (0) ventral margin partially or completely fused medially with episternum; (1) completely separated from mesepisternum [CI=0.500; RI=0.929; RC=0.464].
**[88]** Mesepisternum: epicnemium. (0) absent; (1) present, and completely delimited by carina [CI=0.250; RI=0.727; RC=0.182].
**[89]** Mesothoracic discrimen. (0) sulcate or foveate; (1) raised carina or bump anteriorly, foveate groove posteriorly; (2) raised carina overall; (3) as anchor-like ornamentation with median carina [CI=0.750; RI=0.941; RC=0.706].
**[90]** Position of mesofurcal pit. (0) adjacent to mesocoxal cavity; (1) on mesotrochantinal plate [CI=0.500; RI=0.929; RC=0.464].
**[91]** Shape of metepimeron. (0) subtriangular; (1) broadly rectangular or squared [CI=0.500; RI=0.667; RC=0.333].
**[92]** Number of metafurcal pits. (0) lateral (paired) pits; (1) single median pit; (2) pits absent [CI=1.000; RI=1.000; RC= 1.000].

**[93]** Inner lamella of metadiscrimen. (0) As usual, not especially raised; (1) strongly raised [CI=1.000; RI=1.000; RC=1.000].

**[94]** Metepisternal ventral shelf. (0) absent; (1) present above mid coxae, short; (2) present above mid coxae, long [CI=0.667; RI=0.500; RC=0.333].

**[95]** Submedian teeth at posteroventral edge of metepisternal shelf. (0) absent; (1) present [CI=1.000; RI=1.000; RC=1.000].

**[96]** Ornamentation between metacoxae. (0) median groove; (1) single median carina present; (2) two submedian carinae present, converging posteriorly; (3) two submedian carinae present, parallel and short; (4) absent [CI=0.800; RI=0.947 RC=0.758].

**[97]** Carina connecting hind coxal and propodeal foramina. (0) absent; (1) present [CI=0.333; RI=0.333; RC=0.111].

**[98]** Number of setae on humeral plate. (0) more than four; (1) up to four [CI=1.000; RI=1.000; RC= 1.000].

**[99]** Basal posterior lobe of fore wing. (0) absent; (1) present [CI=1.000; RI=1.000; RC=1.000].

**[100]** Apicoventral tuft of setae on costal cell. (0) absent; (1) present [CI=1.000; RI=1.000; RC=1.000].

**[101]** Hyaline break on parastigma. (0) present; (1) absent [CI=0.250; RI=0.625; RC=0.156].

**[102]** Position of marginal vein relatively to front margin of wing. (0) along margin; (1) somewhat removed from margin [CI=0.500; RI=0.800; RC=0.400].

**[103]** Length of marginal vein of fore wing. (0) between 1-3 times stigmal vein + stigma length; (1) more than 3 times stigmal vein + stigma length; (2) more than 10 times length of stigmal vein plus stigma; (3) more than 10 times length of stigmal vein plus stigma [CI=0.667; RI=0.933 RC=0.622].

**[104]** Length of postmarginal vein of fore wing (fw). (0) longer than stigmal vein + stigma, shorter than costal cell; (1) 1-2 times as long as the stigmal vein; (2) absent or shorter than stigmal vein + stigma [CI=0.500; RI=0.900; RC=0.450].

**[105]** Uncus of stigmal vein of fore wing (0) present and projecting as a linear process; (1) absent [CI=0.333; RI=0.867; RC=0.289].

**[106]** Arrangement of uncal sensilla. (0) arranged in line; (1) grouped in a single cluster; (2) 2 pairs separated by a short space [CI=1.000; RI=1.000; RC=1.000].

**[107]** Location of basal hamulus. (0) near the others; (1) distant from the others [CI=1.000; RI=1.000; RC= 1.000].

**[108]** Shape of first hamulus. (0) curved towards wing surface, like the other hamuli; (1) straight or only slightly curved, with others strongly curved towards wing surface; (2) curved towards base of hind wing, with others curved towards wing surface [CI=0.400; RI=0.813; RC=0.325].

**[109]** Line of setation on posterior surface of procoxa. (0) absent; (1) present [CI=1.000; RI=1.000; RC= 1.000].

**[110]** Apical ornamentation of protibia. (0) without horizontally directed stout spur or elongation; (1) with horizontally directed socketed spur; (2) without socketed spur but distinctly expanded giving the appearance of a spur [CI=1.000; RI=1.000; RC=1.000].

**[111]** Pegs at apex of mesotibia. (0) absent; (1) present [CI=1.000; RI=1.000; RC=1.000].

**[112]** Shape of metacoxa. (0) coxa not enlarged; (1) coxa enlarged and/or elongate [CI=1.000; RI=1.000; RC=1.000].

**[113]** Shape of metafemur. (0) not enlarged (greater than or equal to 3× as long as broad); (1) enlarged (smaller than or equal to 3× as long as broad) [CI=1.000; RI=1.000; RC=1.000].

**[114]** Ventral ornamentation of metafemur. (0) without denticles or teeth ventrally; (1) with small, uniform teeth similar to blade of saw over most of length; (2) with large, regular, lobe like teeth [CI=0.667; RI=0.947; RC=0.632].

**[115]** Position of basal tooth of metafemur. (0) near base of femur; (1) at mid length of femur [CI=0.500; RI=0.875; RC=0.438].

**[116]** Line of stout bristles on inner surface of metafemur. (0) absent; (1) present [CI=1.000; RI=1.000; RC=1.000].

**[117]** Tarsal scrobe on apicodorsal surface of metatibia. (0) absent or short, never with tooth or protrusion above; (1) present, long, without tooth or protrusion above; (2) present, long, with a tooth or protrusion above [CI=0.500; RI=0.778; RC=0.389].

**[118]** Apex of metatibia, shape. (0) truncate at right angle; (1) diagonally truncate, posteroventral corner acute; (2) diagonally truncate, posteroventral corner elongated into spine [CI=0.667; RI=0.947; RC=0.632].

**[119]** Number of apical spurs on metatibia. (0) two spurs; (1) one spur; (2) no spur [CI=0.400; RI=0.842; RC=0.337].

**[120]** Ventral carinae of metatibia. (0) absent; (1) two ventral carinae present, one lateral and one mesal [CI=1.000; RI=1.000; RC=1.000].

**[121]** Carina on inner surface of metatibia. (0) absent; (1) present [CI=0.500; RI=0.900; RC=0.450].

**[122]** Tarsal claws. (0) simple; (1) pectinate, having 1-2 peg-like extra projection(s); (2) with basal tooth [CI=0.667; RI=0.833; RC=0.556]

**[123]** Spatulate seta on hind tarsal claw. (0) absent; (1) present [CI=1.000; RI=1.000; RC=1.000].

**[124]** Basal lamina on petiole. (0) absent; (1) present [CI=0.500; RI=0.857; RC=0.429].

**[125]** Insertion of petiole on propodeum. (0) at apex near metacoxal foramina; (1) at base of propodeum, near posterior margin of metanotum [CI=1.000; RI=1.000; RC=1.000].

**[126]** Relationship between petiole and propodeum. (0) body of petiole entering propodeum; (1) petiole with complete lamina surrounding petiolar foramen of propodeum; (2) petiole with ventral lamina abutting against petiolar foramen; (3) kneecap only entering propodeum [CI=1.000; RI=1.000; RC=1.000].

**[127]** Fusion of petiole with first gastral sternite in females. (0) not fused; (1) fused [CI=0.500; RI=0.750; RC=0.375].

**[128]** Transverse carina in front of cercal plates. (0) absent; (1) present, cerci inserted in foveae situated just behind the carina; (2) present, cerci situated posteriorly to carina [CI=0.400; RI=0.833; RC=0.333].

**[129]** Relative placement of hypopygium tip. (0) in basal half; (1) near tip of gaster; (2) somewhat beyond tip of gaster, apex densely setose [CI=0.500; RI=0.750; RC=0.375].

**[130]** Orientation of ovipositor sheaths. (0) straight; (1) curved downwards [CI=1.000; RI=1.000; RC= 1.000].

Burks R.A. & Heraty J.M. (2015) Subforaminal bridges in Hymenoptera (Insecta), with a focus on Chalcidoidea. *Arthropod Structure & Development*, 44, 173-19
