## AppendixS2 for "Ultra-Conserved Elements and morphology reciprocally illuminate conflicting phylogenetic hypotheses in Chalcididae (Hymenoptera, Chalcidoidea)"

### Appendix S2. Matrix of morphological characters

| Characters | *Tetramesa sp.* | *Rileya JRAS07552* | *Macrorileya inopinata* | *Acanthochalcis nigricans* | *Cratocentrus aff decoratus* | *Cratocentrus GDEL00534* | *Trigonura sp.* | *Trigonura GDEL00437* | *Stypiura sp.* | *Stypiura GDEL00580* | *Brachymeria minuta* | *Brachymeria podagrica* | *Brachymeria tibialis* | *Brachymeria GDEL00453* | *Chalcis myrifex* | *Melanosmicra variventris* | *Conura immaculata* | *Conura marcosensis* | *Conura decisa* | *Conura aff xanthostigma* | *Dirhinus anthracia* | *Hontalia aff. caerulea* | *Epitranus observator* | *Epitranus GDEL00570* | *Smicromorpha doddi* | *Smicromorpha malaica* | *Belaspidia nigra* | *Tropimeris GDEL00333* | *Neochalcis fertoni* | *Hockeria bifasciata* | *Haltichella GDEL00144* | *Antrocephalus sp.* | *Antrocephalus GDEL00481* | *Notaspidium sp.* | *Notaspidium GDEL00469* | *Zavoya cooperi* | *Irichohalticella sp* | *Lasiochalcidia dargelasii* | *Psilochalcis immaculata* | *Invreia subaenea* |
|---|---|---|---|---|---|---|---|---|---|---|---|---|---|---|---|---|---|---|---|---|---|---|---|---|---|---|---|---|---|---|---|---|---|---|---|---|---|---|---|---|
| **[1]** Metallic body color. (0) present; (1) absent | 1 | 1 | 1 | 1 | 1 | 1 | 1 | 1 | 1 | 1 | 1 | 1 | 1 | 1 | 1 | 1 | 1 | 1 | 1 | 1 | 1 | 0 | 1 | 1 | 1 | 1 | 1 | 1 | 1 | 1 | 1 | 1 | 1 | 0 | 0 | 1 | 1 | 1 | 1 | 1 |
| **[2]** Sclerotization of body. (0) head and mesosoma moderately sclerotized, mostly reticulate or coriaceous, gaster slightly sclerotized, collapsing when dried; (1) head and mesosoma moderately sclerotized, mostly reticulate or coriaceous, gaster moderately sclerotized, not collapsing when dried; (2) head and mesosoma strongly sclerotized punctured and/or areolate, gaster distinctly sclerotized, not collapsing when dried; (3) head and mesosoma strongly sclerotized but metasoma collapsing when dried | 0 | 1 | 1 | 2 | 2 | 2 | 2 | 2 | 2 | 2 | 2 | 2 | 2 | 2 | 2 | 2 | 2 | 2 | 2 | 2 | 2 | 2 | 2 | 2 | 3 | 3 | 2 | 2 | 2 | 2 | 2 | 2 | 2 | 2 | 2 | 2 | 2 | 2 | 2 | 2 |
| **[3]** Relation between labrum and clypeus. (0) labrum overlapped by clypeus; (1) labrum exposed and abutting anterior to clypeal margin, not overlapped by clypeus | 0 | 0 | 0 | 1 | 1 | 1 | 1 | 1 | 1 | 1 | 1 | 1 | 1 | 1 | 1 | 1 | 1 | 1 | 1 | 1 | 1 | 1 | 1 | 1 | 1 | 1 | 1 | 1 | 1 | 1 | 1 | 1 | 1 | 1 | 1 | 1 | 1 | 1 | 1 | 1 |
| **[4]** Structure of labrum. (0) lightly sclerotized, without sculpture on surface; (1) plate-like, often with sculpture on surface | 0 | 0 | 0 | 1 | 1 | 1 | 1 | 1 | 1 | 1 | 1 | 1 | 1 | 1 | 1 | 1 | 1 | 1 | 1 | 1 | 1 | 1 | 1 | 1 | 1 | 1 | 1 | 1 | 1 | 1 | 1 | 1 | 1 | 1 | 1 | 1 | 1 | 1 | 1 | 1 |
| **[5]** Labral setae. (0) scattered across surface; (1) restricted to apical margin | - | 0 | 0 | 0 | 1 | 1 | 1 | 1 | 1 | 1 | 1 | 1 | 1 | 1 | 0 | 1 | 1 | 1 | 1 | 1 | 0 | 1 | 1 | 1 | 0 | 0 | 1 | 1 | 1 | 1 | 1 | 1 | 1 | 0 | 0 | 0 | 0 | 0 | 0 | 0 |
| **[6]** Mandibular base. (0) at least dorsally concealed by genal margin; (1) exposed, condyles elongate and visible externally, mouth margin not incised for reception of mandible (lateral to clypeus); (2) exposed, mouth margin thickened and incised for reception of dorsal corner of mandible | 0 | 0 | 0 | 2 | 2 | 2 | 1 | 1 | 1 | 1 | 1 | 1 | 1 | 1 | 1 | 1 | 1 | 1 | 1 | 1 | 1 | 1 | 1 | 1 | 0 | 0 | 1 | 1 | 1 | 1 | 1 | 1 | 1 | 1 | 1 | 1 | 1 | 1 | 1 | 1 |
| **[7]** Mouth margin above mandible. (0) mouth margin not incised for reception of mandible; (1) mouth margin thickened and incised for reception of dorsal corner of mandible | 0 | 0 | 0 | 0 | 0 | 0 | 1 | 1 | 1 | 1 | 1 | 1 | 1 | 1 | 1 | 1 | 1 | 1 | 1 | 1 | 1 | 1 | 1 | 1 | 0 | 0 | 1 | 1 | 1 | 1 | 1 | 1 | 1 | 1 | 1 | 1 | 1 | 1 | 1 | 1 |
| **[8]** Exposed muscle of mandible. (0) below the exposed plane of the mandible and not extending into an incision; (1) on the same plane with mandible and extending into incision on outer surface of mandible | 0 | 0 | 0 | 1 | 1 | 1 | 0 | 0 | 0 | 0 | 0 | 0 | 0 | 0 | 0 | 0 | 0 | 0 | 0 | 0 | 0 | 0 | 0 | 0 | ? | ? | 0 | 0 | 0 | 0 | 0 | 0 | 0 | 0 | 0 | 0 | 0 | 0 | 0 | 0 |
| **[9]** Number of teeth on left mandible. (0) three; (1) two; (2) four | 0 | 2 | 0 | 1 | 1 | 1 | 1 | 1 | 1 | 1 | 1 | 1 | 1 | 1 | 1 | 1 | 1 | 1 | 1 | 1 | 1 | 1 | 1 | 1 | 1 | 1 | 1 | 1 | 1 | 1 | 1 | 1 | 1 | 1 | 1 | 1 | 1 | 0 | 1 | 1 |
| **[10]** Number of teeth on the right mandible. (0) three; (1) two; (2) four | 0 | 2 | 0 | 1 | 1 | 1 | 0 | 0 | 0 | 0 | 1 | 1 | 0 | 0 | 0 | 0 | 0 | 0 | 1 | 0 | 0 | 0 | 0 | 0 | 1 | 1 | 0 | 0 | 0 | 0 | 1 | 0 | 0 | 1 | 1 | 1 | 0 | 0 | 0 | 1 |
| **[11]** Length of mandibular teeth. (0) ventral tooth about the same length as dorsal one; (1) ventral tooth much longer than dorsal one; (2) ventral tooth much shorter than dorsal one | 0 | 0 | 0 | 0 | 0 | 0 | 0 | 0 | 0 | 0 | 0 | 0 | 0 | 0 | 2 | 2 | 0 | 0 | 0 | 0 | 2 | 2 | 1 | 1 | 1 | 1 | 0 | 0 | 0 | 0 | 1 | 0 | 0 | 0 | 0 | 0 | 1 | 1 | 1 | 1 |
| **[12]** Orientation of mandibular teeth. (0) endodont; (1) exod | 0 | 0 | 0 | 0 | 0 | 0 | 0 | 0 | 0 | 0 | 0 | 0 | 0 | 0 | 0 | 0 | 0 | 0 | 0 | 0 | 1 | 1 | 1 | 1 | 0 | 0 | 0 | 0 | 0 | 0 | 0 | 0 | 0 | 0 | 0 | 0 | 0 | 0 | 0 | 0 |
| **[13]** Channel on posterior surface of mandible. (0) absent; (1) present | 0 | 0 | 0 | 0 | 0 | 0 | 0 | 0 | 0 | 0 | 0 | 0 | 0 | 0 | 0 | 0 | 0 | 0 | 0 | 0 | 1 | ? | 1 | ? | - | - | 1 | ? | 1 | 1 | ? | 1 | ? | 1 | 1 | ? | ? | 1 | ? | 1 |
| **[14]** Delimitation of upper margin of clypeus. (0) impressed line (sulcus); (1) visible through change in sculpture; (2) step-like; (3) no evident limit. | ? | 3 | 0 | 2 | 2 | 2 | 2 | 2 | 2 | 2 | 2 | 2 | 2 | 0 | 0 | 2 | 2 | 2 | 2 | 2 | 2 | 2 | 2 | 2 | 3 | ? | 1 | 3 | 2 | 2 | 2 | 2 | 2 | 2 | 2 | 2 | 2 | 2 | 2 | 2 |
| **[15]** Lateral clypeal sulcus. (0) present; (1) absent | ? | 0 | 0 | 1 | 1 | 1 | 1 | 1 | 1 | 1 | 1 | 1 | 1 | 1 | 1 | 1 | 1 | 1 | 1 | 1 | 1 | 1 | - | 1 | 1 | 1 | 1 | 1 | 1 | 1 | 1 | 1 | 1 | 1 | 1 | 1 | 1 | 1 | 1 | 1 |
| **[16]** Shape of clypeus. (0) about as broad as long; (1) more than 3 times as broad as long | 0 | 0 | 0 | 1 | 1 | 1 | 1 | 1 | 1 | 1 | 1 | 1 | 1 | 1 | 1 | 1 | 1 | 1 | 1 | 1 | 0 | 0 | 1 | 1 | 0 | 0 | 1 | 1 | 1 | 1 | 1 | 1 | 1 | 1 | 1 | 1 | 1 | 1 | 1 | 1 |
| **[17]** Position of toruli relative to oral cavity. (0) ventral margin of torulus in lower third of face, not adjacent to clypeus; (1) near middle of head or higher; (2) adjacent to clypeus. | 1 | 1 | 1 | 0 | 0 | 0 | 0 | 0 | 1 | 1 | 0 | 0 | 0 | 0 | 0 | 0 | 1 | 1 | 0 | 0 | 0 | 0 | 0 | 0 | 0 | 0 | 0 | 0 | 2 | 2 | 2 | 2 | 2 | 2 | 2 | 2 | 2 | 2 | 2 | 2 |
| **[18]** Anterior tentorial pits. (0) visible; (1) not visible externally | 0 | 0 | 0 | 1 | 1 | 1 | 1 | 1 | 1 | 1 | 0 | 0 | 0 | 0 | 0 | 0 | 0 | 0 | 0 | 0 | 0 | 0 | ? | ? | 0 | 0 | 0 | 1 | 1 | 1 | 1 | 1 | 1 | 1 | 1 | 1 | 1 | 1 | 1 | 1 |
| **[19]** Malar sulcus. (0) complete; (1) incomplete; (2) absent | 0 | 0 | 0 | 1 | 1 | 1 | 2 | 2 | 2 | 2 | 0 | 0 | 0 | 0 | 0 | 0 | 0 | 0 | 0 | 0 | 2 | 2 | 1 | 1 | 2 | 2 | 2 | 2 | 1 | 0 | 0 | 0 | 0 | 2 | 2 | 2 | 2 | 2 | 0 | 1 |
| **[20]** Genal carina. (0) absent; (1) angular but not carinate; (2) clearly present, raised | 0 | 2 | 0 | 2 | 2 | 2 | 2 | 2 | 2 | 2 | 2 | 2 | 2 | 2 | 1 | 2 | 2 | 2 | 2 | 2 | 2 | 2 | 2 | 2 | 2 | 2 | 2 | 2 | 2 | 2 | 2 | 2 | 2 | 2 | 2 | 2 | 2 | 2 | 2 | 2 |
| **[21]** Subapical genal tooth. (0) postgena not depressed above oral fossa, genal tooth or protrusion absent; (1) postgena distinctly depressed above oral fossa, hence genal carina forming a protrusion at lateral corner of mouth; (2) postgena distinctly depressed above oral fossa, genal carina absent just above mouth corner, forming a tooth at some distance from it | 0 | 1 | 0 | 2 | 2 | 2 | 0 | 0 | 0 | 2 | 1 | 1 | 1 | 1 | 0 | 0 | 0 | 0 | 0 | 0 | 2 | 2 | 2 | 2 | 0 | 0 | 0 | 0 | 0 | 0 | 0 | 0 | 0 | 2 | 2 | 2 | 0 | 0 | 0 | 0 |
| **[22]** Frontal lobe below antennal toruli. (0) absent; (1) present | 0 | 0 | 0 | 0 | 0 | 0 | 0 | 0 | 0 | 0 | 0 | 0 | 0 | 0 | 0 | 0 | 0 | 0 | 0 | 0 | 0 | 0 | 1 | 1 | 0 | 0 | 0 | 0 | 0 | 0 | 0 | 0 | 0 | 0 | 0 | 0 | 0 | 0 | 0 | 0 |
| **[23]** Orientation of antennal toruli. (0) lateral and ventral margins of toruli not raised; (1) lateral and ventral margins of toruli raised | 0 | 0 | 0 | 0 | 0 | 0 | 0 | 0 | 0 | 0 | 0 | 0 | 0 | 0 | 0 | 0 | 0 | 0 | 0 | 0 | 0 | 0 | 0 | 0 | 0 | 0 | 0 | 0 | 1 | 1 | 1 | 1 | 1 | 1 | 1 | 1 | 1 | 1 | 1 | 1 |

| Characters | *Tetramesa sp.* | *Rileya JRAS07552* | *Macrorileya inopinata* | *Acanthochalcis nigricans* | *Cratocentrus aff decoratus* | *Cratocentrus GDEL00534* | *Trigonura sp.* | *Trigonura GDEL00437* | *Stypiura sp.* | *Stypiura GDEL00580* | *Brachymeria minuta* | *Brachymeria podagrica* | *Brachymeria tibialis* | *Brachymeria GDEL00453* | *Chalcis myrifex* | *Melanosmicra variventris* | *Conura immaculata* | *Conura marcosensis* | *Conura decisa* | *Conura aff xanthostigma* | *Dirhinus anthracia* | *Hontalia aff. caerulea* | *Epitranus observator* | *Epitranus GDEL00570* | *Smicromorpha doddi* | *Smicromorpha malaica* | *Belaspidia nigra* | *Tropimeris GDEL00333* | *Neochalcis fertoni* | *Hockeria bifasciata* | *Haltichella GDEL00144* | *Antrocephalus sp.* | *Antrocephalus GDEL00481* | *Notaspidium sp.* | *Notaspidium GDEL00469* | *Zavoya cooperi* | *Irichohalticella sp* | *Lasiochalcidia dargelasii* | *Psilochalcis immaculata* | *Invreia subaenea* |
|---|---|---|---|---|---|---|---|---|---|---|---|---|---|---|---|---|---|---|---|---|---|---|---|---|---|---|---|---|---|---|---|---|---|---|---|---|---|---|---|---|
| **[24]** Separation of toruli. (0) 1-2 times the torular diameter; (1) less than diameter of torulus. | 0 | 0 | 0 | 0 | 0 | 0 | 0 | 0 | 0 | 0 | 0 | 0 | 0 | 0 | 0 | 0 | 0 | 0 | 0 | 0 | 0 | 0 | 0 | 0 | 0 | 0 | 0 | 1 | 1 | 1 | 1 | 1 | 1 | 0 | 0 | 0 | 0 | 0 | 0 | 0 |
| **[25]** Interantennal projection from lateral view. (0) absent, not visible from lateral view or prominent but not discoid; (1) projection prominent and sulcate; (2) projection prominent, discoid, not or hardly sulcate on top | 0 | 0 | 0 | 0 | 0 | 0 | 0 | 0 | 0 | 0 | 0 | 0 | 0 | 0 | 0 | 0 | 0 | 0 | 0 | 0 | 0 | 0 | 2 | 2 | 0 | 0 | 0 | 0 | 1 | 2 | 2 | 2 | 2 | 1 | 1 | 1 | 1 | 1 | 1 | 1 |
| **[26]** Antennal scrobes. (0) present and shallow to moderately deep, never carinately margined; (1) present and deep, mostly carinately margined laterally | 0 | 0 | 0 | 1 | 1 | 1 | 1 | 1 | 1 | 1 | 1 | 1 | 1 | 1 | 0 | 0 | 0 | 0 | 0 | 0 | 1 | 1 | 0 | 0 | 1 | 1 | 0 | 0 | 0 | 0 | 1 | 1 | 1 | 1 | 1 | 1 | 0 | 0 | 0 | 0 |
| **[27]** Frontal horns. (0) absent; (1) present | 0 | 0 | 0 | 0 | 0 | 0 | 0 | 0 | 0 | 0 | 0 | 0 | 0 | 0 | 0 | 0 | 0 | 0 | 0 | 0 | 1 | 1 | 0 | 0 | 0 | 0 | 0 | 0 | 0 | 0 | 0 | 0 | 0 | 0 | 0 | 1 | 0 | 0 | 0 | 0 |
| **[28]** Postoccipital dorsolateral pit for attachment of cervical muscles. (0) absent; (1) present | 0 | 0 | 0 | 1 | 1 | 1 | 1 | 1 | 1 | 0 | 1 | ? | 1 | ? | 0 | ? | 0 | ? | 0 | ? | 0 | 0 | 0 | 0 | 0 | 0 | 0 | ? | 0 | 0 | 0 | 0 | 0 | 0 | ? | 0 | ? | 0 | ? | 0 |
| **[29]** Postgenal groove. (0) absent; (1) present, not accompanied by postgenal lamina; (2) present, accompanied by distinct postgenal lamina | 1 | 1 | 1 | 1 | 1 | ? | 1 | 1 | 1 | 0 | 1 | ? | 1 | ? | 0 | 0 | 0 | ? | 0 | ? | 0 | ? | 1 | 1 | 0 | ? | 0 | ? | 0 | 0 | 0 | 0 | 0 | 0 | ? | 0 | ? | 0 | ? | 0 |
| **[30]** Posterior tentorial pits/sulci. (0) present as sulci; (1) absent; (2) reduced to sulci linking posterior end of posterior tentorial arm (pta) to tentorial bridge pit (tbp) | 0 | 0 | 0 | - | - | ? | 0 | ? | 0 | ? | 1 | ? | 1 | ? | 0 | 0 | 1 | ? | 1 | ? | 1 | ? | 1 | ? | 1 | ? | 2 | ? | 2 | 2 | 2 | 2 | ? | 2 | ? | 2 | ? | 2 | ? | 2 |
| **[31]** Hypostomal bridge (hb). (0) mostly subforaminal bridge expanded, hypostomal bridge quite reduced or not differentiate; (1) hypostomal bridge present, distinct from subforaminal bridge | 0 | 1 | 0 | 0 | 0 | ? | 1 | ? | 1 | ? | 1 | ? | 1 | ? | 1 | 1 | 1 | ? | 1 | ? | 1 | ? | 1 | ? | 1 | ? | 1 | 1 | 1 | 1 | 1 | 1 | ? | 1 | ? | 1 | ? | 1 | ? | 1 |
| **[32]** Level of subforaminal bridge. (0) at same level with postgena; (1) sunk down compared to postgena; (2) in front of postgenal bridge | 0 | 0 | 0 | ? | 2 | ? | 1 | ? | 1 | ? | 1 | ? | 1 | ? | 1 | 1 | 1 | ? | 1 | ? | 1 | ? | 1 | ? | 0 | ? | 1 | ? | 1 | 1 | 1 | 1 | ? | 1 | ? | 1 | ? | 1 | ? | 1 |
| **[33]** Width of hypostomal bridge relative to occipital foramen (of). (0) narrower than foramen; (1) as least as broad as foramen | 0 | 0 | 0 | 0 | 0 | ? | 1 | ? | 1 | ? | 1 | ? | 1 | ? | 1 | 1 | 1 | ? | 1 | 1 | 1 | ? | 1 | ? | 0 | ? | 1 | 1 | 1 | 1 | 1 | 1 | ? | 1 | ? | 1 | ? | 1 | ? | 1 |
| **[34]** Median strip of ornamentation (subforaminal microtrichia of Burks & Heraty, 2015) on hypostomal bridge (hb). (0) present as a set of cuticular ridges or digitiform expansions; (1) absent or virtually so | 0 | 0 | 0 | 0 | 0 | ? | 0 | ? | 0 | ? | 0 | ? | 0 | ? | 0 | 0 | 0 | ? | 0 | ? | 1 | ? | 1 | ? | 1 | ? | 0 | ? | 0 | 0 | 0 | 0 | ? | 1 | ? | 1 | ? | 0 | ? | 0 |
| **[35]** Width of median strip of ornamentation (mso). (0) strip narrow, less than one quarter width of hypostomal bridge; (1) strip wider e.g. at least one third width of hypostomal bridge | 0 | 0 | 0 | 0 | 0 | ? | 0 | ? | 0 | ? | 1 | ? | 1 | ? | 1 | 1 | 1 | ? | 1 | ? | - | ? | - | ? | - | ? | 1 | ? | 1 | 1 | 1 | 1 | ? | - | ? | - | ? | 1 | ? | 1 |
| **[36]** Shape of hypostomal carina (hc). (0) forming a complete arch above the hypostomal bridge; (1) forming an incomplete arch above the hypostoma and hypostomal bridge; (2) extended above and joining the lateral edge postoccipital lateral arm (pola) | 0 | 1 | 0 | 1 | 1 | ? | 0 | ? | 0 | ? | 1 | ? | 1 | ? | 1 | 1 | 1 | ? | 0 | ? | 0 | ? | 0 | ? | 1 | ? | 2 | 2 | 2 | 2 | 2 | 2 | ? | 0 | ? | 2 | ? | 2 | ? | 2 |
| **[37]** Maxillary condyles (mc) (0) close to each other; (1) somewhat distant; (2) far from each other | 0 | 0 | 0 | 0 | 0 | ? | 1 | ? | 1 | ? | 2 | ? | 2 | ? | 2 | 2 | 2 | ? | 2 | ? | 2 | ? | 2 | ? | - | ? | 2 | ? | 2 | 2 | 2 | 2 | ? | 2 | ? | 2 | ? | 2 | ? | 2 |
| **[38]** Cardo. (0) triangular; (1) elongate but with expanded, triangular apex; (2) stick-like; (3) fusiform | ? | ? | 1 | ? | 2 | ? | 2 | ? | 2 | ? | 2 | ? | 2 | ? | 3 | 3 | 3 | ? | 3 | ? | 2 | ? | 0 | ? | ? | ? | ? | ? | ? | 2 | ? | 2 | ? | 2 | ? | ? | ? | ? | ? | 2 |
| **[39]** Orientation of hypostoma and hypostomal bridge relative to subforaminal bridge (sfb). (0) bridges in same plan or hypostomal bridge hardly sloping; (1) bridges forming together an obtuse angle; (2) bridges forming together a right to acute angle | 1 | ? | - | - | - | ? | 2 | ? | 2 | ? | 1 | ? | 1 | ? | 1 | 1 | 1 | ? | 1 | ? | 2 | ? | 2 | ? | 0 | ? | 1 | ? | 1 | 1 | 1 | 2 | ? | 2 | ? | 2 | ? | ? | ? | 1 |
| **[40]** Length of hypostomal bridge. (0) bridge short or vestigial, much shorter than subforaminal bridge; (1) bridge long to very long, at least as long as subforaminal bridge | ? | ? | 0 | 0 | 0 | ? | 1 | ? | 1 | ? | 1 | ? | 1 | ? | 1 | 1 | 1 | ? | 1 | ? | 1 | ? | 1 | ? | 0 | ? | 1 | ? | 1 | 1 | 1 | 1 | ? | 1 | ? | 1 | 1 | 1 | ? | 1 |
| **[41]** Lateral lamella (ll) on anterior tentorial arm (ata). (0) narrow; (1) moderately to very broad; (2) narrow but with broad apical lobe; (3) very broad and continuing on posterior tentorial arm (pta) | ? | ? | 0 | ? | 2 | ? | 1 | ? | 1 | ? | ? | ? | ? | ? | 1 | 1 | 1 | ? | 1 | ? | 1 | ? | 1 | ? | ? | ? | ? | ? | ? | 3 | ? | 3 | ? | 3 | ? | ? | ? | ? | ? | 3 |
| **[42]** Structure of posterior tentorial arm (pta). (0) a sclerotized triangular plate; (1) a simple, thick and strongly sclerotized arm; (2) including 2 arms, dorsally the pta itself, ventrally a subforaminal process (sfp) along surface of subforaminal bridge | 0 | ? | 0 | ? | 2 | ? | 2 | ? | 2 | ? | 1 | ? | 1 | ? | 2 | 2 | 2 | ? | 2 | ? | 2 | ? | 2 | ? | ? | ? | ? | ? | ? | 2 | ? | 2 | ? | 2 | ? | ? | ? | ? | ? | 2 |
| **[43]** Insertion of dorsal tentorial arm (dta). (0) evidently above lower eye margin, at least at mid height of eye; (1) below, at or slightly above ventral eye margin | 0 | ? | 0 | ? | 1 | ? | 1 | ? | 1 | ? | 1 | ? | 1 | ? | 1 | 1 | 1 | ? | 1 | 1 | 1 | ? | 1 | ? | 1 | ? | ? | ? | ? | 1 | ? | 1 | ? | 1 | ? | ? | ? | ? | ? | 1 |

| Characters | *Tetramesa sp.* | *Rileya JRAS07552* | *Macrorileya inopinata* | *Acanthochalcis nigricans* | *Cratocentrus aff decoratus* | *Cratocentrus GDEL00534* | *Trigonura sp.* | *Trigonura GDEL00437* | *Stypiura sp.* | *Stypiura GDEL00580* | *Brachymeria minuta* | *Brachymeria podagrica* | *Brachymeria tibialis* | *Brachymeria GDEL00453* | *Chalcis myrifex* | *Melanosmicra variventris* | *Conura immaculata* | *Conura marcosensis* | *Conura decisa* | *Conura aff xanthostigma* | *Dirhinus anthracia* | *Hontalia aff. caerulea* | *Epitranus observator* | *Epitranus GDEL00570* | *Smicromorpha doddi* | *Smicromorpha malaica* | *Belaspidia nigra* | *Tropimeris GDEL00333* | *Neochalcis fertoni* | *Hockeria bifasciata* | *Haltichella GDEL00144* | *Antrocephalus sp.* | *Antrocephalus GDEL00481* | *Notaspidium sp.* | *Notaspidium GDEL00469* | *Zavoya cooperi* | *Irichohalticella sp* | *Lasiochalcidia dargelasii* | *Psilochalcis immaculata* | *Invreia subaenea* |
|---|---|---|---|---|---|---|---|---|---|---|---|---|---|---|---|---|---|---|---|---|---|---|---|---|---|---|---|---|---|---|---|---|---|---|---|---|---|---|---|---|
| **[44]** ata-pta intersection. (0) far from base of maxillary condyles (mc); (1) near or at base of maxillary condyles | ? | ? | 0 | ? | 0 | ? | 0 | ? | 0 | ? | 1 | ? | 1 | ? | 1 | 1 | 1 | ? | 1 | ? | 1 | ? | 1 | ? | ? | ? | ? | ? | ? | 1 | ? | 1 | ? | 1 | ? | ? | ? | ? | ? | 1 |
| **[45]** dta-pta intersection. (0) at ata-pta intersection; (1) above ata-pta intersection. | ? | ? | 0 | ? | 0 | ? | 0 | ? | 0 | ? | 0 | ? | 0 | ? | 0 | 0 | 0 | ? | 0 | ? | 1 | ? | 1 | ? | ? | ? | ? | ? | ? | 0 | ? | 0 | ? | 0 | ? | ? | ? | ? | ? | 0 |
| **[46]** Position of ata apex relative to surface of hypostomal bridge. (0) ata not joining surface of hypostomal bridge; (1) apical part of ata forming process along lateral edge of hypostomal bridge | ? | ? | 0 | ? | - | ? | 1 | ? | 1 | ? | 1 | ? | 1 | ? | 1 | 1 | 1 | ? | 1 | ? | 1 | ? | 1 | ? | ? | ? | ? | ? | ? | 1 | ? | 1 | ? | 1 | ? | ? | ? | ? | ? | 1 |
| **[47]** Pits at dorsal end of ata. (0) absent; (1) present | 0 | ? | 0 | ? | - | - | 1 | ? | 1 | ? | 1 | ? | 1 | ? | 1 | 1 | 1 | 1 | 1 | ? | 0 | ? | 0 | ? | 0 | ? | 0 | ? | 0 | 0 | ? | 0 | ? | 0 | ? | ? | ? | 0 | ? | 0 |
| **[48]** Position of dorsal end of posterior process (ppd). (0) on ventral margin of occipital foramen (of); (1) below ventral margin of OF. | ? | ? | 0 | ? | 0 | ? | 0 | ? | 0 | ? | 0 | ? | 0 | ? | 0 | 0 | 0 | ? | 0 | ? | 1 | ? | 1 | ? | ? | ? | 0 | ? | 0 | 0 | 0 | 0 | ? | 0 | ? | 0 | ? | 0 | 0 | 0 |
| **[49]** Pits at dorsal end of posterior process. | ? | ? | 0 | ? | 0 | ? | 0 | ? | 0 | ? | 0 | ? | 0 | ? | 0 | 0 | 0 | ? | 0 | ? | 1 | ? | 1 | 0 | ? | ? | ? | ? | ? | 0 | ? | 0 | ? | 0 | ? | ? | ? | 0 | ? | 0 |
| **[50]** Position of ventral end of posterior process (ppv). (0) intercepting pta; (1) on surface of subforaminal bridge | ? | ? | 0 | ? | 0 | ? | 0 | ? | 0 | ? | 1 | ? | 1 | ? | 0 | 0 | 1 | ? | 1 | ? | 0 | ? | 0 | 1 | ? | ? | ? | ? | ? | 0 | ? | 0 | ? | 0 | ? | ? | ? | ? | ? | 0 |
| **[51]** Pits at ventral end of posterior process (pppv). (0) pits absent; (1) pits present | ? | ? | 1 | ? | - | ? | 0 | ? | 0 | ? | 1 | ? | 1 | ? | 0 | 0 | 1 | ? | 1 | ? | 0 | ? | 0 | ? | ? | ? | ? | ? | ? | 1 | ? | 1 | ? | 1 | ? | ? | ? | ? | ? | 1 |
| **[52]** Tentorial bridge (tb). (0) thick, well sclerotized and forming a T-like structure with anterior process (AP); (1) thin, slightly sclerotized and forming Y-like structure with AP. | ? | ? | 0 | ? | 1 | ? | 1 | ? | 1 | ? | 1 | ? | 1 | ? | 1 | 1 | 1 | ? | 1 | ? | 1 | ? | 1 | 1 | ? | ? | ? | ? | ? | 1 | ? | 1 | ? | 1 | ? | ? | ? | ? | ? | 1 |
| **[53]** Postoccipital lateral arm (pola). (0) visible only on either side occipital foramen (OF); (1) joining ventrally the hypostomal carinae | 0 | 0 | 0 | 0 | 0 | ? | 0 | ? | 0 | ? | 0 | ? | 0 | ? | 0 | 0 | 0 | 0 | 0 | ? | 0 | ? | 0 | 0 | ? | ? | 1 | 1 | 1 | 1 | 1 | 1 | ? | 0 | ? | 1 | 1 | 1 | ? | 1 |
| **[54]** Number of separate claval segments in female. (0) three; (1) two; (2) one | 0 | 0 | 0 | 1 | 1 | ? | 0 | 0 | 1 | 1 | 0 | 0 | 0 | 0 | 0 | 0 | 0 | 0 | 0 | 0 | 2 | 2 | 2 | 2 | 2 | 2 | 2 | 1 | 2 | 2 | 2 | 2 | 2 | 2 | ? | 1 | 2 | 2 | 2 | 2 |
| **[55]** Multiporous plate sensilla (mps) position relative to antennal surface in female. (0) all mps raised above surface of flagellum; (1) at least some mps sunken | 0 | 0 | 0 | 1 | 1 | 1 | 1 | 1 | 1 | 1 | 1 | 1 | 1 | 1 | 1 | 1 | 1 | 1 | 1 | 1 | 1 | 1 | 1 | 1 | 0 | 0 | 1 | 1 | 1 | 1 | 1 | 1 | 1 | 1 | 1 | 1 | 1 | 1 | 1 | 1 |
| **[56]** Number of flagellomeres in male. (0) eleven; (1) nine. | 1 | ? | 0 | 0 | 0 | ? | ? | ? | ? | ? | 0 | 0 | 0 | ? | 0 | 0 | 0 | 0 | 0 | 0 | 0 | ? | ? | ? | 1 | ? | 1 | ? | 1 | 1 | 1 | 1 | ? | 1 | ? | ? | 1 | 1 | 1 | 1 |
| **[57]** Modified (long or spatulate) hairs on male flagellomeres. (0) absent; (1) present | 0 | 0 | 0 | 0 | 0 | ? | ? | ? | ? | ? | 1 | 1 | 1 | 1 | 0 | 0 | 0 | 0 | 0 | 0 | 0 | ? | 0 | ? | 0 | ? | 0 | ? | 0 | 0 | 0 | 0 | ? | ? | ? | 0 | 0 | 0 | ? | 0 |
| **[58]** Posterior margin of pronotum. (0) straight or slightly concave; (1) strongly concave | 0 | 0 | 0 | 0 | 0 | 0 | 1 | 1 | 1 | 1 | 0 | 0 | 0 | 0 | 0 | 0 | 0 | 0 | 0 | 0 | 0 | 0 | 0 | 0 | 0 | 0 | 0 | 0 | 0 | 0 | 0 | 0 | 0 | 0 | 0 | 0 | 0 | 0 | 0 | 0 |
| **[59]** Relative position of tegula and humeral plate. (0) tegula not covering humeral plate; (1) tegula covering humeral plate | 0 | 0 | 0 | 0 | 0 | 0 | 0 | 0 | 0 | 0 | 1 | 1 | 1 | 1 | 0 | 0 | 0 | 0 | 0 | 0 | 1 | 1 | 1 | 1 | 1 | 1 | 1 | 1 | 1 | 1 | 1 | 1 | 1 | 1 | 1 | 1 | 1 | 1 | 1 | 1 |
| **[60]** Tegula position. (0) anterior corner of tegula abutting against marginal rim of mesoscutum; (1) tegula evidently tapering anteriorly, its anterior corner covered by lateral rim of mesoscutum [CI=1.000; RI=1.000; RC= 1.000]. | 0 | 0 | 0 | 0 | 0 | 0 | 0 | 0 | 0 | 0 | 0 | 0 | 0 | 0 | 1 | 1 | 0 | 0 | 0 | 0 | 0 | 0 | 0 | 0 | 0 | 0 | 0 | 0 | 0 | 0 | 0 | 0 | 0 | 0 | 0 | 0 | 0 | 0 | 0 | 0 |
| **[61]** Large setiferous cells on mesoscutum. (0) absent; (1) present | 0 | 0 | 0 | 1 | 1 | 1 | 1 | 1 | 1 | 1 | 1 | 1 | 1 | 1 | 1 | 1 | 1 | 1 | 1 | 1 | 1 | 1 | 1 | 1 | 1 | 1 | 1 | 1 | 1 | 1 | 1 | 1 | 1 | 1 | 1 | 1 | 1 | 1 | 1 | 1 |
| **[62]** Parascutal and axillar carinae. (0) V-shaped connection or not meeting at transscutal articulation; (1) U-shaped connection over tegula at transscutal articulation | 0 | 0 | 0 | 1 | 1 | 1 | 1 | 1 | 1 | 0 | 1 | 1 | 1 | 1 | 1 | 1 | 1 | 1 | 1 | 1 | 1 | 1 | 1 | 1 | 1 | 1 | 1 | 1 | 1 | 1 | 1 | 1 | 1 | 1 | 1 | 1 | 1 | 1 | 1 | 1 |
| **[63]** Axilla, tooth facing projecting anterior inner limit of axillula. (0) absent; (1) present | 0 | 0 | 0 | 0 | 0 | 0 | 0 | 0 | 0 | 0 | 0 | 0 | 0 | 0 | 0 | 0 | 0 | 0 | 0 | 0 | 0 | 0 | 0 | 0 | 0 | 0 | 1 | 1 | 1 | 1 | 1 | 1 | 1 | 1 | 1 | 1 | 1 | 1 | 1 | 1 |
| **[64]** Differentiation of axillula. (0) axillula absent or not differentiate; (1) axillula present | 0 | 0 | 0 | 0 | 0 | 0 | 0 | 0 | 0 | 0 | 0 | 0 | 0 | 0 | 0 | 0 | 0 | 0 | 0 | 0 | 0 | 0 | 0 | 0 | 0 | 0 | 1 | 1 | 1 | 1 | 1 | 1 | 1 | 1 | 1 | 1 | 1 | 1 | 1 | 1 |
| **[65]** Structure of inner margin of axillula. (0) no evident structure visible; (1) raised carina | - | - | - | 0 | 0 | 0 | 0 | 0 | 0 | 0 | 0 | 0 | 0 | 0 | 0 | 0 | 0 | 0 | 0 | 0 | 0 | 0 | 0 | 0 | 0 | 0 | 1 | 1 | 1 | 1 | 1 | 1 | 1 | 1 | 1 | 1 | 1 | 1 | 1 | 1 |
| **[66]** Frenal area of the mesoscutellum. (0) not marked dorsally; (1) defined completely across the mesoscutellum | 0 | 0 | 0 | 1 | 1 | 1 | 1 | 1 | 1 | 1 | 1 | 1 | 1 | 1 | 1 | 1 | 1 | 1 | 1 | 1 | 0 | 0 | 0 | 0 | 0 | 0 | 1 | 1 | 1 | 1 | 1 | 1 | 1 | 1 | 1 | 1 | 1 | 1 | 1 | 1 |
| **[67]** Frenum orientation. (0) frenum sloping to vertical; (1) frenum reflexed | - | - | - | 0 | 0 | 0 | 0 | 0 | 0 | 1 | 1 | 1 | 1 | 1 | 1 | 1 | 1 | 1 | 1 | 1 | 0 | 0 | 0 | 0 | 0 | 0 | 0 | 1 | 1 | 1 | 1 | 1 | 1 | 1 | 1 | 1 | 1 | 1 | 1 | 1 |
| **[68]** Declination of propodeum dorsal surface (lateral view). (0) sloping relative to longitudinal axis of mesonotum; (1) flat and in the same plane as mesonotum | 0 | 0 | 0 | 0 | 0 | 0 | 0 | 0 | 0 | 0 | 0 | 0 | 0 | 0 | 0 | 0 | 0 | 0 | 0 | 0 | 1 | 1 | 1 | 1 | 0 | 0 | 0 | 0 | 0 | 0 | 0 | 0 | 0 | 1 | 1 | 1 | 1 | 0 | 0 | 1 |
| **[69]** Shape of propodeal spiracle. (0) subcircular to elliptical; (1) slit-like | 0 | 0 | 0 | 0 | 0 | 0 | 1 | 1 | 1 | 1 | 1 | 1 | 1 | 1 | 1 | 1 | 1 | 1 | 1 | 1 | 1 | 1 | 1 | 1 | 1 | 1 | 1 | 0 | 1 | 1 | 1 | 1 | 1 | 1 | 1 | 1 | 1 | 1 | 1 | 1 |
| **[70]** Orientation of propodeal spiracle. (0) spiracle oblique; (1) spiracle vertical | 0 | 0 | 0 | 0 | 0 | 0 | 0 | 0 | 0 | 0 | 0 | 0 | 0 | 0 | 1 | 1 | 1 | 1 | 1 | 1 | 0 | 0 | 0 | 0 | 0 | 0 | 0 | 0 | 0 | 0 | 0 | 0 | 0 | 0 | 0 | 0 | 0 | 0 | 0 | 0 |
| **[71]** Setose anterolateral areola on propodeum (horizontal areola between anterior margin of propodeum and spiracle). (0) absent; (1) present | 0 | 0 | 0 | 0 | 0 | 0 | 0 | 0 | 0 | 0 | 1 | 1 | 1 | 1 | 1 | 1 | 1 | 1 | 1 | 1 | 0 | 0 | 0 | 0 | 0 | 0 | 0 | 0 | 0 | 0 | 0 | 0 | 0 | 0 | 0 | 0 | 0 | 0 | 0 | 0 |
| **[72]** Posteroventral extension of pronotum. (0) not extending ventrally across prepectus; (1) with an extension that articulates or crosses the prepectus | 0 | 0 | 0 | 1 | 1 | 1 | 1 | 1 | 1 | 1 | 1 | 1 | 1 | 1 | 1 | 1 | 1 | 1 | 1 | 1 | 1 | ? | 1 | 1 | 1 | 1 | 1 | 1 | 1 | 1 | 1 | 1 | 1 | 1 | 1 | 0 | 1 | 1 | 1 | 1 |

| Characters | *Tetramesa sp.* | *Rileya JRAS07552* | *Macrorileya inopinata* | *Acanthochalcis nigricans* | *Cratocentrus aff decoratus* | *Cratocentrus GDEL00534* | *Trigonura sp.* | *Trigonura GDEL00437* | *Stypiura sp.* | *Stypiura GDEL00580* | *Brachymeria minuta* | *Brachymeria podagrica* | *Brachymeria tibialis* | *Brachymeria GDEL00453* | *Chalcis myrifex* | *Melanosmicra variventris* | *Conura immaculata* | *Conura marcosensis* | *Conura decisa* | *Conura aff xanthostigma* | *Dirhinus anthracia* | *Hontalia aff. caerulea* | *Epitranus observator* | *Epitranus GDEL00570* | *Smicromorpha doddi* | *Smicromorpha malaica* | *Belaspidia nigra* | *Tropimeris GDEL00333* | *Neochalcis fertoni* | *Hockeria bifasciata* | *Haltichella GDEL00144* | *Antrocephalus sp.* | *Antrocephalus GDEL00481* | *Notaspidium sp.* | *Notaspidium GDEL00469* | *Zavoya cooperi* | *Irichohalticella sp.* | *Lasiochalcidia dargelasii* | *Psilochalcis immaculata* | *Invreia subaenea* |
|---|---|---|---|---|---|---|---|---|---|---|---|---|---|---|---|---|---|---|---|---|---|---|---|---|---|---|---|---|---|---|---|---|---|---|---|---|---|---|---|---|
| **[73]** Emargination of pronotum around mesothoracic spiracle. (0) present, lateral panel emarginate around spiracle; (1) absent, lateral panel not emarginate around spiracle; (2) inconspicuous, lateral panel slightly emarginate and bearing dense patch of setae hiding spiracle | 1 | 1 | 1 | 0 | 0 | 0 | 1 | 1 | 1 | 1 | 1 | 1 | 1 | 1 | 2 | 2 | 2 | 2 | 1 | 2 | 1 | 1 | 1 | 1 | 1 | 1 | 1 | 1 | 1 | 1 | 1 | 1 | 1 | 1 | 1 | 1 | 1 | 1 | 1 | 1 |
| **[74]** Presence and shape of prosternal discrimen. (0) visible as a channel, a groove or a ridge; (1) absent | 1 | 1 | 1 | 0 | 0 | 0 | 1 | 1 | 1 | 1 | 0 | 0 | 0 | 0 | 1 | 1 | 1 | ? | 1 | ? | 0 | 0 | 1 | 1 | 0 | ? | 1 | 1 | 1 | 0 | 1 | 0 | 0 | 0 | 0 | 1 | 0 | 0 | 0 | 0 |
| **[75]** Prosternum shape. (0) rounded between ventral and posterior surface or surfaces not distinct; (1) angulate or carinate at the limit between ventral and posterior surface; (2) with vertical lamina at limit between ventral and posterior surface | 0 | 1 | 0 | 0 | 0 | 0 | 1 | 1 | 1 | 1 | 1 | 1 | 1 | 1 | 1 | 1 | 1 | 1 | 1 | 1 | 1 | 1 | 1 | 1 | 0 | 0 | 1 | 1 | 1 | 1 | 1 | 1 | 1 | 1 | 1 | 1 | 2 | 2 | 2 | 2 |
| **[76]** Separation between anterior and posterior surface of prosternum. (0) absent or if present straight line; (1) median process, dentiform or lamelliform | 0 | 0 | 0 | 0 | 0 | 0 | 0 | 0 | 0 | 0 | 0 | 0 | 0 | 0 | 0 | 0 | 0 | 0 | 0 | 0 | 0 | ? | 0 | 0 | 0 | 0 | 0 | 0 | 0 | 0 | 0 | 0 | 0 | 0 | 0 | 0 | 1 | 1 | 1 | 1 |
| **[77]** Profurcal pit. (0) present; (1) absent | ? | ? | 0 | ? | 0 | ? | 0 | ? | 0 | ? | 0 | ? | 0 | ? | 1 | 1 | 1 | 1 | 1 | ? | 0 | ? | 0 | ? | 1 | ? | ? | ? | ? | 0 | ? | 0 | ? | ? | ? | ? | ? | ? | ? | 0 |
| **[78]** Lobes on margin of dorsal plate of profurca. (0) absent; (1) two lobes/projections present | ? | ? | 0 | ? | 0 | ? | 0 | ? | 0 | ? | 0 | ? | 0 | ? | 0 | 0 | 1 | ? | ? | ? | 0 | ? | 0 | ? | ? | ? | ? | ? | ? | 0 | ? | 0 | ? | ? | ? | ? | ? | ? | ? | 0 |
| **[79]** Ventral plate of profurca. (0) one surface visible; (1) two surfaces visible, forming a right angle together | ? | ? | 0 | ? | 0 | ? | 0 | ? | 0 | ? | 0 | ? | 0 | ? | 0 | 0 | 0 | ? | 0 | ? | 0 | ? | 0 | ? | ? | ? | ? | ? | ? | 1 | ? | 1 | ? | ? | ? | ? | ? | ? | ? | 1 |
| **[80]** Apical stripe of prosternum. (0) apparent not sunken within body; (1) internal, sunken within body | ? | 0 | 0 | 1 | 1 | ? | 1 | 1 | 1 | ? | 1 | ? | 1 | ? | 1 | 1 | 1 | ? | 1 | ? | 1 | ? | 1 | ? | ? | ? | ? | ? | ? | 1 | ? | 1 | ? | ? | ? | ? | ? | ? | ? | 1 |
| **[81]** Ornamentation of ventral belt of prepectus. (0) medioventral areola present only; (1) small and sharp medioventral tooth; (2) large medioventral tooth | 0 | 0 | 0 | 0 | 0 | 0 | 0 | 0 | 0 | 0 | 0 | 0 | 0 | 0 | 0 | 0 | 0 | 0 | 0 | 0 | 2 | 2 | 0 | 0 | 0 | 0 | 0 | 0 | 1 | 1 | 1 | 1 | 1 | 2 | 2 | 0 | 0 | 0 | 0 | 0 |
| **[82]** Projection of mesothoracic spiracle. (0) not projecting, but visible externally; (1) hidden externally; (2) partly and hardly visible as hidden by a patch of hairs on posterior margin of pronotum | 1 | 1 | 1 | 0 | 0 | 0 | 1 | 1 | 1 | 1 | 1 | 1 | 1 | 1 | 2 | 2 | 2 | 2 | 2 | 2 | 1 | 1 | 1 | 1 | 1 | 1 | 1 | 1 | 1 | 1 | 1 | 1 | 1 | 1 | 1 | 1 | 1 | 1 | 1 | 1 |
| **[83]** Structure of lateral panel of prepectus. (0) without fovea or raised rim; (1) medially foveate, with posterior and/or dorsal rim and small anterodorsal projection | 0 | 0 | 0 | 0 | 0 | 0 | 0 | 0 | 0 | 0 | 0 | 0 | 0 | 0 | 0 | 0 | 0 | 0 | 0 | 0 | 1 | 1 | 0 | 0 | - | - | 0 | 0 | 0 | 0 | 0 | 0 | 0 | 0 | 0 | 0 | 0 | 0 | 0 | 0 |
| **[84]** Size and shape of exposed lateral panel of prepectus. (0) as tall or taller than long, more than half tegula length; (1) longer than tall, more than half tegula length; (2) small, less than half tegula length; (3) lateral panel not apparent | 0 | 2 | 1 | 2 | 2 | 2 | 2 | 2 | 2 | 2 | 2 | 2 | 2 | 2 | 3 | 2 | 2 | 2 | 2 | 2 | 1 | 1 | 2 | 2 | 3 | 3 | 2 | 2 | 2 | 2 | 2 | 2 | 2 | 1 | 1 | 2 | 2 | 2 | 2 | 2 |
| **[85]** Prepectus relationship to tegula. (0) prepectus reaching tegula; (1) prepectus not reaching tegula | 0 | 0 | 0 | 1 | 1 | 1 | 0 | 0 | 0 | 0 | 0 | 0 | 0 | 0 | 1 | 1 | 0 | 0 | 0 | 0 | 0 | 0 | 0 | 0 | 1 | 1 | 0 | 0 | 0 | 0 | 0 | 0 | 0 | 0 | 0 | 0 | 0 | 0 | 0 | 0 |
| **[86]** Setation of lateral panel of prepectus. (0) setose; (1) bare | 1 | 1 | 1 | 1 | 1 | 1 | 0 | 0 | 0 | 0 | 0 | 0 | 0 | 0 | 1 | 1 | 1 | 1 | 1 | 1 | 1 | 1 | 1 | 1 | - | - | 0 | 1 | 0 | 0 | 0 | 0 | 0 | 1 | 1 | 0 | 0 | 0 | 0 | 0 |
| **[87]** Posteroventral margin of prepectus. (0) ventral margin partially or completely fused medially with episternum; (1) completely separated from mesepisternum | 0 | 0 | 0 | 0 | 0 | 0 | 1 | 1 | 1 | 1 | 0 | 0 | 0 | 0 | 0 | 0 | 0 | 0 | 0 | ? | 1 | ? | 1 | 1 | - | - | 1 | 1 | 1 | 1 | 1 | 1 | 1 | 1 | 1 | 1 | 1 | 1 | 1 | 1 |
| **[88]** Mesepisternum: epicnemium. (0) absent; (1) present, and completely delimited by carina | 0 | 0 | 0 | 1 | 1 | 1 | 1 | 1 | 1 | 1 | 1 | 1 | 1 | 1 | 0 | 0 | 0 | 0 | 0 | 0 | 1 | 1 | 1 | 1 | 0 | 0 | 0 | 1 | 1 | 1 | 1 | 1 | 1 | 1 | 1 | 1 | 1 | 1 | 1 | 1 |
| **[89]** Mesothoracic discrimen. (0) sulcate or foveate; (1) raised carina or bump anteriorly, foveate groove posteriorly; (2) raised carina overall; (3) as anchor-like ornamentation with median carin | 0 | 0 | 0 | 3 | 3 | 3 | 1 | 1 | 1 | 1 | 1 | 1 | 1 | 1 | 0 | 0 | 0 | 0 | 0 | 0 | 0 | 0 | 2 | 2 | 0 | 0 | 0 | 0 | 1 | 1 | 1 | 1 | 1 | 1 | 1 | 1 | 1 | 1 | 1 | 1 |
| **[90]** Position of mesofurcal pit. (0) adjacent to mesocoxal cavity; (1) on mesotrochantinal plate | 0 | 0 | 0 | 0 | 0 | 0 | 0 | 0 | 0 | 0 | 0 | 0 | 0 | 0 | 0 | 0 | 0 | 0 | 0 | 0 | 0 | ? | 0 | 0 | 1 | 1 | 1 | ? | 1 | 1 | 1 | 1 | 1 | 1 | 1 | 1 | 1 | 1 | 1 | 1 |
| **[91]** Shape of metepimeron. (0) subtriangular; (1) broadly rectangular or squared | 0 | 0 | 0 | 2 | 2 | 2 | 2 | 0 | 2 | 2 | 2 | 2 | 2 | 2 | 2 | 2 | 2 | 2 | 2 | 2 | 2 | 2 | 2 | 2 | 2 | 2 | 2 | 2 | 2 | 2 | 2 | 2 | 2 | 2 | 2 | 2 | 2 | 2 | 2 | 2 |
| **[92]** Number of metafurcal pits. (0) lateral (paired) pits; (1) single median pit; (2) pits absent | 0 | 0 | 0 | ? | 1 | ? | 2 | ? | 2 | ? | 2 | ? | 2 | ? | 2 | 2 | 2 | ? | 2 | ? | 2 | ? | 2 | ? | 2 | ? | 2 | ? | 2 | 2 | ? | 2 | ? | 2 | ? | 2 | ? | 2 | ? | 2 |
| **[93]** Inner lamella of metadiscrimen. (0) As usual, not especially raised; (1) strongly raised | 0 | 0 | 0 | ? | 1 | ? | 1 | ? | 1 | ? | 1 | ? | 1 | ? | 1 | 1 | 1 | ? | 1 | ? | 1 | ? | 1 | ? | ? | ? | ? | ? | ? | 1 | ? | 1 | ? | ? | ? | ? | ? | 1 | ? | 1 |
| **[94]** Metepisternal ventral shelf. (0) absent; (1) present above mid coxae, short; (2) present above mid coxae, long | 1 | 0 | 1 | 1 | 1 | 1 | 1 | 1 | 1 | 1 | 1 | 1 | 1 | 1 | 1 | 1 | 1 | 1 | 1 | 1 | 1 | 1 | 2 | 2 | 0 | ? | 1 | 1 | 1 | 1 | 1 | 1 | 1 | 1 | 1 | 1 | 1 | 1 | 1 | 1 |
| **[95]** Submedian teeth at posteroventral edge of metepisternal shelf. (0) absent; (1) present | 0 | 0 | 0 | 0 | 0 | 0 | 0 | 0 | 0 | 0 | 0 | 0 | 0 | 0 | 0 | 0 | 0 | 0 | 0 | 0 | 1 | 1 | 1 | 1 | 0 | ? | 0 | 0 | 0 | 0 | 0 | 0 | 0 | 0 | 0 | 0 | 0 | 0 | 0 | 0 |
| **[96]** Ornamentation between metacoxae. (0) median groove; (1) single median carina present; (2) two submedian carinae present, converging posteriorly; (3) two submedian carinae present, parallel and short; (4) absent | 4 | 4 | 4 | 3 | 3 | 3 | 1 | 1 | 1 | 1 | 2 | 2 | 2 | ? | 1 | 1 | 1 | 1 | 1 | 1 | 1 | 1 | 1 | 1 | 0 | ? | 0 | 0 | 0 | 0 | 0 | 0 | 0 | 0 | ? | 0 | 0 | 0 | 0 | 0 |
| **[97]** Carina connecting hind coxal and propodeal foramina. (0) absent; (1) present | ? | 1 | ? | 1 | 1 | 1 | 1 | 1 | 1 | 1 | 1 | 1 | 1 | 1 | 1 | 1 | 1 | 1 | 1 | 1 | 0 | 0 | 1 | 1 | 0 | ? | 1 | 1 | 1 | 1 | 1 | 1 | 1 | 1 | ? | 0 | 1 | 1 | 1 | 1 |
| **[98]** Number of setae on humeral plate. (0) more than four; (1) up to four | ? | 0 | 0 | 0 | 0 | 0 | 0 | 0 | 0 | 0 | 0 | 0 | 0 | 0 | 0 | 0 | 0 | 0 | 0 | 0 | ? | 1 | 1 | 1 | 1 | 1 | 1 | 1 | 1 | 1 | 1 | 1 | 1 | 1 | 1 | 1 | 1 | 1 | 1 | 1 |
| **[99]** Basal posterior lobe of fore wing. (0) absent; (1) present | 0 | 0 | 0 | 1 | 1 | 1 | 0 | 0 | 0 | 0 | 0 | 0 | 0 | 0 | 0 | 0 | 0 | 0 | 0 | 0 | 0 | 0 | 0 | 0 | 0 | 0 | 0 | 0 | 0 | 0 | 0 | 0 | 0 | 0 | 0 | 0 | ? | 0 | 0 | 0 |
| **[100]** Apicoventral tuft of setae on costal cell. (0) absent; (1) present | 0 | 0 | 0 | 1 | 1 | 1 | 0 | 0 | 0 | 0 | 0 | 0 | 0 | 0 | 0 | 0 | 0 | 0 | 0 | 0 | 0 | 0 | 0 | 0 | 0 | 0 | 0 | 0 | 0 | 0 | 0 | 0 | 0 | 0 | 0 | 0 | 0 | 0 | 0 | 0 |

| Characters | Tetramesa sp. | Rileya JRAS07552 | Macrorileya inopinata | Acanthochalcis nigricans | Cratocentrus aff decoratus | Cratocentrus GDEL00534 | Trigonura sp. | Trigonura GDEL00437 | Stypiura sp. | Stypiura GDEL00580 | Brachymeria minuta | Brachymeria podagrica | Brachymeria tibialis | Brachymeria GDEL00453 | Chalcis myrifex | Melanosmicra variventris | Conura immaculata | Conura marcosensis | Conura decisa | Conura aff xanthostigma | Dirhinus anthracia | Hontalia aff. caerulea | Epitranus observator | Epitranus GDEL00570 | Smicromorpha doddi | Smicromorpha malaica | Belaspidia nigra | Tropimeris GDEL00333 | Neochalcis fertoni | Hockeria bifasciata | Haltichella GDEL00144 | Antrocephalus sp. | Antrocephalus GDEL00481 | Notaspidium sp. | Notaspidium GDEL00469 | Zavoya cooperi | Irichohalticella sp | Lasiochalcidia dargelasii | Psilochalcis immaculata | Invreia subaenea |
|---|---|---|---|---|---|---|---|---|---|---|---|---|---|---|---|---|---|---|---|---|---|---|---|---|---|---|---|---|---|---|---|---|---|---|---|---|---|---|---|---|
| **[101]** Hyaline break on parastigma. (0) present; (1) absent | 0 | 1 | 0 | 1 | 1 | 1 | 1 | 1 | 1 | 1 | 1 | 1 | 1 | 1 | 1 | 1 | 0 | 0 | 0 | 0 | 0 | 0 | 1 | 1 | 1 | 1 | 0 | 1 | 1 | 1 | 1 | 1 | 1 | 1 | 1 | 1 | 1 | 1 | 1 | 1 |
| **[102]** Position of marginal vein relatively to front margin of wing. (0) along margin; (1) somewhat removed from margin | 0 | 0 | 0 | 0 | 0 | 0 | 0 | 0 | 0 | 0 | 0 | 0 | 0 | 0 | 0 | 0 | 0 | 0 | 0 | 0 | 0 | 0 | 0 | 0 | 0 | 0 | 0 | 0 | 0 | 0 | 0 | 0 | 0 | 1 | 1 | 0 | 1 | 1 | 1 | 1 |
| **[103]** Length of marginal vein of fore wing. (0) between 1-3 times stigmal vein + stigma length; (1) more than 3 times stigmal vein + stigma length; (2) more than 10 times length of stigmal vein plus stigma | 0 | 0 | 0 | 0 | 0 | 0 | 1 | 1 | 1 | 1 | 1 | 1 | 1 | 1 | 0 | 0 | 0 | 0 | 0 | 0 | 2 | 2 | 2 | 2 | 0 | 0 | 0 | 0 | 0 | 0 | 0 | 0 | 0 | - | - | 1 | 1 | 1 | 1 | 1 |
| **[104]** Length of postmarginal vein of fore wing (fw). (0) longer than stigmal vein + stigma, shorter than costal cell; (1) 1-2 times as long as the stigmal vein; (2) absent or shorter than stigmal vein + stigma | 0 | 0 | 0 | 0 | 0 | 0 | 1 | 1 | 1 | 1 | 1 | 1 | 1 | 1 | 0 | 0 | 0 | 0 | 0 | 0 | 2 | 2 | 2 | 2 | 2 | 2 | 0 | 2 | 0 | 0 | 0 | 0 | 0 | 2 | 2 | 2 | 2 | 2 | 2 | 2 |
| **[105]** Uncus of stigmal vein of fore wing (0) present and projecting as a linear process; (1) absent | 0 | 0 | 0 | 1 | 1 | 1 | 0 | 0 | 0 | 0 | 0 | 0 | 0 | 0 | 0 | 0 | 0 | 0 | 0 | 0 | 1 | 1 | 1 | 1 | 1 | 1 | 0 | 0 | 0 | 0 | 0 | 0 | 0 | 1 | 1 | 1 | 1 | 1 | 1 | 1 |
| **[106]** Arrangement of uncal sensilla. (0) arranged in a line; (1); grouped in a single cluster; (2) 2 pairs separated by a short space. | 1 | ? | 0 | ? | ? | ? | 0 | 0 | 0 | 0 | 0 | 0 | 0 | 0 | 0 | 0 | 0 | 0 | 0 | 0 | 0 | 0 | 0 | 0 | 0 | 0 | 0 | 0 | 0 | 0 | 0 | 0 | 0 | 0 | 0 | - | 0 | 0 | 0 | 0 |
| **[107]** Location of basal hamulus. (0) near the others; (1) distant from the others | 0 | 0 | 0 | 0 | 0 | 0 | 0 | 0 | 0 | 0 | 1 | 1 | 1 | 1 | 0 | 0 | 0 | 0 | 0 | 0 | 0 | 0 | 0 | 0 | 0 | 0 | 0 | 0 | 0 | 0 | 0 | 0 | 0 | ? | 0 | 0 | ? | 0 | 0 | 0 |
| **[108]** Shape of first hamulus. (0) curved towards wing surface, like the other hamuli; (1) straight or only slightly curved, with others strongly curved towards wing surface; (2) curved towards base of hind wing, with others curved towards wing surface | ? | ? | 1 | 0 | 0 | 0 | 1 | 1 | 1 | 1 | 1 | 1 | 1 | 1 | 0 | 0 | 0 | 0 | 0 | 0 | 0 | 0 | 1 | 1 | 1 | 1 | 1 | 1 | 1 | 1 | 1 | 1 | 1 | 2 | 2 | 0 | 2 | 2 | 2 | 2 |
| **[109]** Line of setation on posterior surface of procoxa. (0) absent; (1) present | 0 | 0 | 0 | 0 | 0 | 0 | 0 | 0 | 0 | 0 | 0 | 0 | 0 | 0 | 1 | 1 | 1 | 1 | 1 | 1 | 0 | 0 | 0 | 0 | 0 | 0 | 0 | 0 | 0 | 0 | 0 | 0 | 0 | 0 | 0 | 0 | 0 | 0 | 0 | 0 |
| **[110]** Apical ornamentation of protibia. (0) without horizontally directed stout spur or elongation; (1) with horizontally directed socketed spur; (2) without socketed spur but distinctly expanded giving the appearance of a spur | 0 | 0 | 0 | 1 | 1 | 1 | 2 | 2 | 2 | 2 | 0 | 0 | 0 | 0 | 0 | 0 | 0 | 0 | 0 | 0 | 0 | 0 | 0 | 0 | 0 | 0 | 0 | 0 | 0 | 0 | 0 | 0 | 0 | 0 | 0 | 0 | 0 | 0 | 0 | 0 |
| **[111]** Pegs at apex of mesotibia. (0) absent; (1) present | 0 | 0 | 0 | 1 | 1 | 1 | 0 | 0 | 0 | 0 | 0 | 0 | 0 | 0 | 0 | 0 | 0 | 0 | 0 | 0 | 0 | 0 | 0 | 0 | 0 | 0 | 0 | 0 | 0 | 0 | 0 | 0 | 0 | 0 | 0 | 0 | 0 | 0 | 0 | 0 |
| **[112]** Shape of metacoxa. (0) coxa not enlarged; (1) coxa enlarged and/or elongate | 0 | 0 | 0 | 1 | 1 | 1 | 1 | 1 | 1 | 1 | 1 | 1 | 1 | 1 | 1 | 1 | 1 | 1 | 1 | 1 | 1 | 1 | 1 | 1 | 1 | 1 | 1 | 1 | 1 | 1 | 1 | 1 | 1 | 1 | 1 | 1 | 1 | 1 | 1 | 1 |
| **[113]** Shape of metafemur. (0) not enlarged (greater than or equal to 3× as long as broad); (1) enlarged (smaller than or equal to 3× as long as broad) | 0 | 0 | 0 | 1 | 1 | 1 | 1 | 1 | 1 | 1 | 1 | 1 | 1 | 1 | 1 | 1 | 1 | 1 | 1 | 1 | 1 | 1 | 1 | 1 | 1 | 1 | 1 | 1 | 1 | 1 | 1 | 1 | 1 | 1 | 1 | 1 | 1 | 1 | 1 | 1 |
| **[114]** Ventral ornamentation of metafemur. (0) without denticles or teeth ventrally; (1) with small, uniform teeth similar to blade of saw over most of length; (2) with large, regular, lobe like teeth | 0 | 0 | 0 | 2 | 2 | 2 | 2 | 2 | 2 | 2 | 2 | 2 | 2 | 2 | 2 | 2 | 2 | 2 | 2 | 2 | 1 | 1 | 2 | 2 | 1 | 1 | 1 | 1 | 1 | 1 | 1 | 1 | 1 | 1 | 1 | 1 | 1 | 1 | 1 | 1 |
| **[115]** Position of basal tooth of metafemur. (0) near base of femur; (1) at mid length of femur | - | - | - | 0 | 0 | 0 | 0 | 0 | 0 | 0 | 0 | 0 | 0 | 0 | 0 | 0 | 0 | 0 | 0 | 0 | 0 | 0 | 0 | 0 | 0 | 0 | 1 | 1 | 1 | 1 | - | - | - | 0 | 0 | 1 | 1 | 1 | 1 | 1 |
| [116] Line of stout bristles on inner surface of metafemur. (0) absent; (1) present | 0 | 0 | 0 | 0 | 0 | 0 | 0 | 0 | 0 | 0 | 0 | 0 | 0 | 0 | 1 | 1 | 1 | 1 | 1 | 1 | 0 | 0 | 0 | 0 | 0 | 0 | 0 | 0 | 0 | 0 | 0 | 0 | 0 | 0 | 0 | 0 | 0 | 0 | 0 | 0 |
| **[117]** Tarsal scrobe on apicodorsal surface of metatibia. (0) absent or short, never with tooth or protrusion above; (1) present, long, without tooth or protrusion above; (2) present, long, with a tooth or protrusion above | 0 | 0 | 0 | 1 | 1 | 1 | 1 | 1 | 1 | 1 | 0 | 0 | 0 | 0 | 0 | 0 | 0 | 0 | 0 | 0 | 1 | 1 | 2 | 2 | 0 | 0 | 0 | 0 | 0 | 0 | 0 | 0 | 0 | 0 | 0 | 0 | 0 | 0 | 0 | 0 |
| **[118]** Apex of metatibia, shape. (0) truncate at right angle; (1) diagonally truncate, posteroventral corner acute; (2) diagonally truncate, posteroventral corner elongated into spine | 0 | 0 | 0 | 2 | 2 | 2 | 2 | 2 | 2 | 2 | 1 | 1 | 1 | 1 | 2 | 2 | 2 | 2 | 2 | 2 | 2 | 2 | 2 | 2 | 2 | 2 | 0 | 0 | 0 | 0 | 0 | 0 | 0 | 0 | 0 | 0 | 0 | 0 | 0 | 0 |
| **[119]** Number of apical spurs on metatibia. (0) two spurs; (1) one spur; (2) no spur | 0 | 0 | 0 | 2 | 2 | 2 | 2 | 2 | 1 | 1 | 1 | 1 | 1 | 1 | 1 | 1 | 1 | 1 | 1 | 1 | 1 | 1 | 1 | 1 | 1 | 1 | 0 | 0 | 0 | 0 | 0 | 0 | 0 | 0 | 0 | 1 | 0 | 0 | 0 | 0 |
| **[120]** Ventral carinae of metatibia. (0) absent; (1) two ventral carinae present, one lateral and one mesal | 0 | 0 | 0 | 1 | 1 | 1 | 1 | 1 | 1 | 1 | 1 | 1 | 1 | 1 | 1 | 1 | 1 | 1 | 1 | 1 | 1 | 1 | 1 | 1 | 1 | 1 | 1 | 1 | 1 | 1 | 1 | 1 | 1 | 1 | 1 | 1 | 1 | 1 | 1 | 1 |
| **[121]** Carina on inner surface of metatibia. (0) absent; (1) present | 0 | 0 | 0 | 0 | 0 | 0 | 0 | 0 | 0 | 0 | 0 | 0 | 0 | 0 | 0 | 0 | 0 | 0 | 0 | 0 | 0 | 0 | 0 | 0 | 0 | 0 | 1 | 1 | 1 | 1 | 1 | 1 | 1 | 0 | 0 | 0 | 1 | 1 | 1 | 1 |
| **[122]** Tarsal claws. (0) simple; (1) pectinate, having 1-2 peg-like extra projection(s); (2) with basal tooth | 0 | 0 | 0 | 2 | 2 | 2 | 2 | 2 | 2 | 2 | 0 | 0 | 0 | 0 | 1 | 0 | 0 | 0 | 0 | 0 | 0 | 0 | 0 | 0 | 0 | 0 | 0 | 0 | 0 | 0 | 0 | 0 | 0 | 0 | 0 | 0 | ? | 0 | 0 | 0 |
| **[123]** Spatulate seta on hind tarsal claw. (0) absent; (1) present | 0 | 0 | 0 | 0 | 0 | 0 | 0 | 0 | 0 | 0 | 1 | 1 | 1 | 1 | 0 | 0 | 0 | 0 | 0 | 0 | 0 | 0 | 0 | 0 | 0 | 0 | 0 | 0 | 0 | 0 | 0 | 0 | 0 | 0 | 0 | 0 | 0 | 0 | 0 | 0 |
| **[124]** Basal lamina on petiole. (0) absent; (1) present | 0 | 0 | 0 | 0 | 0 | 0 | 0 | 0 | 0 | 0 | 0 | 0 | 0 | 0 | 1 | 1 | 1 | 1 | 1 | 1 | 0 | 0 | 0 | 0 | 1 | 1 | 0 | 0 | 0 | 0 | 0 | 0 | 0 | 0 | 0 | 0 | 0 | 0 | 0 | 0 |
| **[125]** Insertion of petiole on propodeum. (0) at apex near metacoxal foramina; (1) at base of propodeum, near posterior margin of metanotum | 0 | 0 | 0 | 0 | 0 | 0 | 0 | 0 | 0 | 0 | 0 | 0 | 0 | 0 | 0 | 0 | 0 | 0 | 0 | 0 | 0 | 0 | 0 | 0 | 1 | 1 | 0 | 0 | 0 | 0 | 0 | 0 | 0 | 0 | 0 | 0 | 0 | 0 | 0 | 0 |
| **[126]** Relationship between petiole and propodeum. (0) body of petiole entering propodeum; (1) petiole with complete lamina surrounding petiolar foramen of propodeum; (2) petiole with ventral lamina abutting against petiolar foramen; (3) kneecap only entering propodeum | 0 | 0 | 0 | 0 | 0 | 0 | 0 | 0 | 0 | 0 | 0 | 0 | 0 | 0 | 3 | 3 | 3 | 3 | 3 | 3 | 2 | 2 | 3 | 3 | 3 | 3 | 1 | 1 | 1 | 1 | 1 | 1 | 1 | 1 | 1 | 1 | 1 | 1 | 1 | 1 |

| Characters | *Tetramesa sp.* | *Rileya JRAS07552* | *Macrorileya inopinata* | *Acanthochalcis nigricans* | *Cratocentrus aff decoratus* | *Cratocentrus GDEL00534* | *Trigonura sp.* | *Trigonura GDEL00437* | *Stypiura sp.* | *Stypiura GDEL00580* | *Brachymeria minuta* | *Brachymeria podagrica* | *Brachymeria tibialis* | *Brachymeria GDEL00453* | *Chalcis myrifex* | *Melanosmicra variventris* | *Conura immaculata* | *Conura marcosensis* | *Conura decisa* | *Conura aff xanthostigma* | *Dirhinus anthracia* | *Hontalia aff. caerulea* | *Epitranus observator* | *Epitranus GDEL00570* | *Smicromorpha doddi* | *Smicromorpha malaica* | *Belaspidia nigra* | *Tropimeris GDEL00333* | *Neochalcis fertoni* | *Hockeria bifasciata* | *Haltichella GDEL00144* | *Antrocephalus sp.* | *Antrocephalus GDEL00481* | *Notaspidium sp.* | *Notaspidium GDEL00469* | *Zavoya cooperi* | *Irichohalticella sp* | *Lasiochalcidia dargelasii* | *Psilochalcis immaculata* | *Invreia subaenea* |
|---|---|---|---|---|---|---|---|---|---|---|---|---|---|---|---|---|---|---|---|---|---|---|---|---|---|---|---|---|---|---|---|---|---|---|---|---|---|---|---|---|
| **[127]** Fusion of petiole with first gastral sternite in females. (0) not fused; (1) fused | 1 | 1 | 1 | 0 | 0 | 0 | 0 | 0 | 0 | 0 | 0 | 0 | 0 | 0 | 0 | 0 | 0 | 0 | 0 | 0 | 1 | 1 | 0 | 0 | 0 | 0 | 0 | 0 | 0 | 0 | 0 | 0 | 0 | 0 | 0 | 0 | 0 | 0 | 0 | 0 |
| **[128]** Transverse carina in front of cercal plates. (0) absent; (1) present, cerci inserted in foveae situated just behind the carina; (2) present, cerci situated posteriorly to carina | 0 | 1 | 1 | 2 | 2 | 2 | 0 | 0 | 0 | 0 | 1 | 1 | 1 | 1 | 0 | 0 | 0 | 0 | 0 | 0 | 0 | 0 | 0 | 0 | 0 | 0 | 1 | 1 | 1 | 1 | 1 | 1 | 1 | 1 | 1 | 1 | 1 | 1 | 1 | 1 |
| **[129]** Relative placement of hypopygium tip. (0) in basal half; (1) near tip of gaster; (2) somewhat beyond tip of gaster, apex densely setose | 0 | 0 | 0 | 0 | 0 | 0 | 0 | 0 | 1 | 1 | 1 | 1 | 1 | 1 | 2 | 2 | 0 | 0 | 0 | 0 | 0 | 0 | 0 | 0 | 0 | 0 | 0 | 0 | 0 | 0 | 0 | 0 | 0 | 1 | 1 | 0 | 0 | 0 | 0 | 0 |
| **[130]** Orientation of ovipositor sheaths. (0) straight; (1) curved downwards | 0 | 0 | 0 | 0 | 0 | 0 | 0 | 0 | 1 | 1 | 0 | 0 | 0 | 0 | 0 | 0 | 0 | 0 | 0 | 0 | 0 | 0 | 0 | 0 | 0 | 0 | 0 | 0 | 0 | 0 | 0 | 0 | 0 | 0 | 0 | 0 | 0 | 0 | 0 | 0 |
