## AppendixS4 for "Ultra-Conserved Elements and morphology reciprocally illuminate conflicting phylogenetic hypotheses in Chalcididae (Hymenoptera, Chalcidoidea)"

### Appendix S4. The tentorium and its external landmarks in the Chalcididae

This study is the first in the whole superfamily Chalcidoidea to investigate the tentorium as a phylogenetic character and to establish the connection between the inner skeleton of the cephalic capsule and its external landmarks on the back of the head. In this section, details are provided on the methodology used by GD to examine and code the different bridges.

#### S4.1. Context

Phylogenetic informativeness of the characters of the head capsule in Hymenoptera was recently highlighted (Vilhelmsen 2011; Burks & Heraty 2015; Zimmermann & Vilhelmsen 2016). However, interpretation is difficult and requires landmarks (Burks & Heraty, 2015). More precisely, the identity of the sclerotized structures between the occipital foramen and the oral fossa are still debated. Homology and nomenclature of these structures were established by Snodgrass (1928, 1942 and 1960) and reassessed by Vilhelmsen (1999) and Burks & Heraty (2015). These authors describe various types of 'bridges', such as postgenal, hypostomal and subforaminal bridges, according to the cephalic part – postgena or hypostoma – from which they putatively originate.

In his phylogenetic analyses of the Chalcididae, Wijesekara (1997a & 1997b) used the back of the head – reduced to a single character – and distinguished an 'hypostomal bridge' and a 'genal bridge'. The detailed examination of the back of the head in the Eurytomidae (Lotfalizadeh et al. 2007), probable sister group of the Chalcididae, provided useful characters for their phylogeny and prompted GD to also investigate these characters in the Chalcididae.

Chalcididae exhibit variable and puzzling structures that may be phylogenetically informative but request a thorough identification of homologies among the subfamilies and more largely with other families of Chalcidoidea. Examination of the tentorium appeared the unique way to provide landmarks and additional characters.

#### S4.2. Material and methods

***Sampling***. At least one specimen of each subfamily and tribe was used for the examination of the tentorium. The outgroup includes *Leucospis dorsigera* Illiger (Leucospidae), *Macrorileya inopinata* (Silvestri) (Eurytomidae, Buresiinae), *Tetramesa* sp., *Eurytoma crotalariae* Risbec and *Aximopsis collina* (Zerova) (Eurytomidae, Eurytominae), *Chalcedectus* sp. (Chalcedectini), *Cleonymus brevis* Bouček (Cleonymini) and *Lycisca* sp. (Lyciscini), all presently classified in Pteromalidae subfamily Cleonyminae, *Norbanus* sp. and *Pteromalus* sp. (Pteromalidae, Pteromalinae) and *Glyphomerus stigma* (Fabricius) (Torymidae). The ingroup included *Cratocentrus* aff. *decoratus* (Klug) (Cratocentrinae), *Stypiura* sp. and *Trigonura* sp. (Phasgonophorini), *Brachymeria minuta* (Linnaeus) and *B. tibialis* (Walker) (Brachymeriini), *Chalcis myrifex* (Sulzer), *Conura decisa* (Walker), *C. immaculata* (Cresson) and *C. femorata* (Fabricius), *Melanosmicra variventris* (Cameron) (Chalcidini), *Epitranus observator* Walker (Epitraninae), *Dirhinus anthracia* Walker (Dirhininae), *Hockeria bifasciata* Walker, *Antrocephalus* sp. from Reunion Island, *Notaspidium* sp. from Columbia and *Invreia subaenea* Masi (Haltichellinae).

***Specimen examination and imaging***. The maxilla and labium were removed and the head was fixed with water-soluble glue on a slide. A transverse section of the head was made with a razor blade. The posterior part of the head was then washed in water, cleared through immersion in potassium hydroxide at 10%, followed by washing with increasing concentrations of ethanol. The remaining tissues were removed in order to solely let the tentorium visible. The head was finally fixed at the apex of a minutien pin for examination with a stereomicroscope. The imaging was made with a JVC KY-75U 3CCD digital camera attached to an EntoVision microscope and the stacked, serial images obtained were combined using Cartograph 5.6.0 (Microvision, Evry, France) software. Finally, we used a SEM microscope (Zeiss DSM 950) for more detailed examination and further imaging.

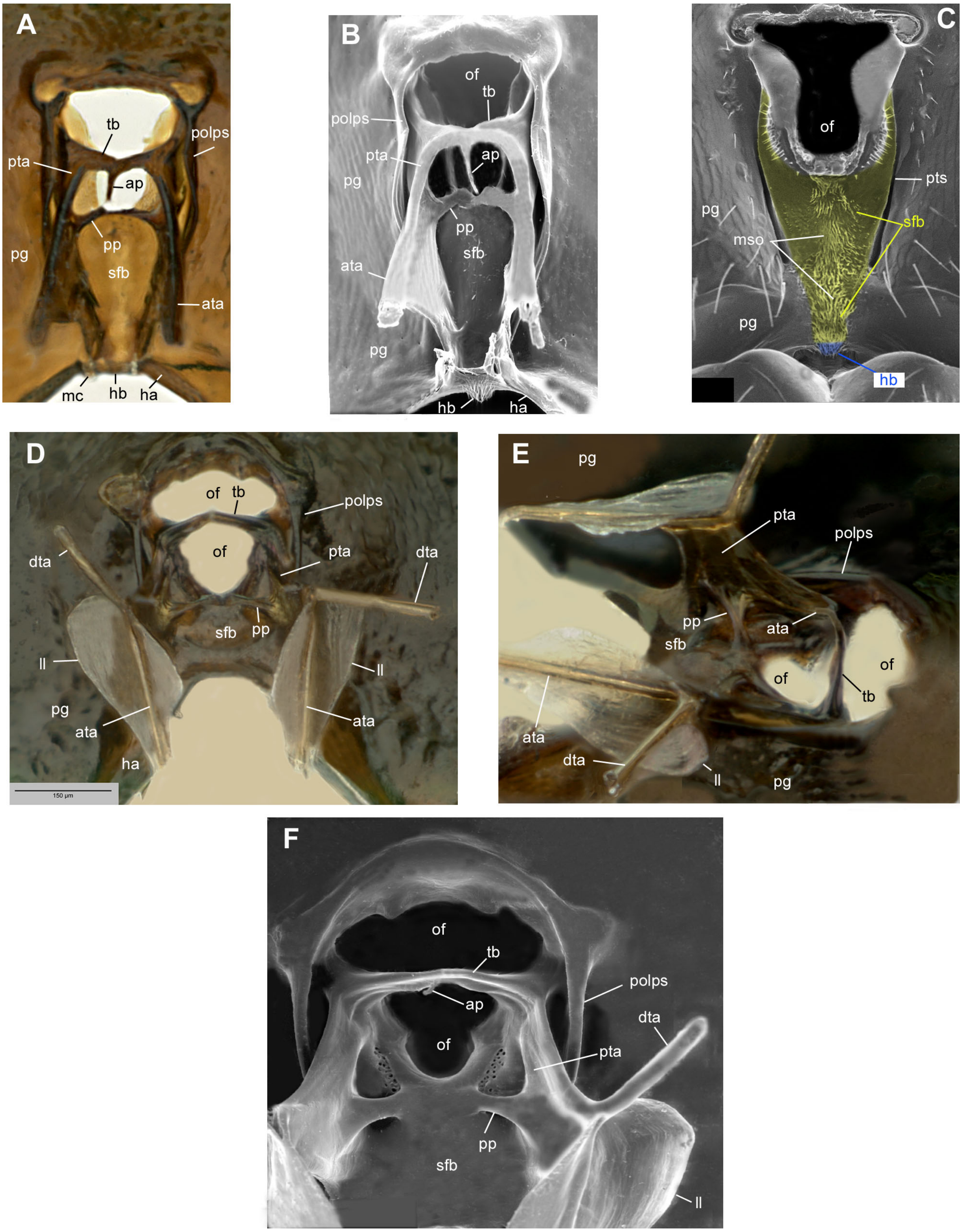


**Figure S4.1**. Tentorium (**B, D-F**) and subforaminal bridge (**A, C**) of Chalcidoidea. B, C and F, SEM Images. **A-C**, *Glyphomerus stigma* (Torymidae). **D-F**, *Chalcedectus* sp. (Chalcedectini, presently classified in Pteromalidae Cleonyminae). A, B, D, F, anterior view; E, anterolateral view. Abbreviations. ap, anterior process, ata, anterior tentorial arm, dta, dorsal tentorial arm; hb, hypostomal bridge; hc, hypostomal carina; ll, lateral lamella; mc, maxillary condyle; mso, median strip of ornamentation; of, occipital foramen; pg, postgena; pgvd, ventral depression of postgena; mc, maxillary condyle; pola, postoccipital lateral arm; polps, postoccipital lateral process; pp, posterior process; pta, posterior tentorial arm; pts, posterior tentorial sulcus; sfb, subforaminal bridge; tb, tentorial bridge.

***Nomenclature***. Zimmermann & Vilhelmsen (2016) defined and described in detail the structures of the tentorium; the nomenclature used in their paper is followed here. Examination of the tentorium provided evidence for defining the landmarks that were used to identify and name the external structures on the back of the head; here we mostly followed the nomenclature proposed by Burks & Heraty (2015) except for some errors of interpretation.

#### S4.3. Abbreviations

| | | |
|---|---|---|
| ap | anterior process | [internal] |
| ata | anterior tentorial arm | [internal] |
| atp | anterior tentorial pit | [external] |
| ca | cardo | [external] |
| dta | dorsal tentorial arm | [internal] |
| ha | hypostoma | [external] |
| hb | hypostomal bridge | [external] |
| hc | hypostomal carina | [external] |
| hp | hypostomal process | [internal] |
| hpp | pit as apex of hypostomal process | [external] |
| ll | lateral lamella of anterior tentorial arm | [internal] |
| mc | maxillary condyle | [external] |
| mso | median strip of ornamentation | [external] |
| of | occipital foramen | [external and internal] |
| pola | postoccipital lateral arm | [external] |
| pp | posterior process | [internal] |
| ppp*d* | pit at dorsal end of posterior process | [external] |
| ppp*v* | pit at ventral end of posterior process | [external] |
| pta | posterior tentorial arm | [internal] |
| ptp | posterior tentorial pit | [external] |
| pts | posterior tentorial sulcus | [external] |
| sfb | subforaminal bridge | [external and internal] |
| tb | tentorial bridge | [internal] |
| tbp | pit at lateral end of tentorial bridge | [external] |

#### S4.4. Results

##### *S4.4.1. Identification of the bridges separating occipital and oral foramen.*

Clearing the head showed that the cuticle was not uniformly thick over its posterior part. The cuticle was thinner along the median strip of ornamentation (mso) than on the postgena or the hypostoma. The relevant surface may therefore be identified as a different structure that constitutes a bridge. The lower-most bridge, situated below the dorsal level of the hypostomal carina, and receding in comparison with the surface of the postgena, is thus identified as the hypostomal bridge (hb). The surface delimited laterally by the posterior tentorial sulci (pts) is the subforaminal bridge (sfb). As the pts does not always reach ventrally the level of the upper limit of hypostoma, the transverse strip between the two surfaces may be hypothesized of postgenal origin. The median strip of ornamentation (mso) which is an extension of the subforaminal bridge (sfb), is thus formed of two parts: a broad dorsal surface delimited by pts and a narrow ventral surface reduced to the median strip of ornamentation (mso).

##### *S4.4.2. Structure of the tentorium in the outgroups*

Except pteromalines which exhibit an original structure of the tentorium and therefore of the back of the head, all other outgroups exhibit the same structure, with little variations (see *Chalcedectus* and

*Glyphomerus* in Fig. S4.1). *Inside the head capsule*. The dorsal tentorial arm (dta) abuts on the upper frons and bears moderately narrow lateral lamellae (ll). The posterior tentorial arm (pta) appears as a triangular, strongly sclerotized plate, standing at a right angle with the surface of the postgena. The posterior process (pp) originates near the base of the pta, along the inner surface of the subforaminal bridge (sfb) and reaches dorsally the ventral margin of the occipital foramen (of). The dorsal edge of the pta continues as the tentorial bridge (tb), here thick and strongly sclerotized, and forms a T-like structure with the anterior process (ap). *Outside the head capsule*. Deep posterior tentorial sulci (pts) are well visible, the hypostomal carina (hc) is moderately broken mesally, the maxillary condyles (mc) are separated with a reduced, shortly sloping and narrow hypostomal bridge (hb).

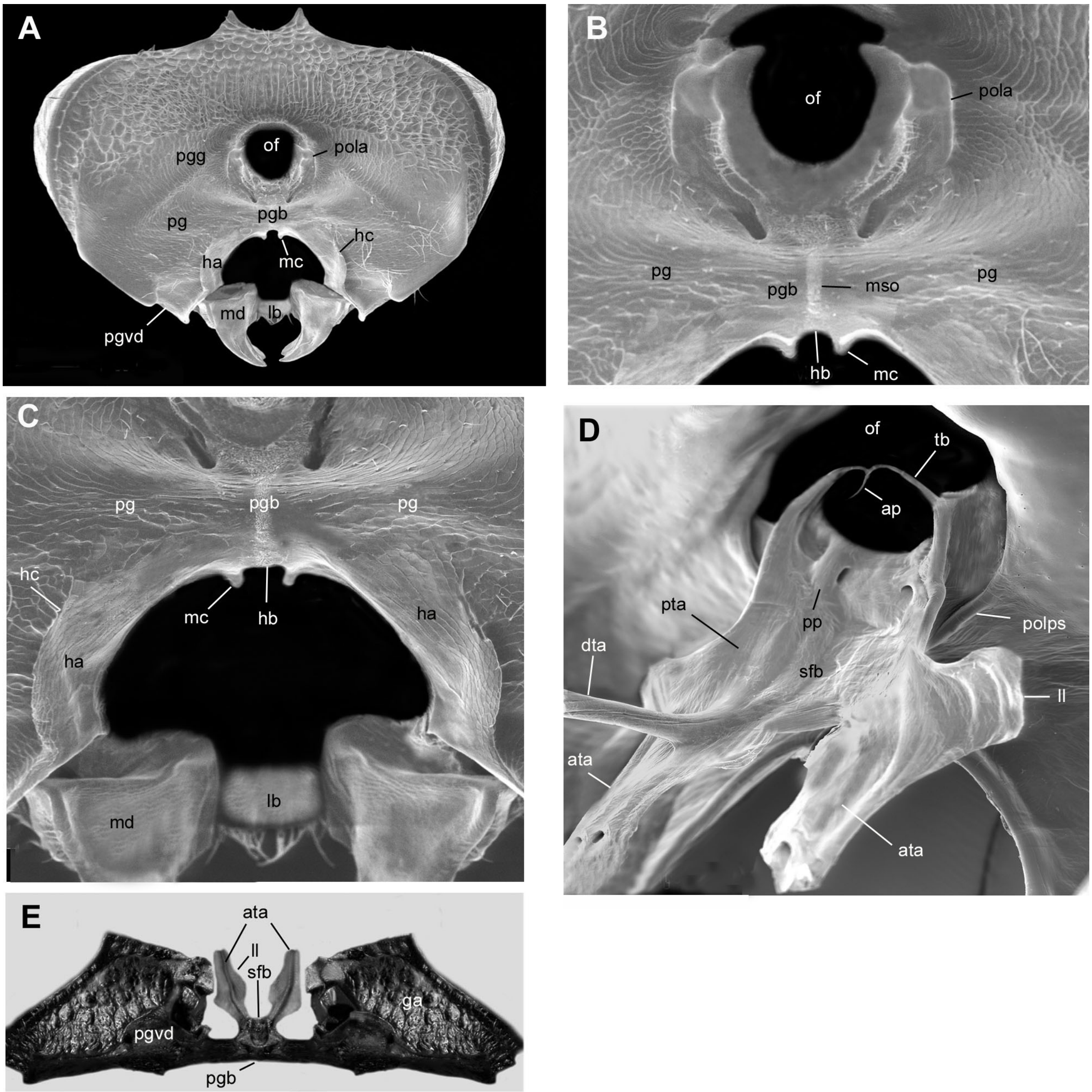


**Figure S4.2**. Tentorium (**D**) in anterodorsal view, head (**A**), postgenal bridge (**B, C**) in caudal view and postgenal bridge plus tentorium in ventral view (**E**) of *Cratocentrus* (Chalcididae: Cratocentrinae). A-D, SEM images. **Abbreviations**. ap, anterior proccess, ata, anterior tentorial arm, dta, dorsal tentorial arm; ga, gena; ha, hypostoma; hc, hypostomal carina; lb, labrum; ll, lateral lamella; md, mandible; mso, median strip of ornamentation; of, occipital foramen; pg, postgena; pgb, postgenal bridge; pgvd, ventral depression of postgena; mc, maxillary condyle; pola, postoccipital lateral arm; pp, posterior process; pta, posterior tentorial arm; sfb, subforaminal bridge; tb, tentorial bridge.

##### *S4.4.3. Structure of the tentorium in the Chalcididae*

Chalcididae differ in three respects from the structure described above: 1) the dorsal tentorial arm (dta) abuts at a much lower level on the cuticle, hardly above the antennal toruli; 2) the dta on the whole bears a wider lateral lamella, especially the outer one; 3) the arms of the tentorial bridge (tb) are thinner and form a Y-like structure with the anterior process (ap).
According to our observations, the cephalic skeleton, the tentorium and the bridges separating the occipital and the oral foramen seemed to have followed two different evolutionary pathways: one in Cratocentrinae and another in all other Chalcididae.

###### S4.4.3.1 Cratocentrinae (Fig. S4.2)

The anterior (ata) and posterior tentorial arms (pta) are broad (Fig. S4.2D) and form a strongly sclerotized strip; the lateral lamella of anterior tentorial arm (ll) is apically very broad (Fig. S4.2D); the hypostomal carina (hc) is widely broken (Fig. S4.2C), the maxillary condyles (mc) are narrowly separated and the hypostomal bridge (hb) is vestigial (Fig. S4.2B). The main difference with all other Chalcididae – that probably represents a unique case within Chalcidoidea – is the presence of two bridges. The subforaminal bridge (sfb) is not visible from outside as is mounted on a base at the surface of the postgena (Fig. S4.2E), which thus forms a true postgenal bridge. Consequently, none of the external landmarks of the tentorium are visible from outside (Fig. S4.2A). In the mesal part of the head, the postgena merges progressively into the hypostoma and the postgenal bridge merges into the hypostomal bridge (hb) without any visible limit.

###### S4.4.3.2 Other Chalcididae

The other Chalcididae differ from the Cratocentrinae by the following character states, that are best illustrated in the Phasgonophorini: 1) posterior tentorial arm (pta) not forming an uniformly sclerotized plate but appearing as a septa, re-enforced by two processes: the true pta continuing as a tentorial bridge (tb) on the dorsal edge (Fig. S4.3D), and, ventrally, the posterior process (pp) (Fig. S4.3E); 2) subforaminal bridge (sfb) forming a right to acute angle with hypostoma (ha) and hypostomal bridge (hb) (Fig. S4.3B); 3) maxillary condyles (mc) more distant from each other (Fig. S4.5B); 4) hypostomal bridge (hb) deeply sloping and longer than wide; 5) laterally, on either side of the hb, anterior tentorial arm (ata) extending into an hypostomal process (hp), visible from the outside as the hypostomal process pit (hpp); 5) narrow posterior tentorial sulcus (pts) visible, converging ventrally (Fig. S4.3C); 6) postoccipital lateral arms (pola) short (Fig. S4.4A), not extending to ventral end of pts, and strongly converging ventrally. We describe below how the different groups differ from this groundplan.

###### *S4.4.3.2.1 Haltichellinae (Fig. S4.4)*

Haltichellinae differs as follows: 1) lateral lamella (ll) of anterior tentorial arm (ata) extending on posterior tentorial arm (pta) (Fig. S4.4D, F); 2) posterior tentorial sulcus (pts) reduced to short grooves diverging ventrally (Fig. S4.4A, B, C) and linking the tentorial bridge pit (tbp) to the ventral pit of the posterior process (ppp*v*); 3) hypostomal carina (hc) continuing above and joining the postoccipital lateral arm (pola) (Fig. S4.4C); 4) hypostomal bridge (hb) quite broad and progressively sloping from subforaminal bridge (sfb) without distinct limit between them (Fig. S4.4D, F, G).

###### *S4.4.3.2.2 Brachymeriini: genus Brachymeria (Fig. S4.5)*

*Brachymeria* differs as follow: 1) posterior tentorial arm (pta) as a thick and strongly sclerotized process, which forms a very acute angle with the surface of the postgena (Fig. S4.5D); 2) posterior tentorial pit (ptp) and posterior tentorial sulcus (pts) absent (Fig. S4.5C); 3) subforaminal bridge (sfb) continuing below as a sloping and quite broad hypostomal process (hb) without carina separating the two bridges (Fig. S4.5C); 4) maxillary condyle (mc) widely separated

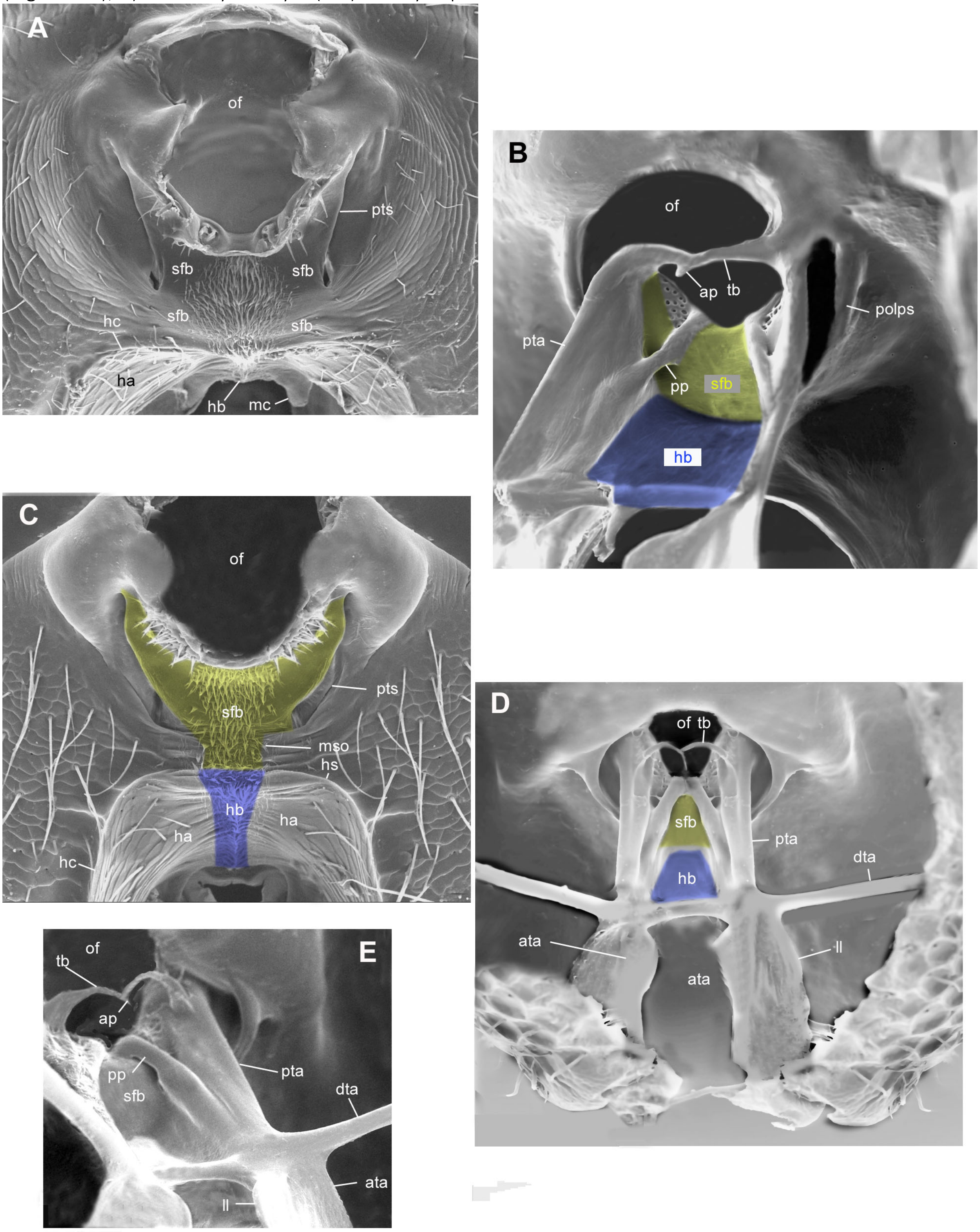


**Figure S4.3**. SEM images of tentorium (**B, D, E**) and subforaminal bridge (**A, C**) of Phasgonophorinae (Chalcididae). Subforaminal bridge shaded in yellow, hypostomal bridge in blue. **A, B**. *Trigonura* sp.; **C-E**. *Stypiura* sp. **Abbreviations**. ap, anterior proccess, ata, anterior tentorial arm, dta, dorsal tentorial arm; hb, hypostomal bridge; hc, hypostomal carina; ll, lateral lamella; mc, maxillary condyle; mso, median strip of ornamentation; of, occipital foramen; pg, postgena; pola, postoccipital lateral arm; polps, postoccipital lateral process; pp, posterior process; pta, posterior tentorial arm; pts, posterior tentorial sulcus; sfb, subforaminal bridge; tb, tentorial bridge.

(Fig. S4.5D); 5) ventral apex of posterior process (pp) not reaching the posterior tentorial arm (pta) (Fig. S4.5D, E); 6) median strip of ornamentation (mso) broad especially on hypostomal process (hb) (Fig. S4.5C); 7) tentorial bridge pit (tbp) present; 8) posterior process (ppp*v*) present; 9) large hypostomal process pit (hpp) present at lateroventral angle of subforaminal bridge (sfb) (Fig. S4.5C); 10) postoccipital lateral arms (pola) parallel to each other and smoothly continuing to the hpp ventrally.

*S4.4.3.2.3 Chalcidini: genera Chalcis and Melanosmicra (Fig. S4.6)*

The structure of the cephalic capsule of *Chalcis* is a mixture of the one described in S4.4.3.2 and in Brachymeriini. The posterior tentorial arm (pta), and posterior process (pp) are identical to those described for the Phasgonophorini (Fig. S4.6B); thin posterior tentorial sulcus (pts) are present and parallel, not converging to each other ventrally (Fig. S4.6A). The morphology of the median strip of ornamentation (mso), subforaminal bridge (sfb), hypostomal bridge (hb), maxillary condyle (mc), postoccipital lateral arm (pola) are mostly identical to that described for *Brachymeria* (Fig. S4.6C, D).

*S4.4.3.2.4 Chalcidini: genus Conura (Fig. S4.7)*

In *Conura* the structure is similar to *Brachymeria*: median strip of ornamentation (mso) (Fig. S4.7B), subforaminal bridge (sfb) (Fig. S4.7C), hypostomal process (hb) and maxillary condyle (mc) are identical (Fig. S4.7E); in addition, the posterior process (ppp*v*) and the hypostomal process (hpp) are also present although the former may be very small according to the *Conura* species. Nevertheless, the posterior tentorial arms (pta) are identical to that described in S4.4.3.2 and the postoccipital lateral arm (pola) are shorter than in *Brachymeria*. Thus, a parallel transformation occurred in the two tribes.

*S4.4.3.2.5 Epitraninae: genus Epitranus (Fig. S4.8)*

The cephalic capsule differs as follow: 1) posterior process (pp) very short, not reaching dorsally the edge of the occipital foramen (Fig. S4.8D), not joining ventrally the posterior tentorial arms (pta); 2) posterior tentorial sulcus (pts) or posterior tentorial pit (ptp) absent (Fig. S4.8A, B); 3) hypostomal process (hpp) absent; 3) pit at dorsal end of posterior process (ppp*d*) present (Fig. S4.8B); 3) median strip of ornamentation (mso) quite narrow, vestigial (Fig. S4.8C); 4) both subforaminal (sfb) and hypostomal bridge (hb) long (Fig. S4.8C); 4) postoccipital lateral arm (pola) diverging (Fig. S4.8B); 5) postgena groove present.

*S4.4.3.2.6 Dirhininae (Fig. S4.9)*

The structure of the cephalic capsule of Dirhininae is similar to that observed in Epitraninae. It differs mainly by the complete absence of a median strip of ornamentation (mso) and of a postgenal groove (Fig. S4.9B) and by the presence of an additional dorsal process on the posterior tentorial arms (pta) that originates from a furcation with the arm forming the tentorial bridge (tb).

*S4.4.3.2.7 Smicromorphinae: genus Smicromorpha*

The head was not prepared to preserve the integrity of the very few specimens in collection, but the part of the inner skeleton could be examined through the thin and translucent cuticle of the postgena. The head of *Smicromorpha* is highly modified and it was difficult to identify the homologies with the structures found in other Chalcidid subfamilies. The hypostoma is vestigial. There is an expanded brigde above the hypostoma that could either be interpreted as a subforaminal bridge or a true post genal bridge. All pits mentionned above were absent; as well as the median strip of ornamentation. Very long submedian arms parallel to the surface of the postgena [that could be either the anterior tentorial arms or the posterior tentorial arms] are present.

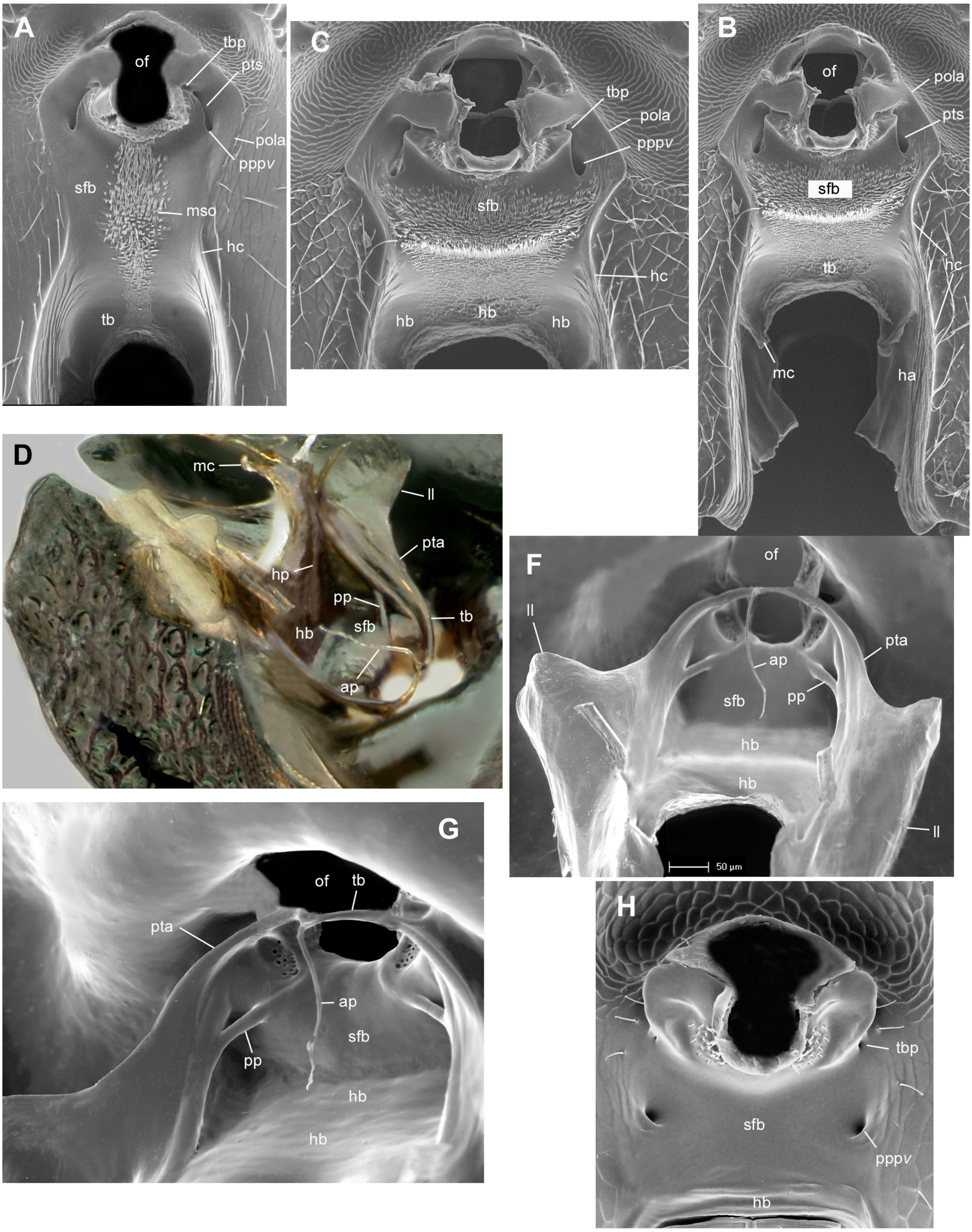


**Figure S4.4**. Tentorium (**D-G**), and subforaminal bridge (**A, C**) of Haltichellinae (Chalcididae). All are SEM images except (D). A, B, C and H in caudal view; D in posterolateral view; F and G in anterolateral view. **A**, *Hockeria bifasciata*; **B-G**, *Antrocephalus sp*., **H**, *Notaspidium giganteum*. **Abbreviations.** ap, anterior proccess, ata, anterior tentorial arm, ha, hypostoma; hb, hypostomal bridge; hc, hypostomal carina; hp, hypostomal process; ll, lateral lamella; of, occipital foramen; pg, postgena; mc, maxillary condyle; mso, median strip of ornamentation; pola, postoccipital lateral arm; pp, posterior process; pppv, pit at ventral end of posterior process; pta, posterior tentorial arm; sfb, subforaminal bridge; tb, tentorial bridge; tbp, pit at lateral end of tentorial bridge.

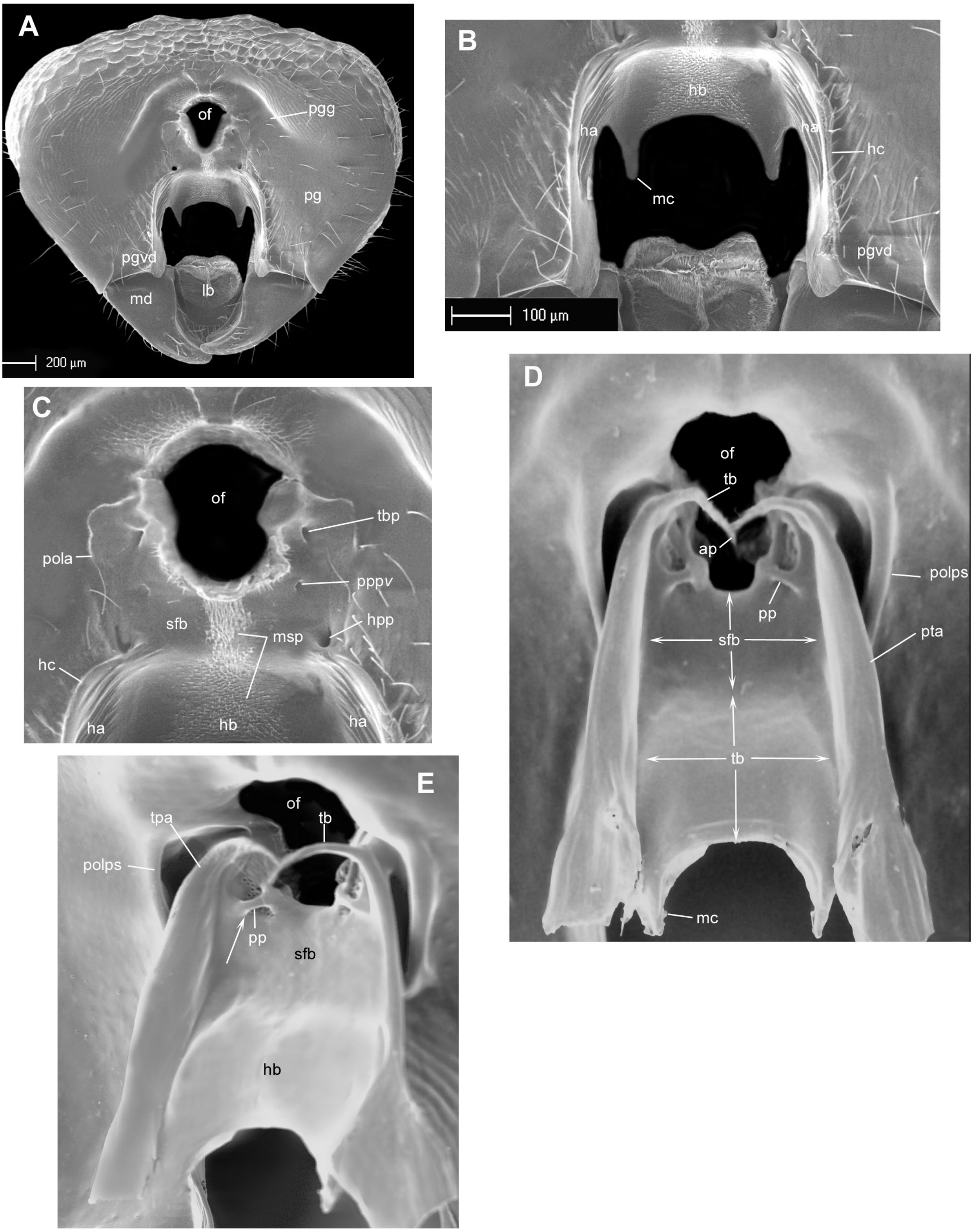


Figure S4.5. SEM images of tentorium (D-G), head (A) and subforaminal bridge (B, C) of *Brachymeria minuta* (Chalcididae: Brachymeriinae). A-C, in caudal view; E, in anterolateral viewD in anterior view. **Abbreviations**. ap, anterior proccess, ata, anterior tentorial arm, ha, hypostoma; hb, hypostomal bridge; hc, hypostomal carina; hpp, pit at dorsal end of hypostomal process; mc, maxillary condyle; md, mandible; mso, median strip of ornamentation; of, occipital foramen; pg, postgena; pgg, postgenal groove; pgvd, ventral depression of postgena; pola, postocciptal lateral arm; polps, postoccipital lateral process; pp, posterior process; pppv, pit at ventral end of posterior processs; pta, posterior tentorial arm; sfb, subforaminal bridge; tb, tentorial bridge; tbp, pit at lateral end of tentorial bridge.

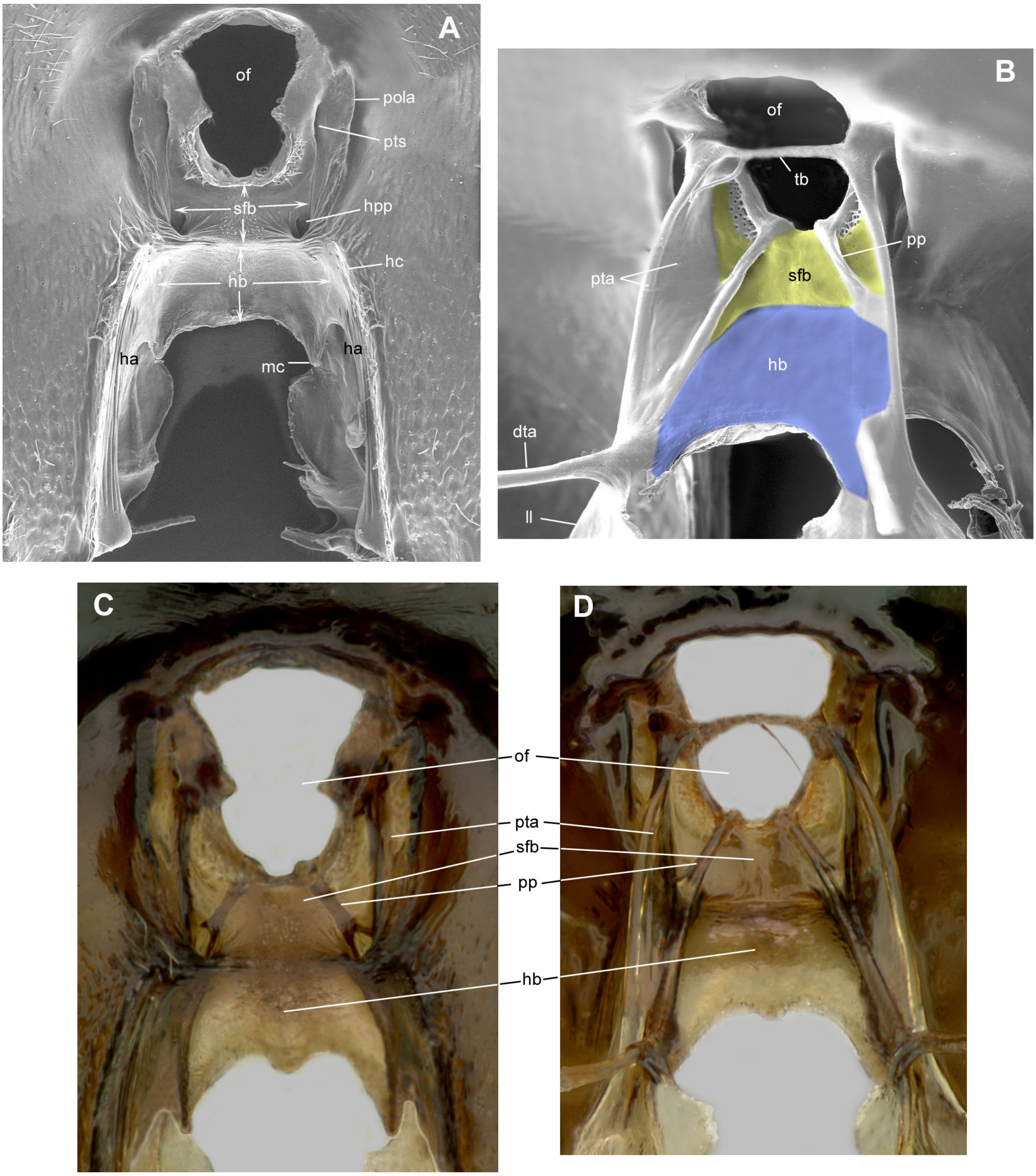


**Figure S4.6**. Tentorium (**B, D**) and subforaminal bridge (**A, C**) of *Chalcis myrifex* (Chalcididae: Chalcidinae). A and B, SEM images. A, C and D in caudal view; B in anterolateral view. Subforaminal bridge shaded in yellow, hypostomal bridge in blue. **Abbreviations**. ap, anterior proccess, dta, dorsal tentorial arm; ha, hypostoma; hb, hypostomal bridge; hc, hypostomal carina; of, occipital foramen; hpp, pit at dorsal end of hypostomal process; ll, lateral lamella; mc, maxillary condyle; pola, postocciptal lateral arm; pp, posterior process; pta, posterior tentorial arm; pts, posterior tentorial sulcus; sfb, subforaminal bridge; tb, tentorial bridge.

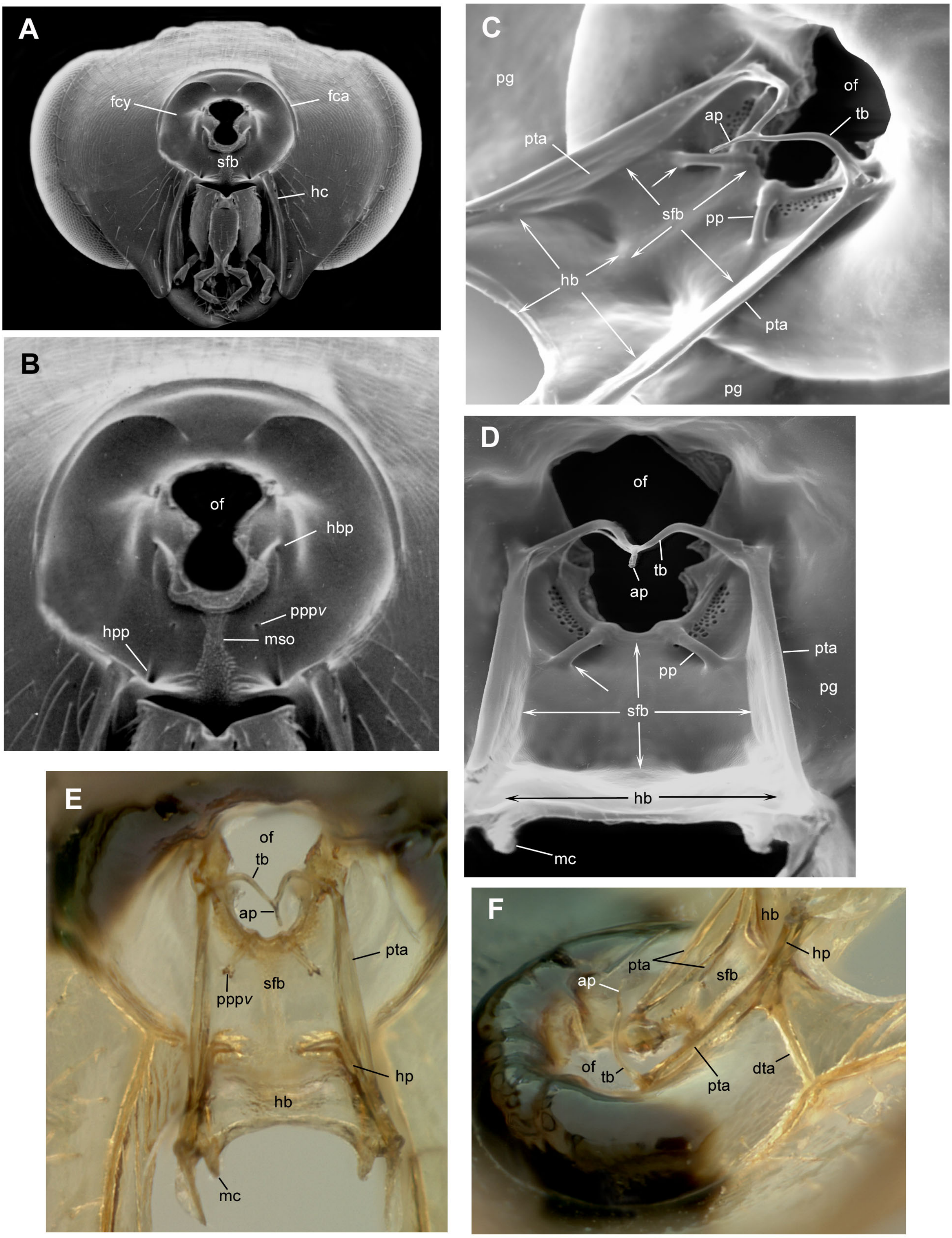


Figure S4.7. Tentorium (C-F), head (A) and subforaminal bridge (B) of *Conura decisa* (Chalcididae: Chalcidinae). A and B in caudal view; D and E in anterior view; C in anterodorsal view; F in lateral view. A-B, SEM images. **Abbreviations**. ap, anterior proccess, dta, dorsal tentorial arm; fca, foraminal carina; fcy, foraminal cavity; hb, hypostomal bridge; hc, hypostomal carina; hpp, pit at dorsal end of hypostomal process; mc, maxillary condyle; mso, median strip of ornamentation; of, occipital foramen; pg, postgena; pp, posterior process; pppv, pit at ventral end of posterior processs; pta, posterior tentorial arm; sfb, subforaminal bridge; tb, tentorial bridge; tbp, pit at lateral end of tentorial bridge.

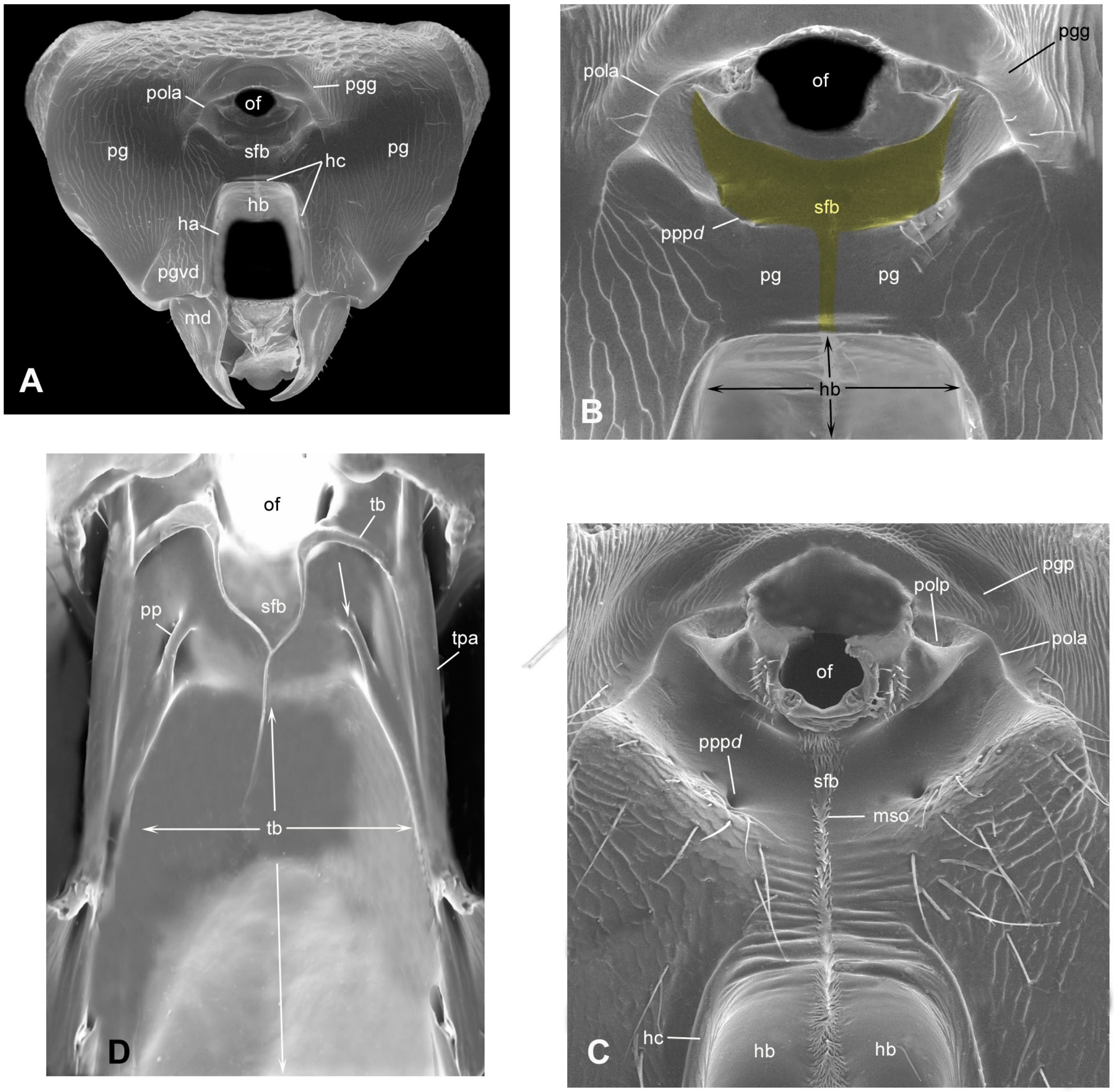


**Figure S4.8**. SEM images of tentorium (**D**), head (**A**) and subforaminal bridge (**B, C**) of Epitraninae (Chalcididae). A-C in caudal view; D in anterior view. Subforaminal bridge shaded in yellow. A and B, *Epitranus inops*; C and D, *E. observator*. **Abbreviations**. ap, anterior proccess, ha, hypostoma; hb, hypostomal bridge; hc, hypostomal carina; mc, maxillary condyle; md, mandible; mso, median strip of ornamentation; of, occipital foramen; pg, postgena; pgg, postgenal groove; pgvd, ventral depression of postgena; pola, postocciptal lateral arm; polps, postoccipital lateral pit; pp, posterior process; pppd, pit at dorsal end of posterior processs; pta, posterior tentorial arm; sfb, subforaminal bridge; tb, tentorial bridge.

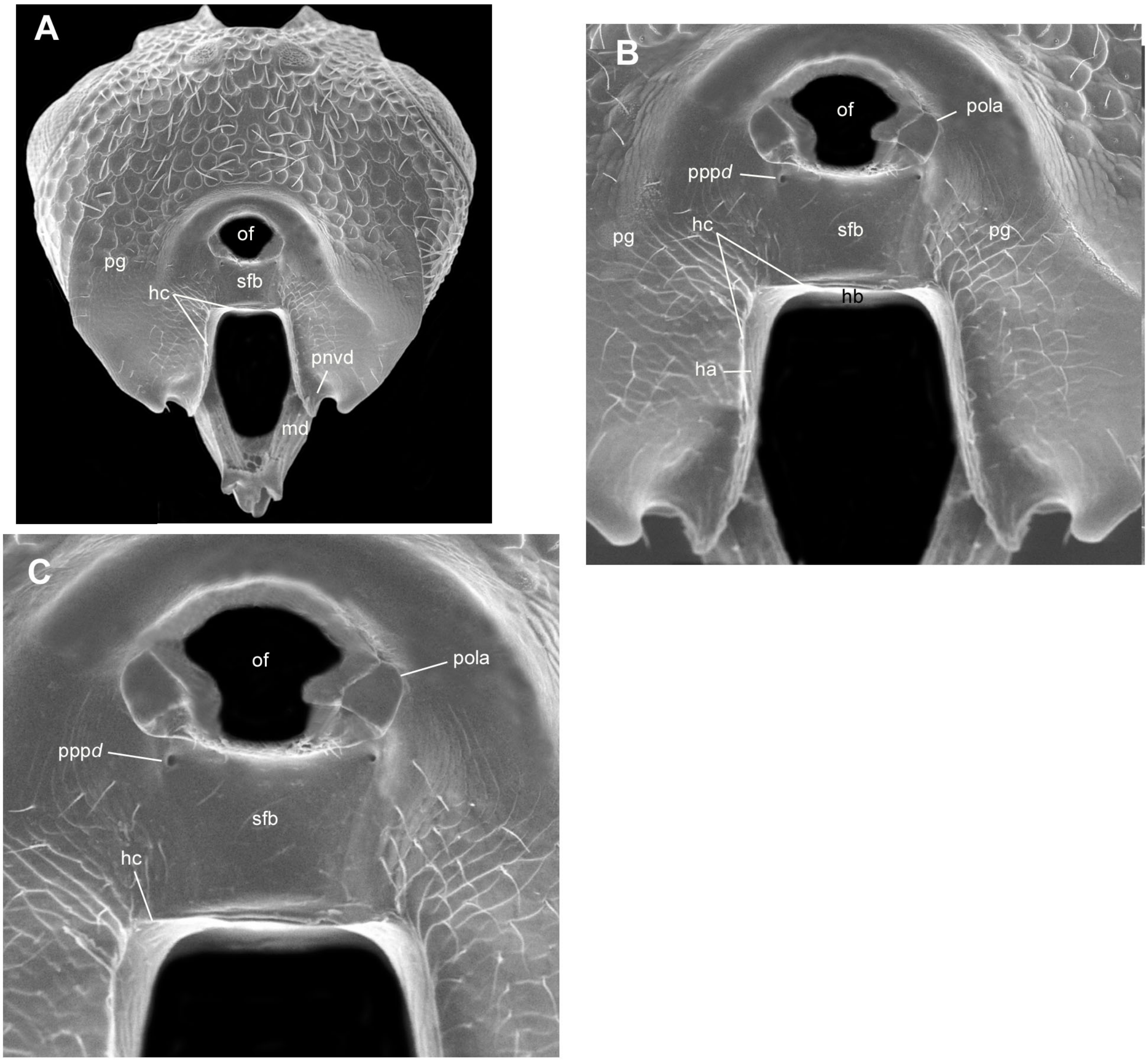


**Figure S4.9**. SEM of head (**A**), ventral half of head (**B**) and subforaminal bridge (**C**) of *Dirhinus* (Chalcididae: Dirhininae); all images in caudal view. **Abbreviations**. hb, hypostomal carina; hc, hypostomal carina; of, occipital foramen; pg, postgena; pgvd, ventral depression of postgena; pola, postocciptal lateral arm; pppd, pit at dorsal end of posterior processs; sfb, subforaminal bridge.

#### S4.5. Conclusion

In conclusion, the possible transformation series that may be hypothesized from the structure of the cephalic capsule of the Chalcididae is presented in Fig. S4.10.

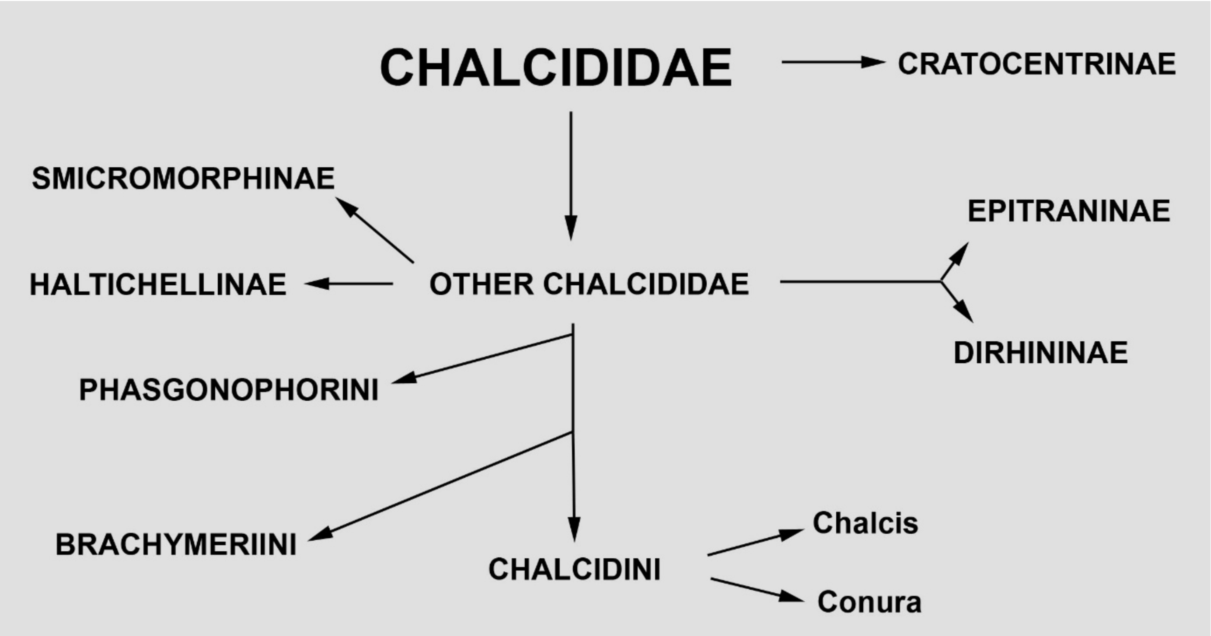

**Figure S4.10**. Hypothesis of transformation series of the structure of the cephalic capsule in Chalcididae
