## AppendixS5 for "Ultra-Conserved Elements and morphology reciprocally illuminate conflicting phylogenetic hypotheses in Chalcididae (Hymenoptera, Chalcidoidea)"

### Appendix S5. Detailed discussion about the morphological phylogeny of the Chalcididae.

This appendix is a more detailed discussion to the results of our morphological analyses. We compare the different topologies and relationships inferred by MP, ML and Bayesian analyses of the morphological characters (Figures 4 and S8) and highlight the morphological character states supporting them. We also list the character states supporting the main clades observed in our inferences. We finally list character states supporting each of the conflicting topologies obtained with our UCE dataset (Topologies A, B, C). Character numbers and states are given between parenthesis. For example, (3,1) is state 1 of character 3. The full list of characters and states can be found in Appendix S1. Stars (*) indicate synapomorphies.

#### S5.1. Monophyly of the Chalcididae

In all analyses, Chalcididae were recovered monophyletic with stronly support (BP MP 100, BP ML 100, PP 1.00). Chalcididae can be defined unequivocally based on 15 synapomorphies [here after indicated with a star *] and 3 homoplastic derived states: labrum exposed and abutting anterior to clypeal margin (3,1)*, labrum plate-like (4,1)*, mandibular base exposed, condyles elongate and visible externally (6,1)*, upper margin of clypeus step-like (14,2)*, cardo stick-like (38,2)*, lateral lamella on anterior tentorial arm moderately to very broad (41,1)*, posterior tentorial arm as a septa re-enforced with 2 sclerotized processes (42,2)*, insertion of dorsal tentorial arm about at level with ventral eye margin (43,1)*, tentorial bridge thin and forming a Y-like structure with the anterior process (52,1)*, multiporous plate sensilla (mps) sunken (55,1), parascutal and axillar carinae with U-shaped connection over tegula at transscutal articulation (62,1)*, pronotum with posteroventral extension that articulates or crosses the prepectus (72,2)*, metepimeron broadly rectangular or square (91,1)*, inner lamella of metadiscrimen strongly raised (93,1)*, metacoxa enlarged and/or elongate (112,1), metafemur enlarged (113,1), metafemur with lobe like teeth (114,2), metatibia with two ventral carinae (120,1)*, petiole entirely sclerotized but not fused with first gastral sternite (127,0)*.

#### S5.2. Monophyly of the currently recognized subfamilies and tribes

In all analyses, Cratocentrinae (BP MP 100, BP ML 100, PP 1), Haltichellinae (BP MP 98, BP ML 98, PP 1), Smicromorphinae (BP MP 100, BP ML 100, PP 1), Epitraninae (BP MP 99, BP ML 92, PP 1) and Dirhininae (BP MP 100, BP ML 98, PP 1) were all retrieved monophyletic with strong support. Chalcidinae as defined before our study were never recovered monophyletic, Phasgonophorini were consistently placed sister to Brachymeriini but did not cluster with Chalcidini. New subfamilies defined in this study were either strongly supported: Chalcidinae (BP MP 100, BP ML 99, PP 1), Brachymeriinae (BP MP 100, BP ML 100, PP 1) or moderately supported: Phasgonophorinae (BP MP 84, BP ML 56, PP 0.63).

##### *S5.2.1. Subfamily Cratocentrinae*

The subfamily is suported by 11 synapomorphies and 3 apomorphic but homoplastic character states: exposed muscle of mandible on the same plane with it and extending into incision on outer surface of mandible (8,1)*, mandible 2-toothed (10,1), postgena distinctly depressed above oral fossa, genal carina absent just above mouth corner, forming a tooth at some distance from it (21,1)*, subforaminal bridge in front of postgenal bridge, not visible from outside (32,1)*, lateral lamella on anterior tentorial arm narrow but with broad apical lobe (41,1)*, two separate claval segments in female (54,1), pronotum with emargination around mesothoracic spiracle (73,1), mesothoracic spiracle visible externally (82,1), mesothoracic discrimen as anchor-like ornamentation with median carina (89,1)*, only a single, median metafurcal pit (92,1), metepisternum with two submedian carinae, parallel and short between metacoxae (96,3)*, fore wing with posterobasal lobe (99,1)*, costal cell with apicoventral tuft of setae (100,1)*, protibia with horizontally directed socketed spur (110,1)*, mesotibia with pegs at apex (111,1)*, transverse carina in front of cercal plates present, cerci situated posteriorly to carina (128,1)*.

We could add to the list the peculiar structure of the gaster in which the gastral tergites 2-4 are weakly sclerotized and only partially visible, hidden by the first tergite.
The states (73,1) and (82,1), relative to the exposition of the mesothoracic spiracle, are ambiguous because the groundplan of the Chalcididae for this state is not known. The mesothoracic spiracle is also hidden in Eurytomidae, the putative sister group of the Chalcididae, possibly figuring a synapomorphy for the two families. If so, the exposed spiracle of the Cratocentrinae is a reversal towards the general condition retrieved in Chalcidoidea. Conversely, a hidden mesothoracic spiracle would be be a homoplastic state if it independently evolved in Eurytomidae and in the clade [Chalcididae minus Cratocentrinae].

##### *S5.2.2. Subfamily Haltichellinae with consideration on tribal relationships*

The subfamily Haltichellinae is strongly supported as monophyletic (BP MP 98, BP ML 98, PP 1) and defined by 9 synapomorphies and 1 homoplastic character state: posterior tentorial sulci short, only linking the posterior process pit to the tentorial, bridge pit (30,3)*, hypostomal carina extended above and joining the postoccipital lateral arm (36,2)*, lateral lamella on anterior tentorial arm very broad and continuing on posterior tentorial arm (41,1)*, postoccipital lateral arm joining ventrally the hypostomal carina (53,1)*, axilla with projecting tooth facing the raised base of axillula (63,1)*, axillula present and completely delimited (64,1)*, inner margin of axillula as raised carina (65,1)*, inner side of metatibia with carina (121,1)*, petiole with complete lamina surrounding ventrally the petiolar foramen of the propodeum (126,1)*, mesofurcal pit on mesotrochantinal plate (90,1). This last state is shared with Smicromorphinae, suggesting a possible sister group relationship with Haltichellinae that was however never retrieved.

*Belaspidia* and *Tropimeris* form a basal grade sister to all other Haltichellinae. In these genera, the clypeus lacks a clear dorsal margin and the malar sulcus is absent. Furthermore, in *Belaspidia,* the mesepisternum lacks the epicnemial carina, whereas epicnemium and ventral shelf are present in all other haltichelline genera. The single autapomorphy for *Belaspidia* is the presence of a posteromedian projection on the mesoscutellum. Conversely, *Tropimeris* exhibits a number of synapomorphies and derived states namely the propodeum with a circular spiracle, its rim being partly hidden by a lobe formed by the anterior end of the sublateral propodeal carina and strongly differ from *Belaspidia*. Therefore, *Belaspidia* and *Tropimeris* must be classified into their own tribes, Belaspidiini **trib. n.** and Tropimeridini Bouček.

The remaining Haltichellinae (*minus Belaspidia* and *Tropimeris*) is supported by the following character states: antennal toruli adjacent to clypeus (17,2)*, lateral and ventral margins of toruli raised (23,1)*, frenum reflexed (67,1), mesothoracic discrimen as raised carina or bump anteriorly, foveate groove posteriorly (89,2). The last two states are also recovered in other subfamilies but are locally apomorphic and unambiguous.

Haltichellini is weakly supported and only two character states were found to support the tribe: 1) the strongly prominent, mostly not sulcate, interantennal projection (25,1)* and 2) the ornamentation of the ventral belt of the prepectus, bearing a sharp medioventral tooth (81,1)*. They are thus mostly defined negatively relatively to their sister group, namely the clade [Notaspidiini + Zavoyini + Hybothoracini]. This heterogeneous clade includes taxa sharing a similar venation of the fore wing, with a short stigmal without uncus (105,1) and missing the postmarginal vein (103,1).

Zavoyini, which includes the single genus *Zavoya* Bouček, is retrieved sister to *Notaspidium* with high support (BP MP 76, BP ML 88, PP 0.99). This relationship is supported by several character states: mandibles bearing two small teeth (10,1), postgena distinctly depressed above oral fossa, the absence of genal carina just above mouth corner, but forming a tooth at some distance from it (21,2), median strip of ornamentation absent on the subforaminal and hypostomal bridges (34,1), hypostoma and hypostomal bridge forming a right angle with subforaminal bridge (39,2), propodeal surface flat and in the same plane as mesonotum (68,1), ventral carinae of metatibia absent (120,0)*. To the exception of the last state, all other are homoplastic, being retrieved in other subfamilies. However, they are unambiguous within Haltichellinae and certainly figure local synapomorphies. We could add the morphology of the gaster. In the two groups, the large first gastral tergite has a broad truncate and carinate base followed by longitudinal carinulae; in addition, the first sternite bears a sharp tooth. In all topologies inferred from UCEs, *Zavoya* forms a grade with *Belaspidia* and *Tropimeris* that is basal to Haltichellinae, which contradicts the character states supporting the sister group relationship between *Zavoya* and *Notaspidium.*

Furthermore, *Notaspidium* and the other Hybothoracini do not cluster together though they share a similar venation with the marginal vein somewhat removed from the front margin of the wing (102,1), a short stigmal vein lacking uncus (105, 1) and an absence of the postmarginal vein (103,1). According to our results (UCEs and morphology), these character states are homoplastic. *Notaspidium* deserves a new tribe, Notaspidiini **trib. n** as it is well defined morphologically by: ventral belt of prepectus with acute median tooth (81,2), lateral panel of prepectus longer than tall (84,1). Besides, the first gastral tergite is large, broadly truncate anteriorly, and with transverse carina followed by longitudinal carinae. It is noteworthy that Notaspidiini should also contain the genus *Steinvreia* Bouček. This last genus was excluded from our analysis because the very few specimens available did not enable us to encode characters of the head, and the amount of missing data hampered phylogenetic inference. However, *Steinvreia* exhibits the apomorphies of the tribe.

Hybothoracini *s.s.* is sustained by the following states: ventral tooth of mandibles longer than the dorsal one (11,1), prosternum with vertical lamina at limit between the ventral and the posterior surfaces (76,3)*, basal tooth of metafemur near base of femur (115,0)*. The first state is homoplastic but locally unambiguous. The last state, figuring a symplesiomorphy for the Chalcididae, is associated with the position of the metatibia at rest, overlapping the base of the femur and hiding it and is therefore apomorphic.

##### *S5.2.3. Subfamily Smicromorphinae*

The subfamily only includes the genus *Smicromorpha* Girault which exhibits highly modified morphology and behavior. Many of these characters are cases of character reversals [here scored #]: gaster weakly sclerotized, collapsing when dried (2,3)#, mandibular base dorsally concealed by genal margin (6,0)#, mouth margin above mandible not incised for reception of mandible (7,0)#, clypeus without visible dorsal margin (14,3), subforaminal bridge at same level with postgena (32,0)#, hypostomal bridge narrower than occipital foramen (33,0)#, hypostomal bridge short or vestigial, much shorter than subforaminal bridge (40,0)#, mps raised above surface of flagellum (55,0)#, pronotum without emargination around mesothoracic spiracle (73,1)#, lateral panel of prepectus not apparent (84,3)*, mesofurcal pit on mesotrochantinal plate (90,1), metepisternal ventral shelf absent, (94,0)#, uncal sensilla grouped in a single cluster (106,0), petiole with basal lamina (124,1) and inserted at base of propodeum (125,1)*. The states (2,3), (55,0) (84,3) and (125,1) are uniquely derived within the family.

##### *S5.2.4. Subfamily Epitraninae*

The subfamily comprises only the genus *Epitranus* Walker, and its monophyly has never been questioned. The following character states support the monophyly of Epitraninae: head with frontal lobe below antennal toruli (22,1)*, interantennal projection prominent and discoid (25,2), postgenal groove present (29,1), cardo triangular (38,0)*, mesothoracic discrimen as raised carina overall (89,2), metepisternal ventral shelf quite long (94,2)*, metatibia with long tarsal scrobe on apicodorsal surface, with a tooth or protrusion above (117,2)*. Additionally, the gaster is strongly bulging ventrally and the antennal scrobes are shallow and often delimited laterally by faint carinae.

##### S5.2.5. *Subfamily Dirhininae*

This is another highly specialized subfamily. Its monophyly has never been contested and is supported by the following derived states: ventral (= inner) tooth of mandibles much shorter than dorsal one (11,2)*, postgena distinctly depressed above oral fossa, genal carina absent just above mouth corner, forming a tooth at some distance from it (21,2), antennal scrobes quite deep and carinately margined laterally (26,2), head with frontal horns (27,1), ventral belt of prepectus with large medioventral tooth (81,2)*, lateral panel of prepectus large, medially foveate and with small anterodorsal projection (83,1)* and (84(1)*, parastigma with hyaline break (101,0), petiole with ventral lamina abutting against petiolar foramen (126,2), petiole fused with first gastral sternite in female (127,1)*. In addition, the propodeum has a peculiar ornamentation including an anteromedian areola and the spiracle is placed on the bottom of a setose depression.

##### S5.2.6. *Subfamily Chalcidinae, with consideration on tribal relationships*

Phasgonophorini (BP MP 84, BP ML 64, PP 0.76) are well defined by 3 synapomorphies and 7 homoplastic states: anterior tentorial pits not visible externally (18,1), malar sulcus absent (19,2), posterior tentorial sulci present (30,2), hypostomal carina forming a complete arch above the hypostoma and hypostomal bridge (36,0), maxillary condyles somewhat distant from each other (37,1)*, hypostomal bridge forming a right to acute angle with subforaminal bridge (39,2), posterior margin of pronotum strongly concave (58,1)*, posteroventral margin of prepectus completely separated from mesepisternum (87,1), protibia without socketed spur but distinctly expanded giving the appearance of a spur (110,2)*, metatibia with long tarsal scrobe (117,1).

We argue that Brachymeriini includes only the genus *Brachymeria*. Indeed, the examination of the type species of all other genera of Brachymeriini showed that only *Brachymeria* can be retained as valid. The Brachymeriini is sustained by 5 synapomorphies and 5 homoplastic derived states: postgena distinctly depressed above oral fossa, the genal carina forming a protrusion at lateral corner of mouth (21,1), posterior tentorial arm a simple, thick and strongly sclerotized process (42,1)*, ventral end of posterior process not joining posterior tentorial arm (50,1), male flagellomeres with modified hairs on the underside (57,1)*, tegula covering humeral plate (59,1), frenum reflexed (67,1), propodeum with setose anterolateral areola (71,1), metepisternum with two submedian carinae, converging posteriorly between metacoxa (96,2)*, first hamulus distant from the others (107,1)*, hind tarsal claw with special spatulate seta (123,1)*.

A sister group relationship between Brachymeriini and Phasgonophorini is sustained by the following homoplastic states: mesothoracic discrimen as raised carina or bump anteriorly, foveate groove posteriorly (89,2), postmarginal vein short, only 1-2 times as long as the stigmal vein and evidently shorter than the marginal vein 104(1), tip of hypopygium near tip of gaster 129(1).

The monophyly of Chalcidini was always recovered and strongly supported (BP MP 100, BP ML 98, PP 1). The subfamily is sustained by: cardo fusiform (38,3)*, ventral end of posterior process distant from posterior tentorial arm (50,1), frenum reflexed (67,1), propodeum with spiracle in vertical orientation (70,1)*, propodeum with setose anterolateral areola (71,1), mesothoracic spiracle partly and hardly visible as hidden by a patch of hairs on posterior margin of pronotum (82,2)*, procoxa with line of setation on posterior surface (109,1)*, metafemur with line of stout bristles on inner surface (116,1)*, petiole with basal lamina (124,1).
*Hovachalcis* Steffan a key Chalcidini taxon could not be included in our analysis as only one specimen is known (the holotype of the type species) that could not be dissected. Consequently, numerous characters could not be coded and these missing data hampered phylogenetic inference. However, *Hovachalcis* shares a number of derived states with the Chalcidini [e.g. 70(1) and 124(1)].

#### S5.3. Relationships between the subfamilies

##### S5.3.1. Position of Cratocentrinae

Two conflicting positions of the Cratocentrinae are recovered in the UCE topologies.

***1) Cratocentrinae sister to all other Chalcididae*** **[UCE topologies A & C]**. This is supported by 5 apomorphies: mandibular base exposed, condyles elongate and visible externally, mouth margin not incised for reception of mandible (lateral to clypeus) (6,1)*, subforaminal bridge sunk down compared to postgena (31,1)*, lateral lamella on anterior tentorial arm narrow but with broad apical lobe (41,2)*, single metafurcal pit medially (92,1)*, two parallel and short submedian carinae present between metacoxae, (96,3)*.

***2) Cratocentrinae included within the clade (Chalcidinae + Smicromorphinae + Dirihininae + Epitraninae + Brachymeriinae)*** **[UCE topologies B]**. This position is only supported by 2 apomorphies at 3 homoplastic derived characters: ventral margin of torulus in lower third of face, not adjacent to clypeus (17,0) (this character state also occurs in *Brachymeria* and is homoplastic in Chalcidoidea), no raised carina on the inner margin of axillula 65,0)*, ventral ornamentation of metafemur with large, regular, lobe like teeth (114,2)(however Dirhininae have small teeth alike Haltichellinae), basal tooth of metafemur near base of femur (115,0)(character state also present in *Notaspidium*), apex of metatibia diagonally truncate (118,1)*.

##### S5.3.2. Clade [Chalcididae minus Cratocentrinae]

In our analyses, this clade was never supported but is nevertheless sustained by 4 synapomorphies and 7 derived states: mouth margin thickened and incised for reception of dorsal corner of mandible (7,1), hypostomal bridge present and distinct from subforaminal bridge (31,1), hypostomal bridge at least as broad as occipital foramen (33,1), width of median strip of ornamentation at least one-third width of hypostomal bridge (35,1), maxillary condyles far from each other (37,2)*, hypostomal bridge at least as long as subforaminal bridge (40,1), apical part of anterior tentorial arm forming process along lateral edge of hypostomal bridge (46,1)*, pit at dorsal end of hypostomal process present (47,1), propodeal spiracle slit-like (69,1), emargination of pronotum around mesothoracic spiracle inconspicuous (73,2), mesothoracic spiracle hidden externally (82,2)*; metafurcal pits absent (92,2)*. The projection of the mesothoracic spiracles is ambiguous because they are also hidden in the Eurytomidae. It might therefore represent a synapomorphy for this most inclusive clade [Eurytomidae + Chalcididae] or may have evolved independently in Eurytomidae and in the clade [Chalcididae *minus* Cratocentrinae].

##### S5.3.3. Relationships within Chalcidinae

The relationships between the Chalcidini and other tribes and subfamilies were always unresolved (Wijesekara 1997a and 1997b; Heraty et al. 2013). In the MP analysis, the tribe is recovered sister to the clade [(Smicromorphinae (Dirhininae, Epitraninae) Haltichellinae)] with weak support (BP<50). This relationship is mostly based on reversals (absence of the dorsolateral postoccipital (28,0) and postgenal groove (29,0), mesepisternum without epicnemium (88,0)) which would also require secondary reversals in some of these subfamilies. Two other character states that support this relationship are obviously homoplastic (antennal scrobes not laterally carinate (26,1) and petiole with basal lamina (126,3)). No morphological characters support a close relationship between Chalcidinae and Haltichellinae (as observed in the UCE topology C).

##### S5.3.1. Clade [Cratocentrinae – Phasgonophorinae]

ML and Bayesian analyses recovered this clade with weak support (BP ML 70, PP 0.84). This relationship is only supported by a single apomorphy and 5 homoplastic derived states: Anterior tentorial pits not visible externally (18,1), width of median strip of ornamentation (mso) narrow, less than one quarter width of hypostomal bridge (35,0), ata-pta intersection far from base of maxillary condyles (mc) (44,2), tarsal scrobe on apicodorsal surface of metatibia present, long, without tooth or protrusion above (117,1), no spurs on metatibia (*Stypiura* bears one spur), tarsal claws with basal tooth (122,2)*.

##### S5.3.2. *Clade [Smicromorphinae + Dirhininae + Epitraninae + Haltichellinae]*

This clade is poorly supported in our parsimony analysis (BP MP 59) but more strongly supported in our ML (BP 81) and Bayesian analyses (PP 0.83). The clade is supported by the following character states: mandible with channel on posterior surface (13,1), clava one-segmented in both sexes (54,2) (56,2)*, tegula covering humeral plate (59,1), posteroventral margin of prepectus completely separated from mesepisternum (87,1), metepisternum with median groove between metacoxae (96,0), humeral plate with reduced number of setae (98,1)*, postmarginal vein of fore wing absent (104,2), metafemur serrulate on ventral margin (114,1). The first character could not be checked in several haltichelline species, the others are either homoplastic [(59,1) (87,1) (114,1)), or figure a symplesiomorphy for the Chalcididae (96,0); at last the state (104,2) needs an unrealistic reversal for *Belaspidia* and the Haltichellini (postmarginal vein reappearing after having been lost). In addition, the cephalic capsule of the Haltichellinae exhibits original characters that contradict a close relationship with the other subfamilies included in this clade.

##### S5.3.3. *Clade [Smicromorphinae + Dirhininae + Epitraninae]*

In all analyses the Smicromorphinae was recovered sister to Dirhininae + Epitraninae always with low support. This relationship is supported by the following character states: median strip of ornamentation on hypostomal bridge absent or virtually so (34,1)*, axillula absent or not differentiate (64,0)*, mesoscutellum without differentiate frenum (66,0)*, postmarginal vein absent (104,2), uncus of stigmal vein absent (105,1). All these states are ambiguous and homoplastic. A sister group relationship between Smicromorphinae and Epitraninae was never retrieved in our analyses. However, several characters may support a close relationship between *Smicromorpha* and *Epitranus*. Both genera share a long petiole and symmetric mandibles in which the ventral tooth is longer than the other teeth. Furthermore, *Epitranus* and *Smicromorpha* species exhibit pale coloration of the body, a feature recurrently associated with nocturnal activity in other Chalcid wasps (i.e. some Agaonidae, Epichrysomallinae etc). Some *Epitranus*, such as *E. hamoni* (Risbec) are only collected with light trap and may have a nocturnal activity, others frequent caves where they search for Tineidae larvae developing on bat dung (Bouček 1982). In the same manner, *Smicromorpha*, (e.g. *S. doddi* Girault) are frequently collected at light.

##### S5.3.4. *Clade [Dirhininae – Epitraninae]*

In all analyses, Dirhininae was recovered sister to Epitraninae with relatively strong support (BP MP 88, BP ML 75, PP 0.96). This relationship is supported by 3 synapomorphies and 13 homoplastic derived states: both mandibles 3-toothed (9,0), mandibular teeth exodont (12,1)*, median strip of ornamentation on hypostomal bridge absent or virtually so (34,1), hypostomal carina a complete arch above the hypostomal bridge (36,0), hypostoma and hypostomal bridge forming a right to acute angle relative to the subforaminal bridge (39,2), dorsal end of posterior process not joining ventral margin of occipital foramen (48,1)*, pits at ventral end of posterior process absent (51,0), axillula absent or not differentiate (64,0), mesoscutellum without differentiate frenum (66,0), propodeum flat and in the same plane as mesonotum (68,1), metepisternal shelf with submedian teeth at posteroventral edge (95,1)*, postmarginal vein absent (104,2), single median carina present between the metacoxae (96,1), marginal vein of fore wing quite long (103,3), uncus of stigmal vein absent (105,1), postmarginal vein absent (104,2); tarsal groove present on metatibia (117,1). Our list also includes apomorphies attributed to the clade [Smicromorphinae (Dirhininae, Epitraninae)], however, the apomorphies mostly result from missing data in *Smicromorpha*. We consider this relationship unrealistic as it also hypothesizes unlikely reversals in either the Dirhininae or the Epitraninae.

#### S5.4. Conclusion

Our morphological study supports the monophyly of Chalcididae with strong support. It also supports the monophyly of most subfamilies (Cratocentrinae, Dirhininae, Epitraninae, Smicromorphinae and Haltichellinae) and of all Chalcidinae tribes. The subfamily Chalcidinae was not recovered monophyletic. Therefore, all Chalcidinae tribes are raised to subfamily rank (Phasgonophorinae stat. rev., Brachymeriinae stat. rev. and Chalcidinae stat. rev). The present subdivision of Haltichellinae into tribes appears inconsistent, indeed Haltichellini and Hybothoracini were recovered polyphyletic. Therefore, two new tribes are proposed: the Belaspidiini for the single genus *Belaspidia* and the Notaspidiini that include the genera *Notaspidium* and *Steninvreia*. Horewever, morphological characters failed at resolving the deepest nodes of the phylogeny. This is certainly due to the presence of highly specialized groups exhibiting a strongly modified body relatively to the groundplan of the family, which complicates the interpretation of homologies, especially for the Smicromorphinae. The amount of convergence also constitutes a background noise that obscure relationships for the deepest nodes.

A carefull analysis of all character states reavealed that between the 3 conflicting topologies inferred from UCEs the one that received the most support from morphological data was topology C. No characters supported Chalcidini sister to Haltichellinae as recovered in topology A. Some morphological support was recovered for topology B, though less than for topology C.
