## AppendixS6 for "Ultra-Conserved Elements and morphology reciprocally illuminate conflicting phylogenetic hypotheses in Chalcididae (Hymenoptera, Chalcidoidea)"

### Appendix S6. Diagnoses of the family, subfamilies and tribes

#### S6.1. Family Chalcididae

Body generally without metallic reflections and strongly sclerotized with coarse sculpture, umbilic punctures on head and dorsum of mesosoma; genal carina generally present; prepectus reduced and hardly visible, and/or mesothoracic spiracle hidden; metacoxa and metafemur enlarged; ventral margin of metafemur either toothed or serrulate; fore wing not folded longitudinally at rest.

#### S6.2. Subfamily Cratocentrinae

Body very strongly sclerotized on various parts, especially on terminal tergites, often with silvery fasciae, formed with dense and adpressed setation; clypeus transverse; gena with an acute tooth above corner of oral fossa in connection with the depressed ventral surface of the postgena; antennal scrobes completely delimited; mesosoma partly cristate dorsally; lateral panel of pronotum and mesopleuron with deep depressions for accommodation of the pro- and mesofemora; propodeum strongly sloping and bearing either long, erect and thin setae overall or adpressed setation on a horizontal spiracular areola; protibia with a socketed apicodorsal spur; mesotibia with pegs on its apicodorsal half; metafemur toothed on ventral margin; metatibia with obliquely truncate projection but without spur; fore wing with a basal lobe on posterior margin and a tuft of setae apically on the underside of costal cell; stigmal and postmarginal veins well expanded, the latter longer than the marginal vein; uncus absent; vestigial veins quite visible as folds; petiole entirely concealed dorsally and anelliform ventrally. Second to fourth tergites weakly sclerotized and mostly covered by first tergite; syntergum often divided by a transverse step or carina and never expanded into a stylus; ovipositor sheaths often very long.

#### S6.3. Subfamily Brachymeriinae status rev.

Body most often black, sometimes with red parts, rarely entirely red, exceptionally yellow or with evident metallic reflections; tegula and legs at least partly bright or pale yellow; species most often with lanceolate silvery setae, sometimes masking the integument, especially on lower face and frons; mandible formula generally 2.3 but occasionally 2.2; clypeus transverse; gena very often with special postorbital carina that diverges from the posterior margin of eye; scrobal depression completely smooth and at least carinately delimited laterally; interantennal space with a laterally compressed projection; frenum often reflexed and always delimited by a raised carina; propodeum completely areolate, generally with a median areola and a delimited anterolateral callus bearing dense setation; epicnemial carina complete, delimiting anteriorly a horizontal ventral shelf; transepimeral groove present; procoxa depressed anteriorly and laterally carinate; metacoxa flattened on outerdorsal surface; metafemur with ventral margin toothed; metatibia obliquely truncate at apex and with a single spur; hind claw with a special, spatulate and falcate seta; fore wing with postmarginal vein only slightly more than twice as long as the stigmal vein; first hamulus distant from the following ones.

#### S6.4. Subfamily Phasgonophorinae status rev.

Body hard, most often with rasp-like sculptures on the mesonotum; no trace of malar sulcus; antennal scrobes quite deep and entirely carinately delimited; frenum delimited anteriorly but sloping, not reflexed; propodeum often strongly sloping; mesopleuron always with a ventral shelf that is often sloping and becomes longer than the epicnemium; procoxa with a deep depression on anterior side, delimited by an oblique carina raised into a flange; postmarginal vein relatively short, only slightly surpassing the stigmal vein in length; petiole very short, almost entirely concealed dorsally and visible as a ring-like sclerite but more visible ventrally; syntergum often elongated into a stylus.

##### *S6.4.1. Tribe Phasgonophorini*

Antennal toruli about at the same level as the lower ocular line; metatibia lacking any spur; ovipositor sheaths straight.

##### *S6.4.2. Tribe Stypiuriini trib. n.*

Lower margin of antennal toruli well above lower ocular line; metatibia with a short, hardly visible apical spur; ovipositor sheaths curved downwards apically.

#### S6.5. Subfamily Chalcidinae

Lower face somewhat receding; clava 3-segmented in both sexes in the Chalcidini [but clava 1-segmented in *Hovachalcis*]; frenum reflexed and always delimited by a raised carina; propodeal spiracle vertical; propodeum with a well delimited anterolateral callus; mesopleuron without epicnemial carina, the ventral carina, visible in many *Conura* being an extension of the ridge bordering anteriorly the femoral depression [but complete epicnemial carina present in *Hovachalcis*]; procoxa with a row of setae on the posterior side; metafemur with a row of large ones on the inner side; metatibia obliquely truncate at apex and bearing a single spur; submarginal vein ending in a discoloured strip and postmarginal vein at least as long as the marginal vein and much longer than the stigmal vein; petiole apparent dorsally, most often obvious and sometimes very long; petiole with a basal lamina, most often ventrally and dorsally as well and inserted at apex of propodeum; all gastral tergites well sclerotized.

#### S6.6. Subfamily Dirhininae

Head high in frontal view; frons with lateral horns and scrobal depression quite deep; mandible elongate and exodont; clypeus about at long as high; anterior tentorial pits visible; malar sulcus completely absent; mesosoma compressed dorsoventrally; frenum and axillulae absent on mesoscutellum; propodeum with a special ornamentation including anteromedian areola and propodeal spiracle on the bottom of a depression; lateral panel of prepectus large, foveate with small anterodorsal projection, ventral belt bearing a large tooth; mesodiscrimen visible as a crenulate groove; procoxa depressed anteriorly and laterally carinate; metafemur serrulate on ventral margin; metatibial apex as a curved spine and without any spur; submarginal vein apex as a discoloured strip; marginal vein quite long compared to the very short stigmal and postmarginal veins; gaster petiolate, most often with 2 pairs of dorsal carinae; first gastral tergite generally with a strigose surface.

#### S6.7. Subfamily Epitraninae

Head most often with frontoclypeal lobe, clypeus reflexed and entirely hidden; mandible moderately long an exodont; antennal scrobes vestigial and weakly delimited laterally; temple with postorbital groove; pronotum and mesonotum with umbilic punctures; axillae and frenum absent on mesoscutellum; propodeum horizontal; mesopleuron with complete epicnemial carina and ventral shelf; metapleuron with long ventral shelf; procoxa depressed anteriorly and laterally carinate; metafemur toothed, less frequently serrulate, on ventral margin; metatibia with a conspicuous tarsal sulcus following a dorsal protrusion and ending as a curved spine; submarginal vein without discoloured strip at apex; marginal vein quite long compared to the very short stigmal and postmarginal veins; petiole long, with kneecap entering propodeal foramen; gaster strongly prominent ventrally.

#### S6.8. Subfamily Smicromorphinae

Body mostly yellowish testaceous, possibly with brown or black spots; mandible with a long, acute and falcate lower tooth; flagellum 5- to 7-segmented with raised multiporous plate sensilla; frenum and axillae absent on mesoscutellum; mesopleuron without epicnemial carina, as the ventral carina, visible in some *Smicromorpha*, is actually an extension of the ridge that borders anteriorly the femoral depression; metafemur with ventral margin serrulate; metatibia obliquely truncate at apex and with a single spur; petiole inserted at the base of propodeum, tail-like and with a basal lamina; gaster hardly sclerotizedn, shriving when dried.

#### S6.9. Subfamily Haltichellinae

Axillula forming a raised plate on the side of mesoscutellum, with its dorsal margin carinately delimited and its anterior apex facing a tooth on the axilla; metatibia truncate at apex and bearing two spurs; petiole with anteroventral margin forming a flange embracing the edge of the propodeal foramen. Sexual dimorphism is most often evident in male antenna: apex of scape sometimes excavated ventrally or/and bearing a ventral projection, first flagellomere strongly transverse; the rest of flagellum thicker and eventually bent.

##### *S6.9.1. Tribe Belaspidiini trib. n.*

Upper margin of the clypeus only discernible by the presence of clearly visible tentorial pits; base of antennal toruli clearly distant from those pits; malar sulcus absent; interantennal space simply bumped; clava 1-segmented in female; mesoscutellum with a median tooth or tubercle at apex; frenum irregularly delimited anteriorly and sloping; mesepisternum lacking epicnemial carina and ventral shelf; procoxa not depressed on anterior side and not carinate; marginal vein at the anterior margin of the fore wing, stigmal and postmarginal vein well expanded; sexual dimorphism slight, with the male flagellomeres (including the first) only moderately shorter than those of the female.

##### *S6.9.2. Tribe Tropimeridini*

Head subtriangular in frontal view; malar space long in contrast with the narrow oral fossa; clypeus with upper margin discernible only by change in sculpture; antennal toruli distant from clypeus and adjacent to each other; interantennal space with small flange; antennal scrobes shallow; mesoscutellum lacking frenum, with posterior margin rounded; propodeal spiracle circular, its rim partly hidden by a lobe formed by the anterior end of the sublateral propodeal carina; mesopleuron with complete epicnemial carina and ventral shelf; procoxa carinate; metafemur quite slender with at least a sharp tooth at mid-length, possibly followed by a subapical one; metatibia with or without additional outer carina; fore wing with short marginal vein removed from margin of wing; stigmal vein very short; postmarginal absent; gaster broadly truncate anteriorly, with transverse carina followed by longitudinal wrinkles; female syntergum with dorsal bump; sexual dimorphism slight but special: male gena very faintly pubescent, mimicking velvet, syntergum without evident dorsal bump.

##### *S6.9.3. Tribe Haltichellini*

Mandibular formula most often 2.3; clypeus frequently receding and only partly visible in frontal view; antennal toruli generally adjacent to upper margin of clypeus; interantennal projection frequently strongly projecting and compressed laterally, with a subcircular edge; clava 1-segmented in female; mesoscutellum nearly always projecting posteriorly, with frenal carina forming submedian or median lobe(s) and frenum reflexed; axillula subtriangular, raised relatively to surface of mesoscutellum ; propodeum with submedian and sublateral carinae converging posteriorly, the latter often forming a spiracular tooth at their anterior end; ventral belt of prepectus with acute median tooth; mesepisternum with mesodiscrimen as a bump of carina dorsally and a crenulate groove of fovea ventrally, epicnemial carina completely delimiting epicnemium and ventral shelf; procoxa depressed anteriorly and laterally carinate; metafemur with ventral margin variously ornamented but basically with a tooth at mid-length followed by a serrulate lobe; metatibia not overlapping base of femur at rest, with or without additional outer carina; fore wing with marginal vein nearly always on anterior margin; stigmal vein variously expanded; postmarginal present although possibly vestigial.

##### *S6.9.4. Tribe Notaspidiini trib. n.*

Body very often with bright metallic reflections (in *Notaspidium*); prosternum lacking flange at junction between ventral and posterior surfaces; ventral belt of prepectus with acute median tooth; prepectus relatively large (quite so in *Steinvreia*); fore wing with marginal vein removed from wing margin; stigmal vein vestigial and postmarginal absent; first gastral tergite large, broadly truncate anteriorly, with transverse carina followed by longitudinal carinae.

##### *S6.9.5. Tribe Zavoyini*

Head with deep scrobal depression delimited by a raised carina; eye with sparse but long setae; mesoscutellum produced into a long horizontal horn; metafemur slender, its ventral margin with a tooth near base followed by serrulation; metatibia stout at apex, with a single apical spur; marginal vein excessively long but not tubular; stigmal vein very short; and postmarginal vein absent; first gastral tergite very large, truncate anteriorly, with transverse carina followed by longitudinal carinae; first sternite with sharp ventral tooth.

##### *S6.9.6. Tribe Hybothoracini*

Clypeus often strongly receding, only partly visible in frontal view; antennal toruli adjacent to upper margin of clypeus; interantennal projection moderately prominent and broadly sulcate, not laterally compressed; frontal depression generally not very deep and without preorbital carina; first flagellomere often elongate in female and discoid in male; pronotal carina only visible dorsally; frenal carina sometimes delimiting submedian lobes; axillula narrow and elongate; propodeum entirely areolate, prosternum with triangular flange at junction between ventral and posterior surfaces; mesopleuron with epicnemial carina mostly evanescent along the large epicnemial areola; metafemur with ventral margin with a tooth near base, sinuate and serrulate apically up to the subapical lobe; metatibia overlapping base of femur and hiding it at rest; metatibia most often without additional outer carina; fore wing with marginal vein removed from anterior margin of wing; stigmal vein short; postmarginal vein absent; petiole barely visible, quite transverse in dorsal view; first gastral tergite variable either rounded at base or more or less broadly truncate.
