## Supplementary material for "Ultra-Conserved Elements and morphology reciprocally illuminate conflicting phylogenetic hypotheses in Chalcididae (Hymenoptera, Chalcidoidea)": FigureS1

**Figure S1A**
RAxML tree, complete UCE data set
no partition
Bootstrap replicates at nodes (100 replicates)
Current classification is used to annotate tree

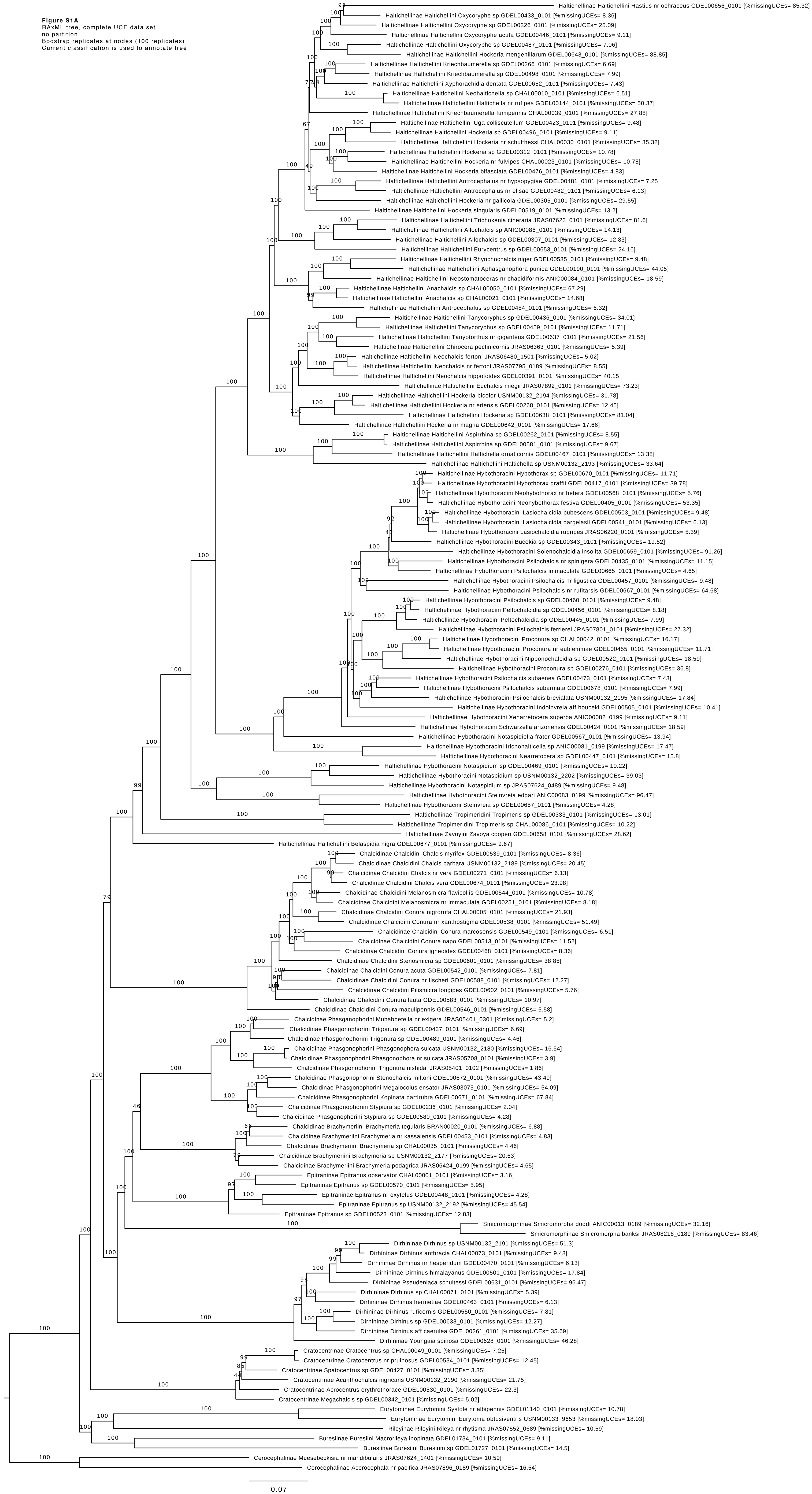

**Figure S1B**
RAxML tree, complete UCE data set with partition
Boostrap replicates at nodes (100 replicates)
Current classification is used to annotate tree

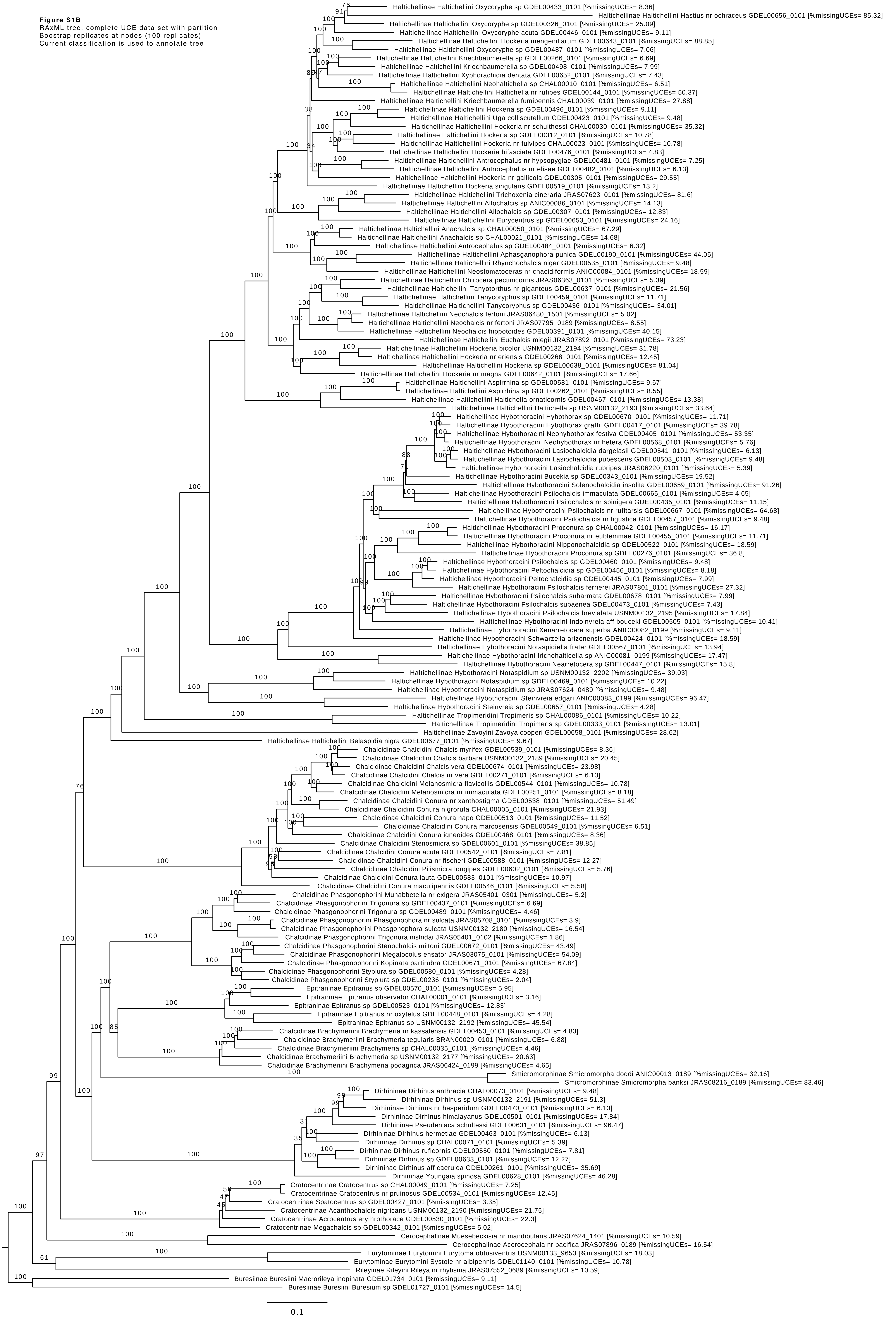

**Figure S1C**
midpoint rooted RAxML tree, complete UCE data set
no outgroup
no partition
Bostrap replicates at nodes (100 replicates)
Current classification is used to annotate tree

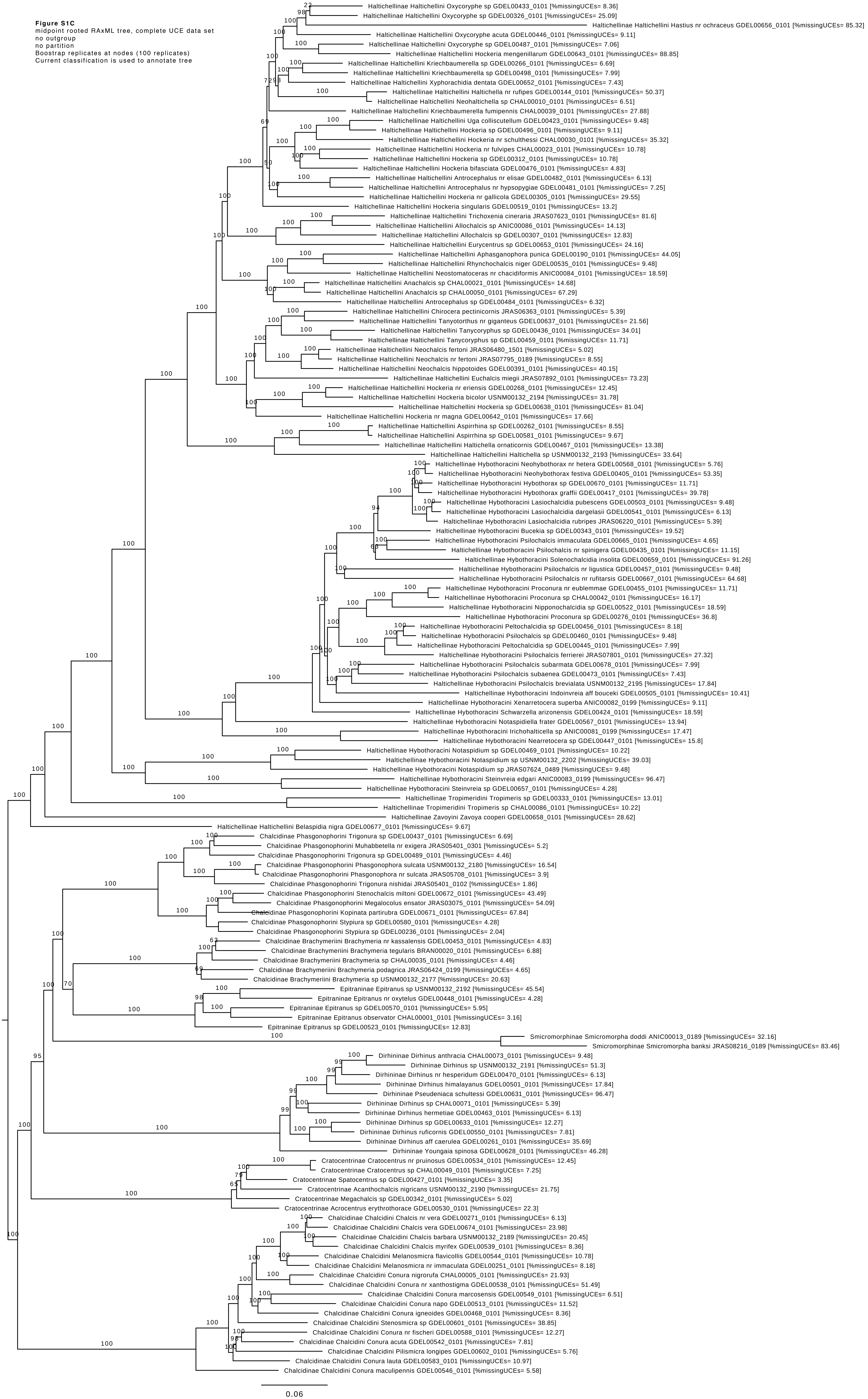
