## Supplementary material for "Ultra-Conserved Elements and morphology reciprocally illuminate conflicting phylogenetic hypotheses in Chalcididae (Hymenoptera, Chalcidoidea)": FigureS2

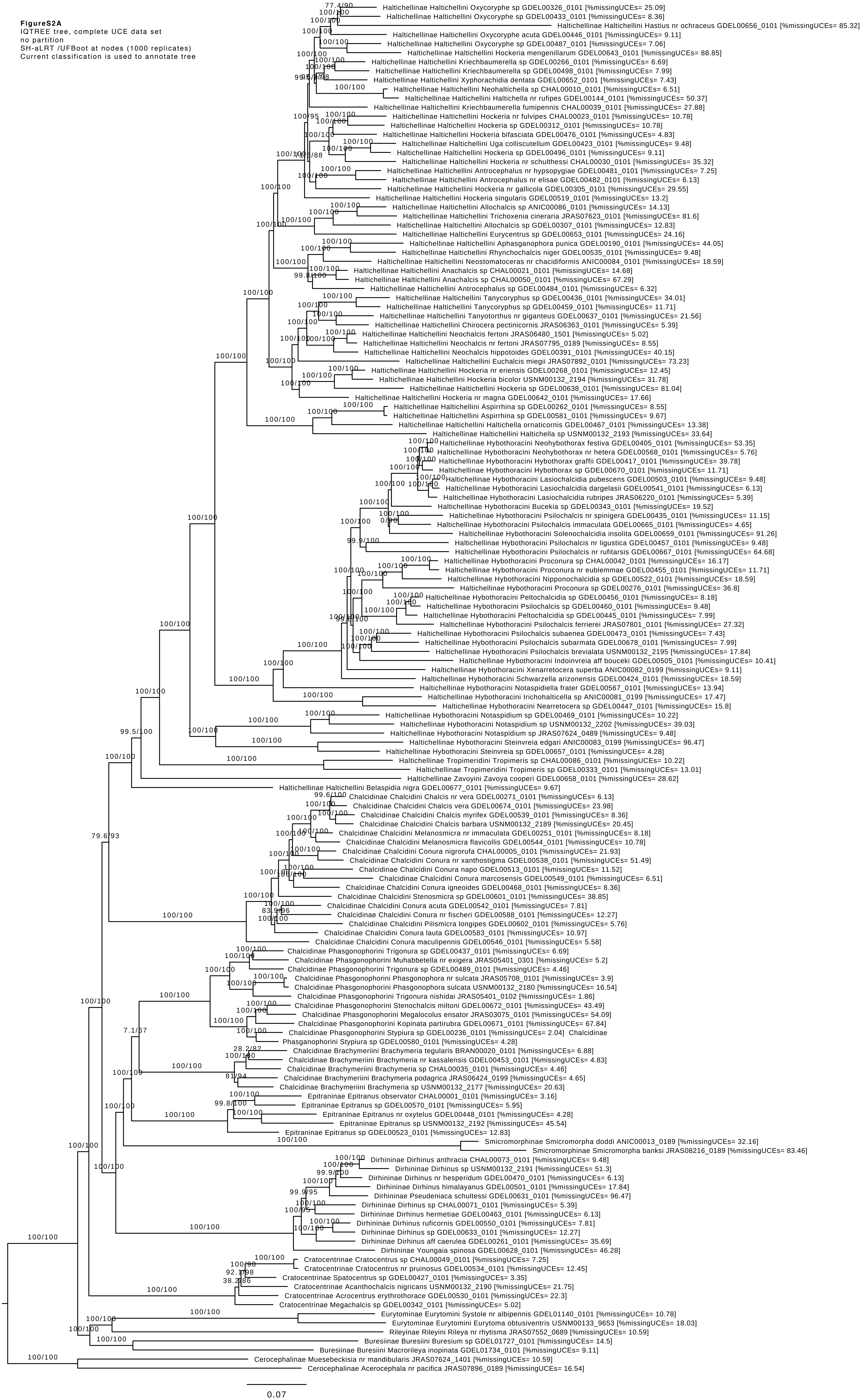


**FigureS2A**
IQTREE tree, complete UCE data set
no partition
SH-aLRT /UFBoot at nodes (1000 replicates)
Current classification is used to annotate tree

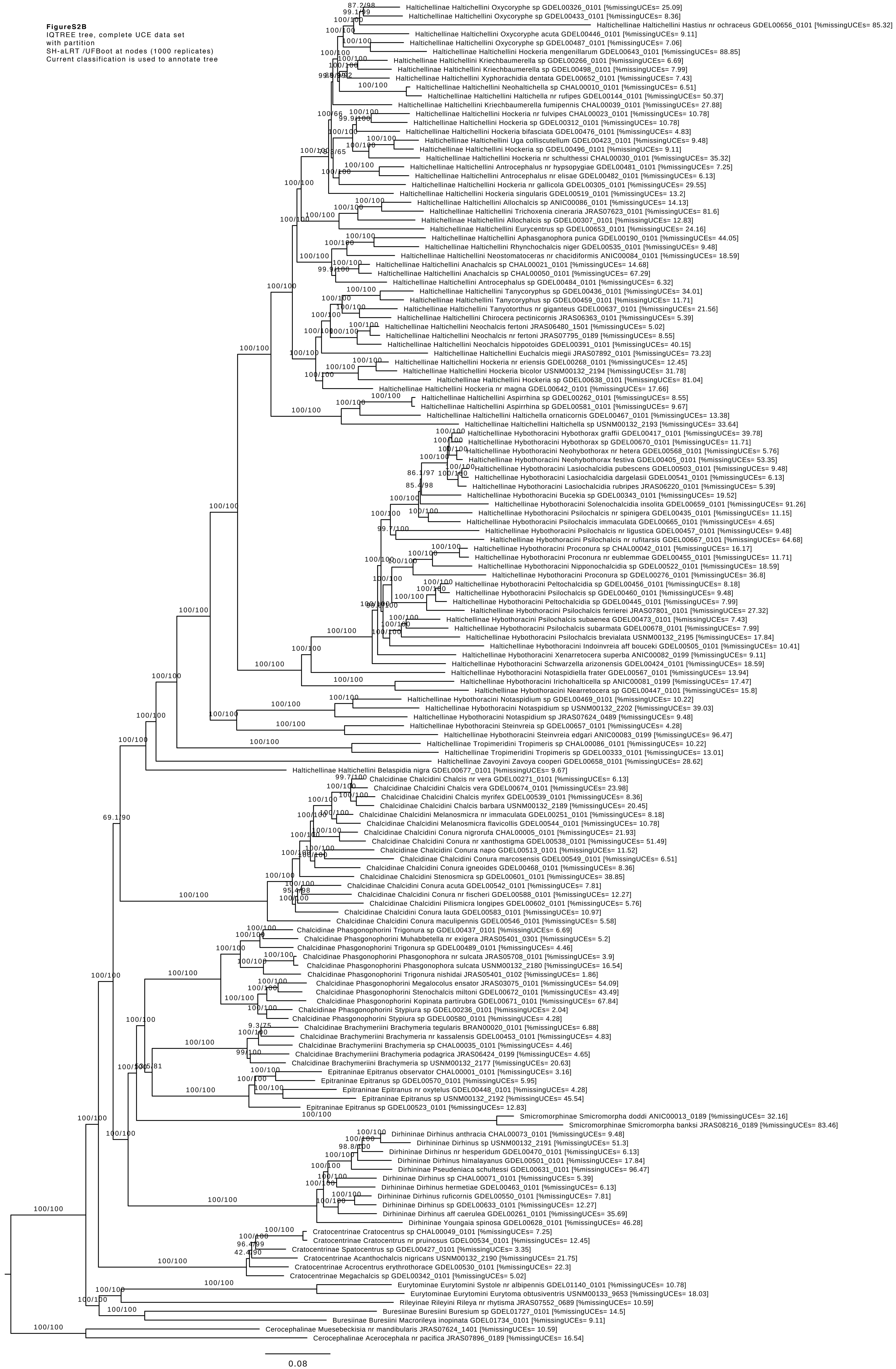
FigureS2B
IQTREE tree, complete UCE data set
with partition
SH-aLRT /UFBoot at nodes (1000 replicates)
Current classification is used to annotate tree
Haltichellinae Haltichellini Oxycoryphe sp GDEL00326_0101 [%missingUCEs= 25.09]
Haltichellinae Haltichellini Oxycoryphe sp GDEL00433_0101 [%missingUCEs= 8.36]
Haltichellinae Haltichellini Hastius nr ochraceus GDEL00656_0101 [%missingUCEs= 85.32]
Haltichellinae Haltichellini Oxycoryphe acuta GDEL00446_0101 [%missingUCEs= 9.11]
Haltichellinae Haltichellini Oxycoryphe sp GDEL00487_0101 [%missingUCEs= 7.06]
Haltichellinae Haltichellini Hockeria mengenillarum GDEL00643_0101 [%missingUCEs= 88.85]
Haltichellinae Haltichellini Kriechbaumerella sp GDEL00266_0101 [%missingUCEs= 6.69]
Haltichellinae Haltichellini Kriechbaumerella sp GDEL00498_0101 [%missingUCEs= 7.99]
Haltichellinae Haltichellini Xyphorachidia dentata GDEL00652_0101 [%missingUCEs= 7.43]
Haltichellinae Haltichellini Neohaltichella sp CHAL00010_0101 [%missingUCEs= 6.51]
Haltichellinae Haltichellini Haltichella nr rufipes GDEL00144_0101 [%missingUCEs= 50.37]
Haltichellinae Haltichellini Kriechbaumerella fumipennis CHAL00039_0101 [%missingUCEs= 27.88]
Haltichellinae Haltichellini Hockeria nr fulvipes CHAL00023_0101 [%missingUCEs= 10.78]
Haltichellinae Haltichellini Hockeria sp GDEL00312_0101 [%missingUCEs= 10.78]
Haltichellinae Haltichellini Hockeria bifasciata GDEL00476_0101 [%missingUCEs= 4.83]
Haltichellinae Haltichellini Uga colliscutellum GDEL00423_0101 [%missingUCEs= 9.48]
Haltichellinae Haltichellini Hockeria sp GDEL00496_0101 [%missingUCEs= 9.11]
Haltichellinae Haltichellini Hockeria nr schulthessi CHAL00030_0101 [%missingUCEs= 35.32]
Haltichellinae Haltichellini Antrocephalus nr hypsopygiae GDEL00481_0101 [%missingUCEs= 7.25]
Haltichellinae Haltichellini Antrocephalus nr elisae GDEL00482_0101 [%missingUCEs= 6.13]
Haltichellinae Haltichellini Hockeria nr gallicola GDEL00305_0101 [%missingUCEs= 29.55]
Haltichellinae Haltichellini Hockeria singularis GDEL00519_0101 [%missingUCEs= 13.2]
Haltichellinae Haltichellini Allochalcis sp ANIC00086_0101 [%missingUCEs= 14.13]
Haltichellinae Haltichellini Trichoxenia cineraria JRAS07623_0101 [%missingUCEs= 81.6]
Haltichellinae Haltichellini Allochalcis sp GDEL00307_0101 [%missingUCEs= 12.83]
Haltichellinae Haltichellini Eurycentrus sp GDEL00653_0101 [%missingUCEs= 24.16]
Haltichellinae Haltichellini Aphasganophora punica GDEL00190_0101 [%missingUCEs= 44.05]
Haltichellinae Haltichellini Rhynchochalcis niger GDEL00535_0101 [%missingUCEs= 9.48]
Haltichellinae Haltichellini Neostomatoceras nr chacidiformis ANIC00084_0101 [%missingUCEs= 18.59]
Haltichellinae Haltichellini Anachalcis sp CHAL00021_0101 [%missingUCEs= 14.68]
Haltichellinae Haltichellini Anachalcis sp CHAL00050_0101 [%missingUCEs= 67.29]
Haltichellinae Haltichellini Antrocephalus sp GDEL00484_0101 [%missingUCEs= 6.32]
Haltichellinae Haltichellini Tanycoryphus sp GDEL00436_0101 [%missingUCEs= 34.01]
Haltichellinae Haltichellini Tanycoryphus sp GDEL00459_0101 [%missingUCEs= 11.71]
Haltichellinae Haltichellini Tanyotorthus nr giganteus GDEL00637_0101 [%missingUCEs= 21.56]
Haltichellinae Haltichellini Chirocera pectinicornis JRAS06363_0101 [%missingUCEs= 5.39]
Haltichellinae Haltichellini Neochalcis fertoni JRAS06480_1501 [%missingUCEs= 5.02]
Haltichellinae Haltichellini Neochalcis nr fertoni JRAS07795_0189 [%missingUCEs= 8.55]
Haltichellinae Haltichellini Neochalcis hippotoides GDEL00391_0101 [%missingUCEs= 40.15]
Haltichellinae Haltichellini Euchalcis miegii JRAS07892_0101 [%missingUCEs= 73.23]
Haltichellinae Haltichellini Hockeria nr eriensis GDEL00268_0101 [%missingUCEs= 12.45]
Haltichellinae Haltichellini Hockeria bicolor USNM00132_2194 [%missingUCEs= 31.78]
Haltichellinae Haltichellini Hockeria sp GDEL00638_0101 [%missingUCEs= 81.04]
Haltichellinae Haltichellini Hockeria nr magna GDEL00642_0101 [%missingUCEs= 17.66]
Haltichellinae Haltichellini Aspirrhina sp GDEL00262_0101 [%missingUCEs= 8.55]
Haltichellinae Haltichellini Aspirrhina sp GDEL00581_0101 [%missingUCEs= 9.67]
Haltichellinae Haltichellini Haltichella ornaticornis GDEL00467_0101 [%missingUCEs= 13.38]
Haltichellinae Haltichellini Haltichella sp USNM00132_2193 [%missingUCEs= 33.64]
Haltichellinae Hybothoracini Hybothorax graffii GDEL00417_0101 [%missingUCEs= 39.78]
Haltichellinae Hybothoracini Hybothorax sp GDEL00670_0101 [%missingUCEs= 11.71]
Haltichellinae Hybothoracini Neohybothorax nr hetera GDEL00568_0101 [%missingUCEs= 5.76]
Haltichellinae Hybothoracini Neohybothorax festiva GDEL00405_0101 [%missingUCEs= 53.35]
Haltichellinae Hybothoracini Lasiochalcidia pubescens GDEL00503_0101 [%missingUCEs= 9.48]
Haltichellinae Hybothoracini Lasiochalcidia dargelasii GDEL00541_0101 [%missingUCEs= 6.13]
Haltichellinae Hybothoracini Lasiochalcidia rubripes JRAS06220_0101 [%missingUCEs= 5.39]
Haltichellinae Hybothoracini Bucekia sp GDEL00343_0101 [%missingUCEs= 19.52]
Haltichellinae Hybothoracini Solenochalcidia insolita GDEL00659_0101 [%missingUCEs= 91.26]
Haltichellinae Hybothoracini Psilochalcis nr spinigera GDEL00435_0101 [%missingUCEs= 11.15]
Haltichellinae Hybothoracini Psilochalcis immaculata GDEL00665_0101 [%missingUCEs= 4.65]
Haltichellinae Hybothoracini Psilochalcis nr ligustica GDEL00457_0101 [%missingUCEs= 9.48]
Haltichellinae Hybothoracini Psilochalcis nr rufitarsis GDEL00667_0101 [%missingUCEs= 64.68]
Haltichellinae Hybothoracini Proconura sp CHAL00042_0101 [%missingUCEs= 16.17]
Haltichellinae Hybothoracini Proconura nr eublemmae GDEL00455_0101 [%missingUCEs= 11.71]
Haltichellinae Hybothoracini Nipponochalcidia sp GDEL00522_0101 [%missingUCEs= 18.59]
Haltichellinae Hybothoracini Proconura sp GDEL00276_0101 [%missingUCEs= 36.8]
Haltichellinae Hybothoracini Peltochalcidia sp GDEL00456_0101 [%missingUCEs= 8.18]
Haltichellinae Hybothoracini Psilochalcis sp GDEL00460_0101 [%missingUCEs= 9.48]
Haltichellinae Hybothoracini Peltochalcidia sp GDEL00445_0101 [%missingUCEs= 7.99]
Haltichellinae Hybothoracini Psilochalcis ferrierei JRAS07801_0101 [%missingUCEs= 27.32]
Haltichellinae Hybothoracini Psilochalcis subaenea GDEL00473_0101 [%missingUCEs= 7.43]
Haltichellinae Hybothoracini Psilochalcis subarmata GDEL00678_0101 [%missingUCEs= 7.99]
Haltichellinae Hybothoracini Psilochalcis brevialata USNM00132_2195 [%missingUCEs= 17.84]
Haltichellinae Hybothoracini Indoinvreia aff bouceki GDEL00505_0101 [%missingUCEs= 10.41]
Haltichellinae Hybothoracini Xenarretocera superba ANIC00082_0199 [%missingUCEs= 9.11]
Haltichellinae Hybothoracini Schwarzella arizonensis GDEL00424_0101 [%missingUCEs= 18.59]
Haltichellinae Hybothoracini Notaspidiella frater GDEL00567_0101 [%missingUCEs= 13.94]
Haltichellinae Hybothoracini Irichohalticella sp ANIC00081_0199 [%missingUCEs= 17.47]
Haltichellinae Hybothoracini Nearretocera sp GDEL00447_0101 [%missingUCEs= 15.8]
Haltichellinae Hybothoracini Notaspidium sp GDEL00469_0101 [%missingUCEs= 10.22]
Haltichellinae Hybothoracini Notaspidium sp USNM00132_2202 [%missingUCEs= 39.03]
Haltichellinae Hybothoracini Notaspidium sp JRAS07624_0489 [%missingUCEs= 9.48]
Haltichellinae Hybothoracini Steinvreia sp GDEL00657_0101 [%missingUCEs= 4.28]
Haltichellinae Hybothoracini Steinvreia edgari ANIC00083_0199 [%missingUCEs= 96.47]
Haltichellinae Tropimeridini Tropimeris sp CHAL00086_0101 [%missingUCEs= 10.22]
Haltichellinae Tropimeridini Tropimeris sp GDEL00333_0101 [%missingUCEs= 13.01]
Haltichellinae Zavoyini Zavoya cooperi GDEL00658_0101 [%missingUCEs= 28.62]
Haltichellinae Haltichellini Belaspidia nigra GDEL00677_0101 [%missingUCEs= 9.67]
Chalcidinae Chalcidini Chalcis nr vera GDEL00271_0101 [%missingUCEs= 6.13]
Chalcidinae Chalcidini Chalcis vera GDEL00674_0101 [%missingUCEs= 23.98]
Chalcidinae Chalcidini Chalcis myrifex GDEL00539_0101 [%missingUCEs= 8.36]
Chalcidinae Chalcidini Chalcis barbara USNM00132_2189 [%missingUCEs= 20.45]
Chalcidinae Chalcidini Melanosmicra nr immaculata GDEL00251_0101 [%missingUCEs= 8.18]
Chalcidinae Chalcidini Melanosmicra flavicollis GDEL00544_0101 [%missingUCEs= 10.78]
Chalcidinae Chalcidini Conura nigrorufa CHAL00005_0101 [%missingUCEs= 21.93]
Chalcidinae Chalcidini Conura nr xanthostigma GDEL00538_0101 [%missingUCEs= 51.49]
Chalcidinae Chalcidini Conura napo GDEL00513_0101 [%missingUCEs= 11.52]
Chalcidinae Chalcidini Conura marcosensis GDEL00549_0101 [%missingUCEs= 6.51]
Chalcidinae Chalcidini Conura igneoides GDEL00468_0101 [%missingUCEs= 8.36]
Chalcidinae Chalcidini Stenosmicra sp GDEL00601_0101 [%missingUCEs= 38.85]
Chalcidinae Chalcidini Conura acuta GDEL00542_0101 [%missingUCEs= 7.81]
Chalcidinae Chalcidini Conura nr fischeri GDEL00588_0101 [%missingUCEs= 12.27]
Chalcidinae Chalcidini Pilismicra longipes GDEL00602_0101 [%missingUCEs= 5.76]
Chalcidinae Chalcidini Conura lauta GDEL00583_0101 [%missingUCEs= 10.97]
Chalcidinae Chalcidini Conura maculipennis GDEL00546_0101 [%missingUCEs= 5.58]
Chalcidinae Phasgonophorini Trigonura sp GDEL00437_0101 [%missingUCEs= 6.69]
Chalcidinae Phasgonophorini Muhabbetella nr exigera JRAS05401_0301 [%missingUCEs= 5.2]
Chalcidinae Phasgonophorini Trigonura sp GDEL00489_0101 [%missingUCEs= 4.46]
Chalcidinae Phasgonophorini Phasgonophora nr sulcata JRAS05708_0101 [%missingUCEs= 3.9]
Chalcidinae Phasgonophorini Phasgonophora sulcata USNM00132_2180 [%missingUCEs= 16.54]
Chalcidinae Phasgonophorini Trigonura nishidai JRAS05401_0102 [%missingUCEs= 1.86]
Chalcidinae Phasgonophorini Megalocolus ensator JRAS03075_0101 [%missingUCEs= 54.09]
Chalcidinae Phasgonophorini Stenochalcis miltoni GDEL00672_0101 [%missingUCEs= 43.49]
Chalcidinae Phasgonophorini Kopinata partirubra GDEL00671_0101 [%missingUCEs= 67.84]
Chalcidinae Phasgonophorini Stypiura sp GDEL00236_0101 [%missingUCEs= 2.04]
Chalcidinae Phasgonophorini Stypiura sp GDEL00580_0101 [%missingUCEs= 4.28]
Chalcidinae Brachymeriini Brachymeria tegularis BRAN00020_0101 [%missingUCEs= 6.88]
Chalcidinae Brachymeriini Brachymeria nr kassalensis GDEL00453_0101 [%missingUCEs= 4.83]
Chalcidinae Brachymeriini Brachymeria sp CHAL00035_0101 [%missingUCEs= 4.46]
Chalcidinae Brachymeriini Brachymeria podagrica JRAS06424_0199 [%missingUCEs= 4.65]
Chalcidinae Brachymeriini Brachymeria sp USNM00132_2177 [%missingUCEs= 20.63]
Epitraninae Epitranus observator CHAL00001_0101 [%missingUCEs= 3.16]
Epitraninae Epitranus sp GDEL00570_0101 [%missingUCEs= 5.95]
Epitraninae Epitranus nr oxytelus GDEL00448_0101 [%missingUCEs= 4.28]
Epitraninae Epitranus sp USNM00132_2192 [%missingUCEs= 45.54]
Epitraninae Epitranus sp GDEL00523_0101 [%missingUCEs= 12.83]
Smicromorphinae Smicromorpha doddi ANIC00013_0189 [%missingUCEs= 32.16]
Smicromorphinae Smicromorpha banksi JRAS08216_0189 [%missingUCEs= 83.46]
Dirhininae Dirhinus anthracia CHAL00073_0101 [%missingUCEs= 9.48]
Dirhininae Dirhinus sp USNM00132_2191 [%missingUCEs= 51.3]
Dirhininae Dirhinus nr hesperidum GDEL00470_0101 [%missingUCEs= 6.13]
Dirhininae Dirhinus himalayanus GDEL00501_0101 [%missingUCEs= 17.84]
Dirhininae Pseudeniaca schultessi GDEL00631_0101 [%missingUCEs= 96.47]
Dirhininae Dirhinus sp CHAL00071_0101 [%missingUCEs= 5.39]
Dirhininae Dirhinus hermetiae GDEL00463_0101 [%missingUCEs= 6.13]
Dirhininae Dirhinus ruficornis GDEL00550_0101 [%missingUCEs= 7.81]
Dirhininae Dirhinus sp GDEL00633_0101 [%missingUCEs= 12.27]
Dirhininae Dirhinus aff caerulea GDEL00261_0101 [%missingUCEs= 35.69]
Dirhininae Youngaia spinosa GDEL00628_0101 [%missingUCEs= 46.28]
Cratocentrinae Cratocentrus sp CHAL00049_0101 [%missingUCEs= 7.25]
Cratocentrinae Cratocentrus nr pruinosus GDEL00534_0101 [%missingUCEs= 12.45]
Cratocentrinae Spatocentrus sp GDEL00427_0101 [%missingUCEs= 3.35]
Cratocentrinae Acanthochalcis nigricans USNM00132_2190 [%missingUCEs= 21.75]
Cratocentrinae Acrocentrus erythrothorace GDEL00530_0101 [%missingUCEs= 22.3]
Cratocentrinae Megachalcis sp GDEL00342_0101 [%missingUCEs= 5.02]
Eurytominae Eurytomini Systole nr albipennis GDEL01140_0101 [%missingUCEs= 10.78]
Eurytominae Eurytomini Eurytoma obtusiventris USNM00133_9653 [%missingUCEs= 18.03]
Rileyinae Rileyini Rileya nr rhytisma JRAS07552_0689 [%missingUCEs= 10.59]
Buresiinae Buresiini Buresium sp GDEL01727_0101 [%missingUCEs= 14.5]
Buresiinae Buresiini Macrorileya inopinata GDEL01734_0101 [%missingUCEs= 9.11]
Cerocephalinae Muesebeckisia nr mandibularis JRAS07624_1401 [%missingUCEs= 10.59]
Cerocephalinae Acerocephala nr pacifica JRAS07896_0189 [%missingUCEs= 16.54]
0.08
