## Supplementary material for "Ultra-Conserved Elements and morphology reciprocally illuminate conflicting phylogenetic hypotheses in Chalcididae (Hymenoptera, Chalcidoidea)": FigureS3

**FigureS3A**
ASTRAL tree, complete UCE data set
nodes with Bootstrap < 10 were collapsed in gene trees prior to analysis
Local Posterior Probabilities at nodes
Current classification is used to annotate tree

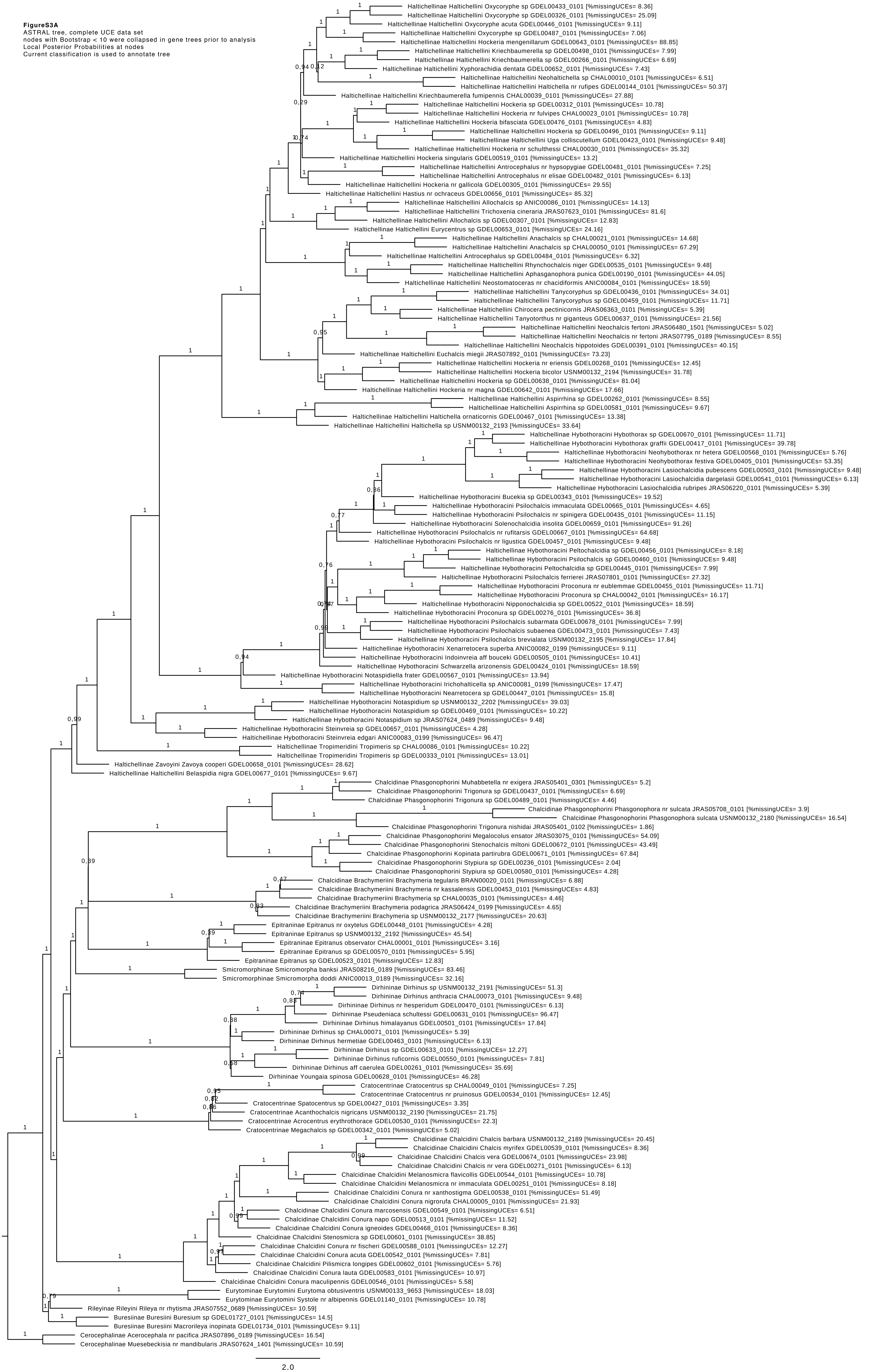

**FigureS3B**
ASTRAL tree, complete UCE data set
nodes with Bootstrap < 50 were collapsed in gene trees prior to analysis
Local Posterior Probabilities at nodes
Current classification is used to annotate tree

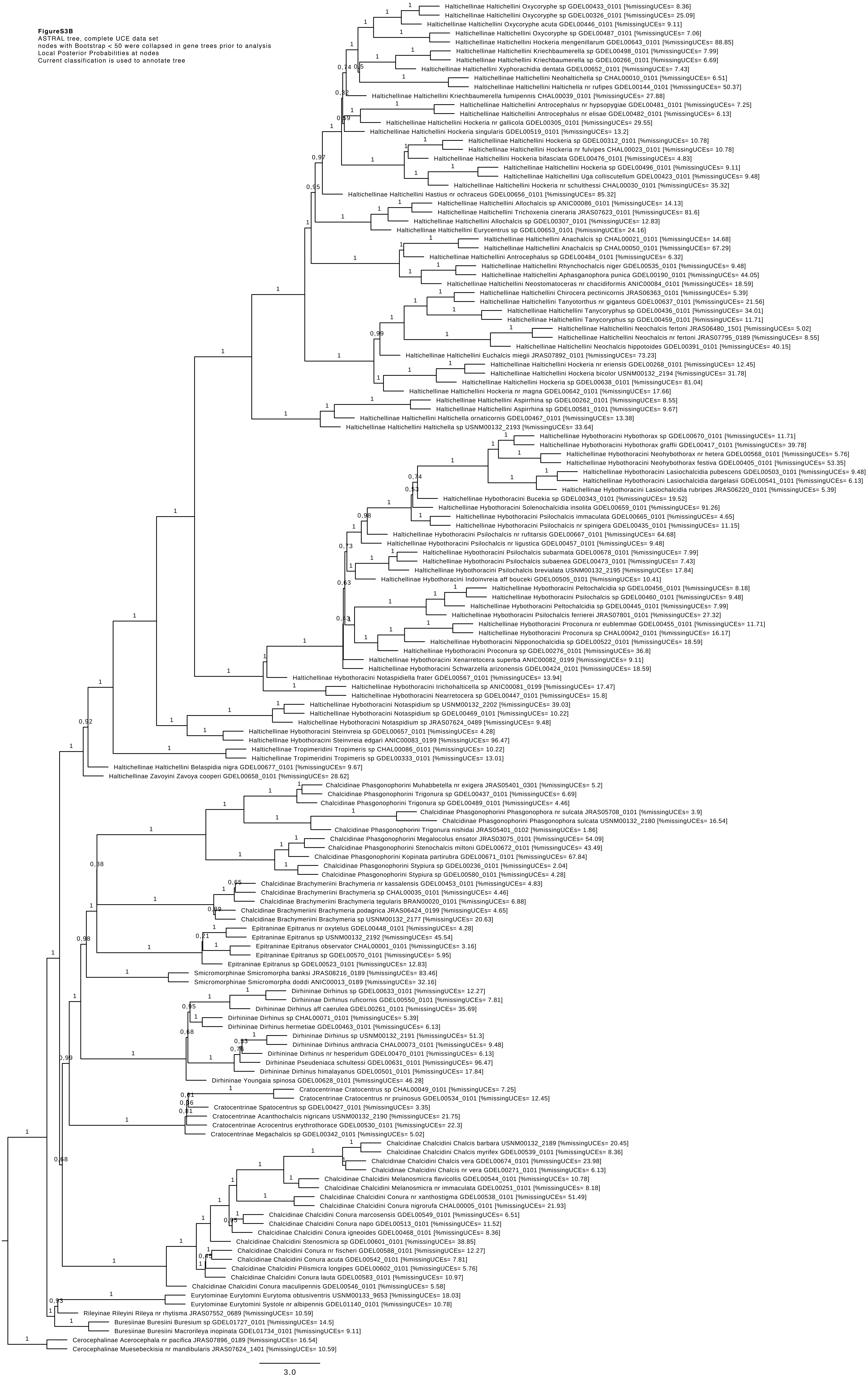

**FigureS3C**
ASTRAL tree, 51 UCEs with monophyletic ingroup
nodes with Bootstrap < 10 were collapsed in gene trees prior to analysis
Local Posterior Probabilities at nodes
Current classification is used to annotate tree

the following UCE trees were used as input :
uce-18, uce-35, uce-58, uce-99, uce-102, uce-103, uce-120, uce-129, uce-134, uce-189, uce-220, uce-265, uce-278, uce-280, uce-327, uce-348, uce-350, uce-374, uce-384, uce-434, uce-442, uce-447, uce-495, uce-583, uce-596, uce-616, uce-635, uce-650, uce-683, uce-694, uce-698, uce-820, uce-855, uce-946, uce-965, uce-966, uce-988, uce-1008, uce-1011, uce-1066, uce-1109, uce-1191, uce-1195, uce-1236, uce-1237, uce-1246, uce-1254, uce-1335, uce-1367, uce-1379, uce-1415

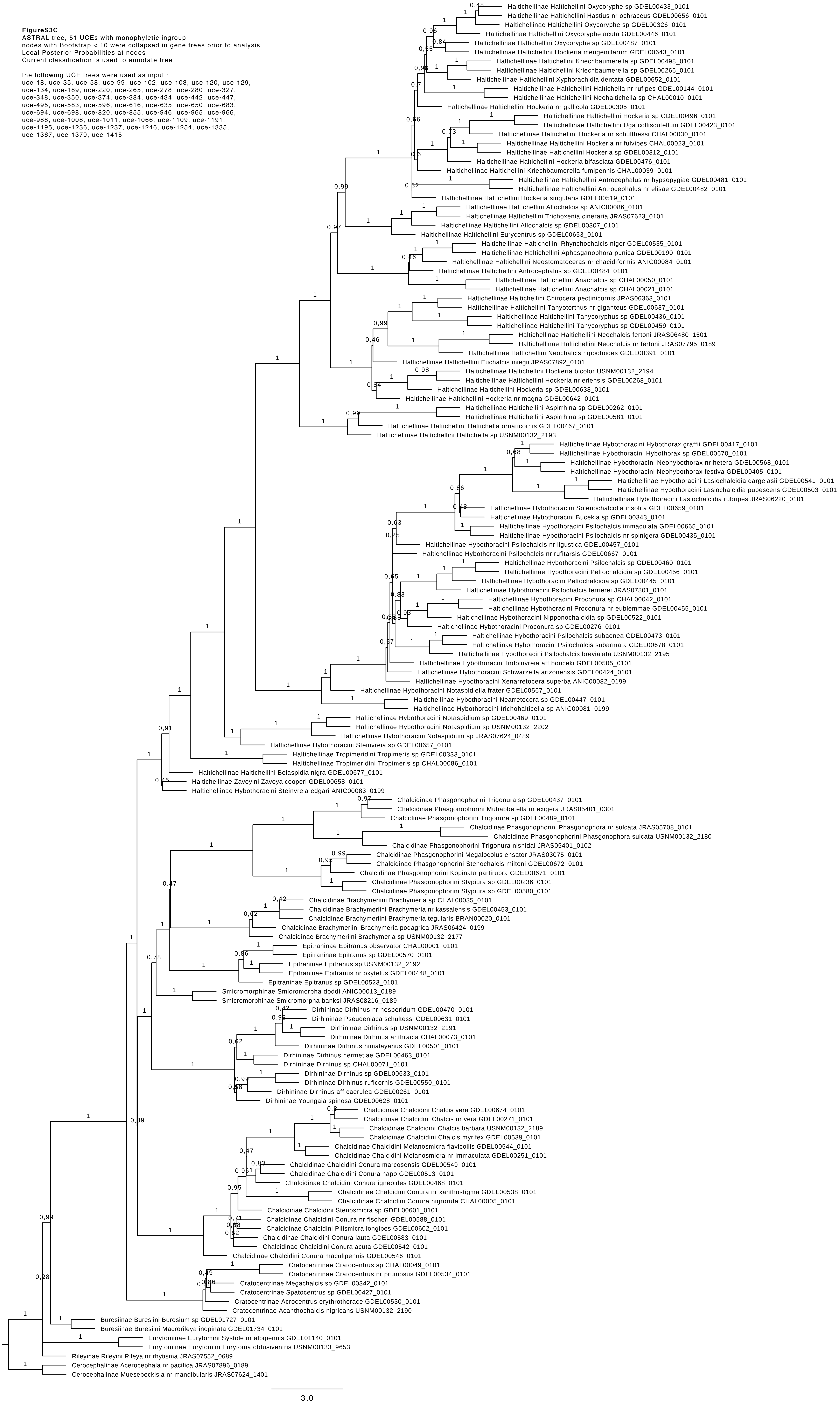
