## Supplementary material for "Ultra-Conserved Elements and morphology reciprocally illuminate conflicting phylogenetic hypotheses in Chalcididae (Hymenoptera, Chalcidoidea)": FigureS4

**Figure S4**
ASTRID tree, complete UCE data set
nodes with Bootstrap < 10 were collapsed in gene trees prior to analysis
Bootstrap support at nodes (100 replicates)
Current classification is used to annotate tree

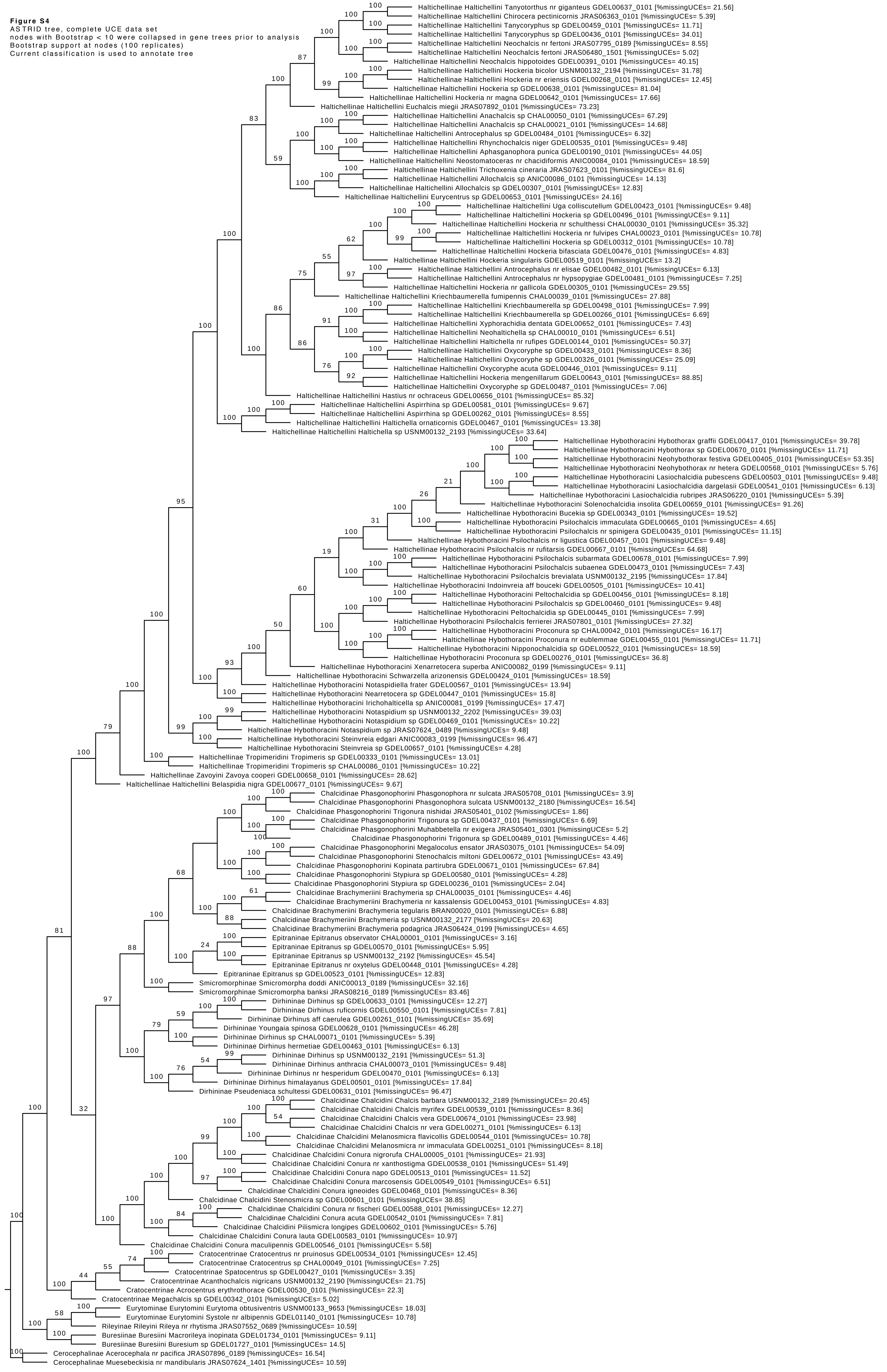
