## Supplementary figures and images for "Ultra-Conserved Elements and morphology reciprocally illuminate conflicting phylogenetic hypotheses in Chalcididae (Hymenoptera, Chalcidoidea)"

### FigureS5

**Figure S5. PAM clustering of the 538 UCE trees.**

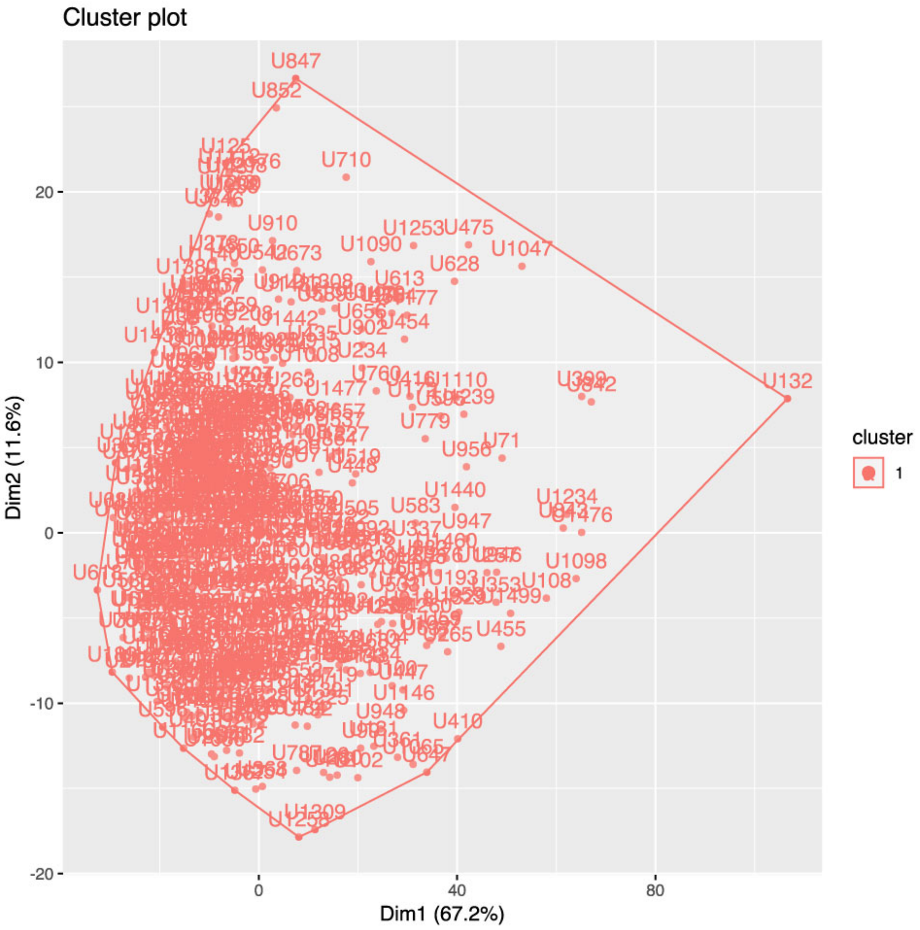
