## Supplementary material for "Ultra-Conserved Elements and morphology reciprocally illuminate conflicting phylogenetic hypotheses in Chalcididae (Hymenoptera, Chalcidoidea)": FigureS6

**Figure S6. Correlation analysis for the properties of the UCEs (Spearman rank-based correlation)**

Diagonal = distribution of each variable (GC content, Difference between the observed GC content and that predicted under the substitution model, Alpha parameter of the Gamma distribution, Parsimony informative sites content, Average bootstrap support of trees, Long Branch Score heterogeneity of trees)

Upper triangular matrix = Spearman correlation coefficients with significance level. p.value between 0 and 0.001: ***; p.value between 0.001 and 0.01: **; p.value between 0.01 and 0.05: *; p.value between 0.05 and 0.1: square

Lower triangular matrix = bivariate scatter plots

Analysis was performed with the R package PerformanceAnalytics (Peterson & Carl, 2018) with a patch to the function chart.Correlation as described in https://github.com/braverock/PerformanceAnalytics/issues/90

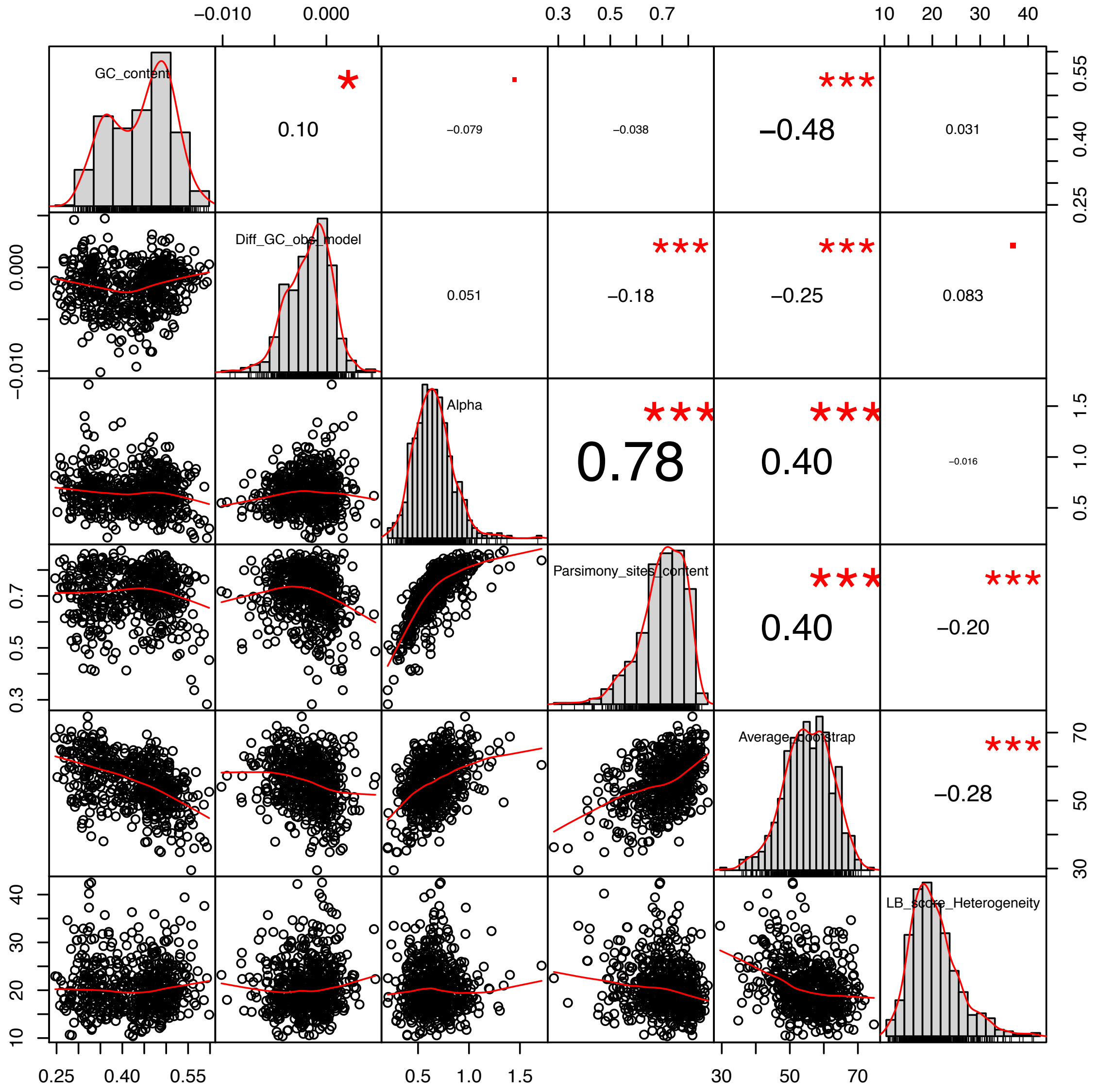


### Cited reference


Peterson BG and Carl P 2018. PerformanceAnalytics: Econometric Tools for Performance and Risk Analysis. R package version 1.5.2. https://CRAN.R-project.org/package=PerformanceAnalytics
