## Supplementary material for "Ultra-Conserved Elements and morphology reciprocally illuminate conflicting phylogenetic hypotheses in Chalcididae (Hymenoptera, Chalcidoidea)": FigureS7

**Figure S7A** **Dendrogram of samples based on GC content of UCEs (distance matrix: Gower; clustering method: Ward.D2) Current classification is used to annotate dendrogram**

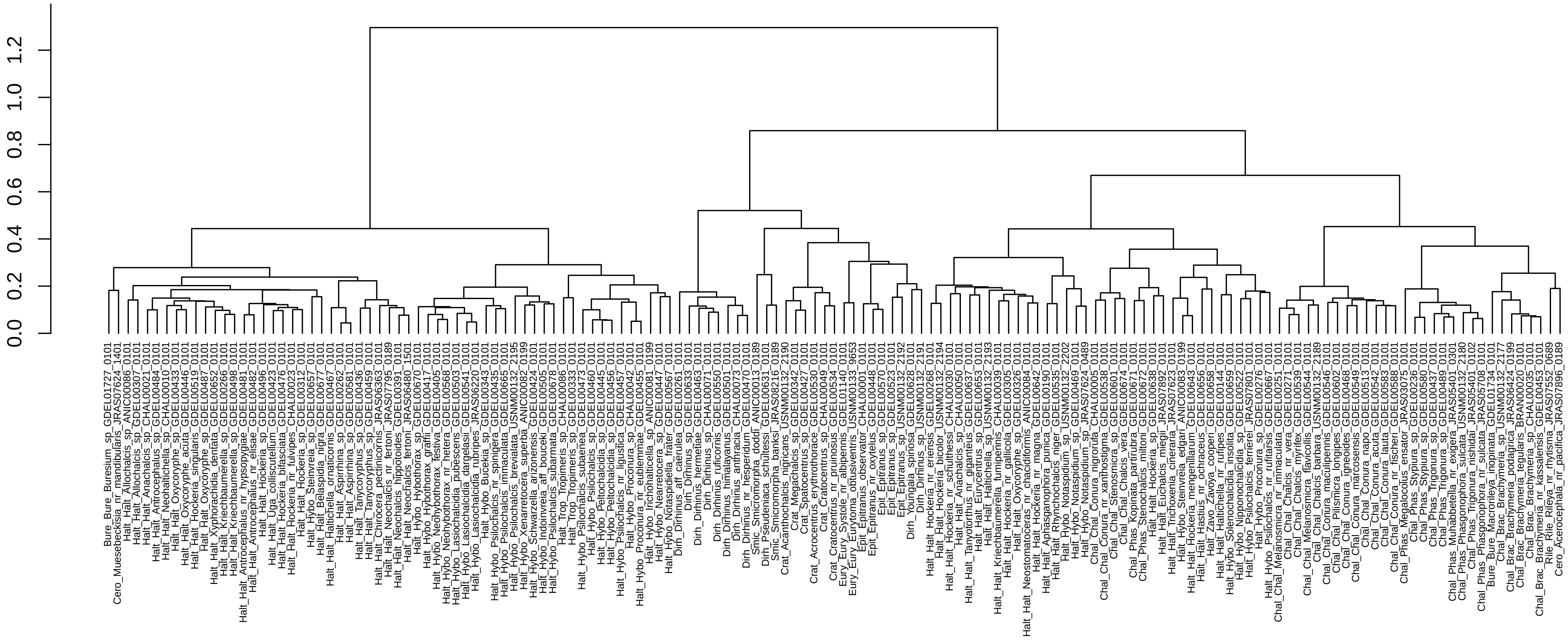

**Figure S7B** Dendrogram of samples based on LB score heterogeneity of UCEs
(distance matrix: Gower; clustering method: Ward.D2)
Current classification is used to annotate dendrogram

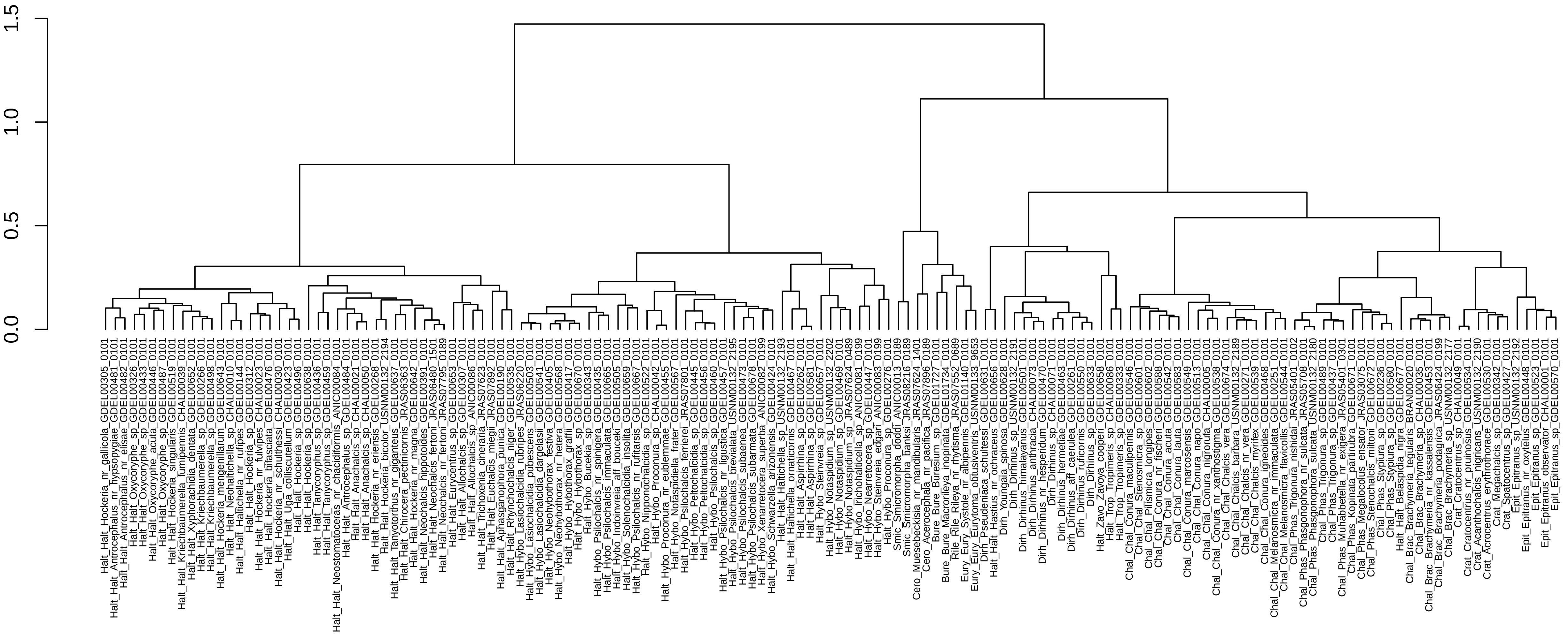
