## Supplementary material for "Ultra-Conserved Elements and morphology reciprocally illuminate conflicting phylogenetic hypotheses in Chalcididae (Hymenoptera, Chalcidoidea)": FigureS8

**Figure S8. Phylogenetic trees inferred from the morphological matrix**. A) RAxML tree (bootstrap supports at nodes – 100 replicates); B) MrBayes tree (Posterior Probabilities at nodes), nodes with PP <0.5 are collapsed

A

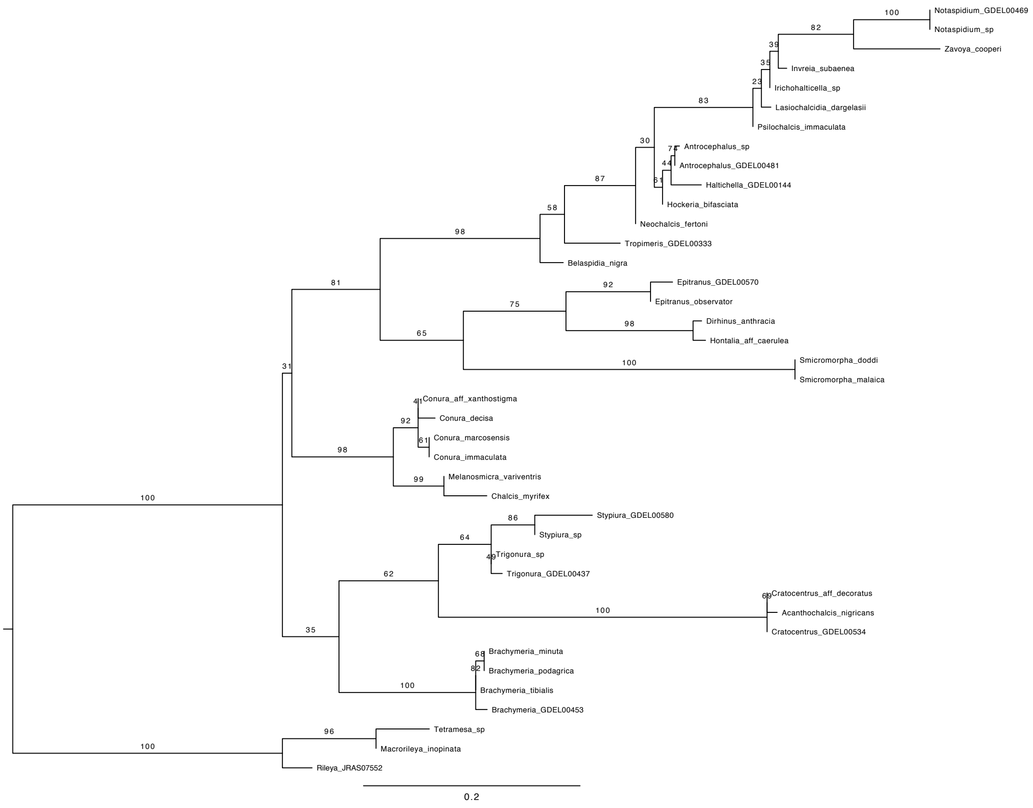


B

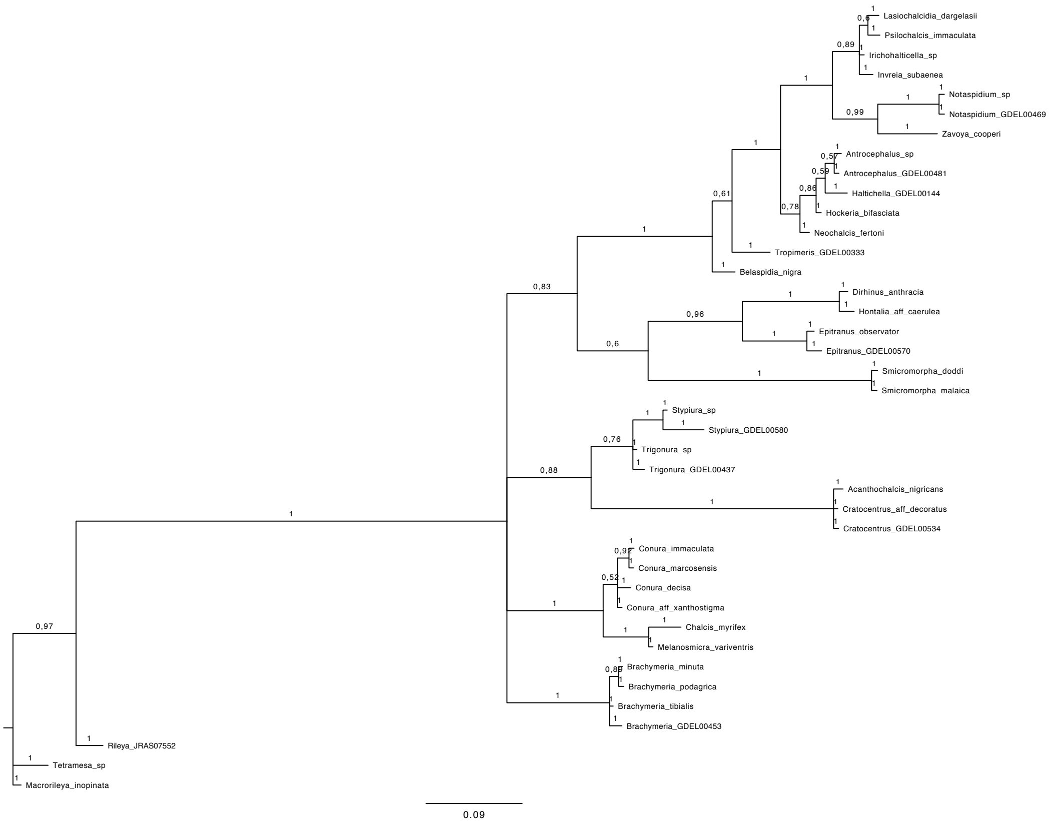
