## Supplementary material for "Ultra-Conserved Elements and morphology reciprocally illuminate conflicting phylogenetic hypotheses in Chalcididae (Hymenoptera, Chalcidoidea)": TableS2

### Table S2. UCEs in which samples were detected as outlier long branches by TreeShrink (b=20)

| Sample codes | UCEs flagged by TreeShrink (Round 1) | UCEs flagged by TreeShrink (Round 2) | Total nb of flagged UCEs |
|---|---|---|---|
| ANIC00013_0189 | uce-6, uce-38, uce-49, uce-87, uce-115, uce-244, uce-435, uce-513, uce-528, uce-595, uce-643, uce-789, uce-792, uce-898, uce-982, uce-1007, uce-1237, uce-1239, uce-1331, uce-1473 | uce-1097, uce-110, uce-1391, uce-1445, uce-1477, uce-353, uce-379, uce-384, uce-507, uce-577, uce-580, uce-597, uce-617, uce-633, uce-752, uce-913, uce-936, uce-947, uce-991 | 39 |
| ANIC00081_0199 | uce-295, uce-881, uce-1037, uce-1074 | | 4 |
| ANIC00082_0199 | uce-639 | | 1 |
| ANIC00083_0199 | uce-848 | | 1 |
| ANIC00086_0101 | uce-554, uce-1463 | | 2 |
| BRAN00020_0101 | uce-1246, uce-1279 | | 2 |
| CHAL00001_0101 | uce-848 | | 1 |
| CHAL00005_0101 | uce-888 | | 1 |
| CHAL00010_0101 | uce-714 | | 1 |
| CHAL00021_0101 | uce-597, uce-745 | | 2 |
| CHAL00030_0101 | uce-1112 | | 1 |
| CHAL00035_0101 | uce-1246 | | 1 |
| CHAL00039_0101 | uce-617, uce-846 | | 2 |
| CHAL00042_0101 | uce-125, uce-379, uce-1415 | | 3 |
| CHAL00050_0101 | uce-617 | | 1 |
| CHAL00071_0101 | uce-1214, uce-1380 | | 2 |
| CHAL00073_0101 | uce-1214, uce-1380 | | 2 |
| CHAL00086_0101 | uce-566, uce-1112, uce-1140, uce-1445 | | 4 |
| GDEL00144_0101 | uce-582, uce-1510 | | 2 |
| GDEL00190_0101 | uce-110, uce-198, uce-205, uce-792, uce-1090 | uce-898 | 6 |
| GDEL00261_0101 | uce-330, uce-877, uce-1214, uce-1380 | | 4 |
| GDEL00266_0101 | uce-49 | | 1 |
| GDEL00268_0101 | uce-632 | | 1 |
| GDEL00271_0101 | uce-1246 | | 1 |
| GDEL00276_0101 | uce-222, uce-361, uce-1237 | | 3 |
| GDEL00305_0101 | uce-566 | | 1 |
| GDEL00312_0101 | uce-582, uce-1194 | | 2 |
| GDEL00326_0101 | uce-343, uce-566 | | 2 |
| GDEL00333_0101 | uce-566, uce-597, uce-618 | | 3 |
| GDEL00342_0101 | uce-676 | | 1 |
| GDEL00343_0101 | uce-597 | | 1 |
| GDEL00391_0101 | uce-597 | | 1 |
| GDEL00417_0101 | uce-684 | | 1 |
| GDEL00423_0101 | uce-542, uce-582, uce-888 | | 3 |
| GDEL00424_0101 | uce-840, uce-969 | | 2 |
| GDEL00433_0101 | uce-343 | | 1 |
| GDEL00436_0101 | uce-163, uce-379, uce-881 | | 3 |

| | | | |
|---|---|---|---|
| GDEL00437_0101 | uce-1213 | | 1 |
| GDEL00445_0101 | uce-384 | | 1 |
| GDEL00446_0101 | uce-343 | | 1 |
| GDEL00447_0101 | uce-30, uce-597, uce-969, uce-1074 | | 4 |
| GDEL00448_0101 | uce-848 | | 1 |
| GDEL00453_0101 | uce-711, uce-1246 | | 2 |
| GDEL00455_0101 | uce-125, uce-253, uce-374, uce-1415 | | 4 |
| GDEL00456_0101 | uce-384 | | 1 |
| GDEL00457_0101 | uce-212 | | 1 |
| GDEL00459_0101 | uce-840 | | 1 |
| GDEL00460_0101 | uce-384, uce-879 | | 2 |
| GDEL00463_0101 | uce-1047, uce-1214, uce-1380 | | 3 |
| GDEL00467_0101 | uce-676 | | 1 |
| GDEL00468_0101 | uce-676 | | 1 |
| GDEL00469_0101 | uce-1055, uce-1246 | | 2 |
| GDEL00470_0101 | uce-1214, uce-1380 | | 2 |
| GDEL00481_0101 | uce-888 | | 1 |
| GDEL00487_0101 | uce-198 | | 1 |
| GDEL00496_0101 | uce-888, uce-1112, uce-1194 | | 3 |
| GDEL00498_0101 | uce-49, uce-1374 | | 2 |
| GDEL00501_0101 | uce-1214, uce-1380 | | 2 |
| GDEL00505_0101 | uce-840 | | 1 |
| GDEL00513_0101 | uce-110, uce-966 | | 2 |
| GDEL00519_0101 | uce-1208 | | 1 |
| GDEL00522_0101 | uce-125, uce-133, uce-253, uce-1415 | | 4 |
| GDEL00523_0101 | uce-848 | | 1 |
| GDEL00530_0101 | uce-925 | | 1 |
| GDEL00538_0101 | uce-645 | | 1 |
| GDEL00539_0101 | uce-1246 | | 1 |
| GDEL00542_0101 | uce-194, uce-1174 | uce-588 | 3 |
| GDEL00550_0101 | uce-1214, uce-1380 | | 2 |
| GDEL00567_0101 | uce-188, uce-270, uce-521, uce-537, uce-554, uce-633, uce-1074 | | 7 |
| GDEL00568_0101 | uce-597 | | 1 |
| GDEL00570_0101 | uce-848 | | 1 |
| GDEL00583_0101 | uce-684 | | 1 |
| GDEL00601_0101 | uce-434, uce-1159, uce-1174, uce-1239, uce-1327 | | 5 |
| GDEL00628_0101 | uce-102, uce-588, uce-662, uce-842, uce-1214, uce-1259, uce-1391 | | 7 |
| GDEL00631_0101 | uce-656 | | 1 |
| GDEL00633_0101 | uce-1214, uce-1380 | | 2 |
| GDEL00637_0101 | uce-633 | | 1 |
| GDEL00643_0101 | uce-848, uce-1446 | | 2 |

| | | | |
|---|---|---|---|
| GDEL00656_0101 | uce-343, uce-454, uce-1099, uce-1234 | uce-369, uce-594, uce-848 | 7 |
| GDEL00657_0101 | uce-1055 | | 1 |
| GDEL00658_0101 | uce-47, uce-49, uce-346, uce-527, uce-656, uce-686, uce-702, uce-745, uce-881, uce-930, uce-1097 | uce-676 | 12 |
| GDEL00659_0101 | uce-991 | | 1 |
| GDEL00667_0101 | uce-384 | | 1 |
| GDEL00670_0101 | uce-513 | | 1 |
| GDEL00671_0101 | uce-1140 | | 1 |
| GDEL00672_0101 | uce-676 | | 1 |
| GDEL01140_0101 | uce-35, uce-50, uce-379, uce-614, uce-618, uce-1246, uce-1327, uce-1379, uce-1391, uce-1446 | | 10 |
| GDEL01727_0101 | uce-35, uce-189, uce-617, uce-644, uce-684, uce-840, uce-881, uce-888, uce-1092, uce-1191, uce-1246 | | 11 |
| GDEL01734_0101 | uce-35, uce-295, uce-969, uce-1191, uce-1246, uce-1298, uce-1379 | | 7 |
| JRAS03075_0101 | uce-125 | | 1 |
| JRAS05401_0102 | uce-1213 | | 1 |
| JRAS05708_0101 | uce-1213 | | 1 |
| JRAS06424_0199 | uce-1246 | | 1 |
| JRAS07552_0689 | uce-35, uce-38, uce-95, uce-126, uce-134, uce-278, uce-340, uce-384, uce-432, uce-438, uce-764, uce-899, uce-969, uce-975, uce-1086, uce-1379 | | 16 |
| JRAS07624_0489 | uce-563, uce-1055, uce-1191 | | 3 |
| JRAS07624_1401 | uce-120, uce-134, uce-154, uce-270, uce-271, uce-340, uce-389, uce-577, uce-597, uce-617, uce-650, uce-672, uce-686, uce-881, uce-923, uce-975, uce-1050, uce-1159, uce-1171, uce-1378, uce-1379, uce-1446 | | 22 |
| JRAS07892_0101 | uce-60, uce-645, uce-848 | | 3 |
| JRAS07896_0189 | uce-95, uce-120, uce-134, uce-154, uce-271, uce-340, uce-434, uce-438, uce-470, uce-485, uce-563, uce-588, uce-599, uce-617, uce-650, uce-672, uce-686, uce-923, uce-975, uce-1066, uce-1086, uce-1140, uce-1378, uce-1379, uce-1446 | uce-618 | 26 |
| JRAS08216_0189 | uce-6, uce-910, uce-1239, uce-1331 | uce-1249, uce-1429 | 6 |
| USNM00132_2177 | uce-1112, uce-1246 | | 2 |
| USNM00132_2180 | uce-1213 | | 1 |
| USNM00132_2191 | uce-130, uce-1214 | | 2 |
| USNM00132_2192 | uce-683, uce-1159 | | 2 |
| USNM00132_2193 | uce-125, uce-343, uce-410, uce-538, uce-617, uce-632, uce-745 | | 7 |
| USNM00132_2194 | uce-49, uce-582 | | 2 |
| USNM00132_2195 | uce-32 | | 1 |
| USNM00132_2202 | uce-1050, uce-1055, uce-1249 | | 3 |
| USNM00133_9653 | uce-35, uce-189, uce-600, uce-614, uce-618, uce-684, uce-1246, uce-1379, uce-1391, uce-1446 | | 10 |
